# Towards Principled Evaluation of Single-Cell Perturbation Prediction Models

**DOI:** 10.64898/2026.07.23.740433

**Authors:** Philipp S. L. Schäfer, Kendall A. Reid, Zach Boldyga, Ekin D. Aksu, Hugo Hakem, Julio Saez-Rodriguez

## Abstract

Single-cell perturbation experiments measure how interventions alter cellular phenotypes. However, the number of possible perturbations and biological contexts far exceeds what can be tested experimentally. Motivated by this constraint, predictive models aim to extrapolate cellular responses to unseen conditions. Despite substantial efforts in model development, benchmark studies have reached inconsistent conclusions about the capabilities of current perturbation-response models. A major challenge is that evaluation protocols vary widely across studies, making results difficult to compare. Furthermore, the lack of consensus on evaluation hampers progress because it is unclear which predictive capabilities new models should prioritize. To help build consensus on evaluation principles, we develop a taxonomy that decomposes evaluation protocols into their representation, metric, score transformation, and reporting strategies. We characterize how these choices determine which aspects of prediction quality a benchmark measures and discuss criteria for selecting and assessing protocols in relation to specific benchmarking goals. We additionally provide scPertEval, a Python package with reference implementations of selected evaluation protocols, and use it to assess protocol behavior across seven publicly available single-cell perturbation datasets. By making evaluation choices and their underlying trade-offs explicit, we aim to stimulate a community discussion about developing more comparable and task-aligned evaluation protocols.

**Graphical Abstract:** 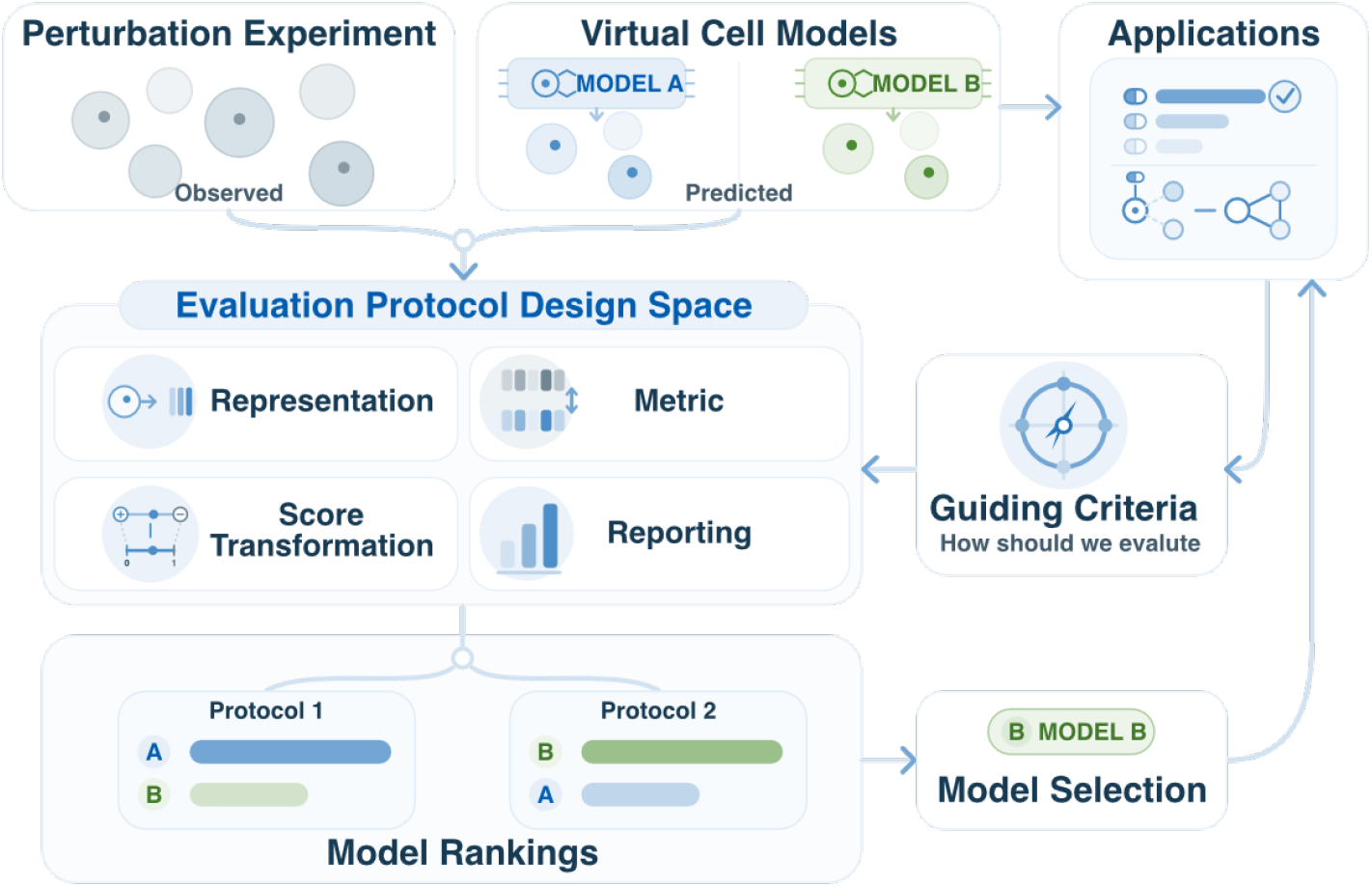

## 1 Introduction

Perturbation experiments with single-cell RNA-seq readout connect interventions to transcriptome-wide measurements in individual cells, allowing researchers to identify affected genes, altered pathways, and heterogeneous responses within the same experiment [1, 26, 42]. The scale of these assays has grown rapidly. Individual studies now profile tens of millions of cells across hundreds of small-molecule perturbations [119]. Others perform genome-wide CRISPR screens in primary cell types [122]. Nevertheless, the space of possible interventions remains far larger than what can be measured experimentally: even within a single biological context, exhaustive screening of combinatorial perturbations is infeasible. In addition, many biological systems cannot be perturbed directly for technological or ethical reasons. These limitations motivate models, including emerging “virtual cell” models [13, 74] that predict cellular responses in unseen conditions [25, 89]. The growing availability of large-scale datasets and predictive models has prompted a growing number of machine-learning competitions [30, 71, 91, 102], benchmark studies [4, 50, 55, 56, 69, 82, 105, 108, 110, 114], and evaluation approaches with corresponding software frameworks [2, 8, 37, 59, 68, 93, 104, 105, 114].

However, these efforts have not yielded a coherent view of the capabilities of perturbation-response models. Newly proposed models report substantial gains over previous methods and baselines, suggesting rapid progress, whereas benchmark studies of earlier model generations have often found no consistent advantage over simple baselines [4, 50, 105, 112]. Some of this discordance reflects differences in datasets, perturbation types, and generalization tasks, but it also reflects the wide variation in evaluation protocols across studies [93]. Even for closely related tasks, such as predicting held-out genetic perturbations in a specific cell line, studies differ in the metrics they use. These differences can substantially alter model rankings, with no single model performing best across all metrics (as illustrated using benchmarking results from Wei et al. [108]; Supplementary Figure 1). Moreover, even when studies use the same metric, they often compute it on different feature spaces. For example, one benchmark may compare predicted and observed profiles across all genes, whereas another restricts the comparison to a subset of genes. Similarly, one benchmark may compare mean normalized expression profiles, whereas another compares differential-expression statistics that summarize changes relative to control cells. As a result, performance differences between models can be difficult to attribute: they may reflect genuine differences in predictive capability, but they may also reflect differences in what the evaluation protocol rewards. The lack of consensus regarding evaluation also affects model development, since it remains unclear which predictive capabilities new models should be designed to improve. Progress in this field therefore requires greater consensus on how perturbation-response models should be evaluated.

To structure this discussion, we propose a taxonomy of evaluation protocols (Figure 1) and outline criteria for assessing the resulting protocols themselves (Table 1). The taxonomy organizes evaluation design into the representation in which predicted and observed responses are compared, the metric^1^ used to compare them, strategies such as retrieval and effect-size evaluation that compare predictions across many perturbations, and the score transformations and reporting strategies used to summarize model performance. To help researchers experiment with evaluation design choices, we additionally provide an accompanying Python package, available at https://github.com/Virtual-Cell-Research-Community/scPertEval, with reference implementations of selected evaluation protocols. Together, these resources provide a starting point for discussion by introducing a shared vocabulary for evaluation design and outlining considerations that guide protocol choices.

**Figure 1:**
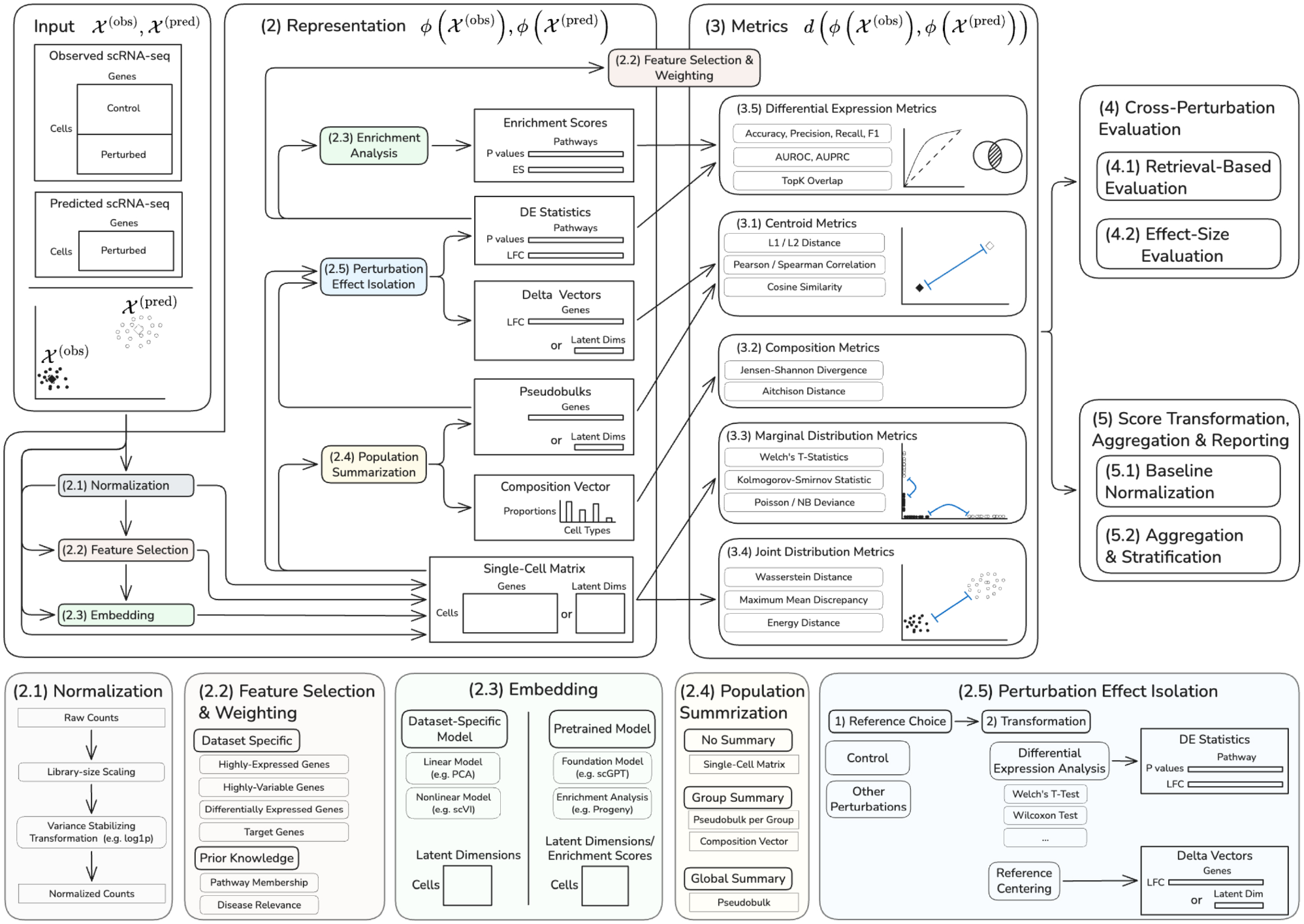
Taxonomy of evaluation protocols for single-cell perturbation prediction models. Observed single-cell perturbation data *X*^(obs)^ and predicted data *X*^(pred)^ are transformed by a benchmark-specific representation map *ϕ* before comparison. Metrics then compare the predicted and observed representations, denoted as *d*(*ϕX*(^(obs)^), *ϕX*(^(pred)^)). Beyond direct comparison of individual perturbation responses, cross-perturbation evaluation asks whether predictions rank perturbations appropriately or recover relative perturbation effect sizes. Score transformation and reporting determine how raw metric values are normalized relative to reference predictors and how scores are aggregated or stratified across perturbations, contexts, or metrics. Rounded boxes denote operations or design choices; rectangular boxes denote data representations. Numbers in parentheses refer to the corresponding sections and subsections of the manuscript. The figure uses two simplifications: pseudobulking is shown after normalization, although it is often performed on counts before normalization, and predicted responses are illustrated as single-cell profiles, although models may instead output aggregate profiles, differential-expression summaries, or other representations of the perturbation response. Abbreviations: DE, differential expression; ES, enrichment score; LFC, log fold change.

**Table 1:** Axes for assessing evaluation protocols.

| Axis | Guiding question |
| --- | --- |
| Benchmark Validity | Does the benchmark withhold the information required by the intended generalization task? |
| Task Alignment & Interpretability | Does the evaluation protocol measure the aspect of prediction quality that matters for the intended use case, and can its values be meaningfully interpreted? |
| Signal Recovery | Is the evaluation protocol sensitive to the perturbation-response signal it is intended to measure? |
| Complementarity | Do the components of the evaluation protocol capture distinct aspects of prediction quality rather than redundant ones? |
| Scalability & Estimation Error | Can the evaluation protocol be computed efficiently across large datasets, and estimated reliably from finite, high-dimensional samples? |

## 2 Representation

In perturbation-response benchmarks, the observed response for a given intervention is a population of single-cell expression profiles. For perturbation *a*, we write 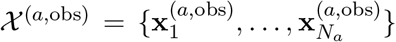, where each cell is represented by a high-dimensional gene expression vector 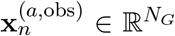. The cell-level representation *X*^(*a*,obs)^ is directly comparable to model predictions only when the model predicts a population of single-cell profiles *X*^(*a*,pred)^ [35, 51, 76]. Many models instead generate a summary representation, such as an average response [59, 90, 109], or signed differential-expression labels [43, 79, 101, 113, 115]. In such cases, the observed data, the model predictions, or both must be transformed into a shared representation before metric calculation. We formalize this choice through representation maps *ϕ*_obs_ and *ϕ*_pred_, which transform the respective available data for a perturbation into a shared format that is then compared by the evaluation metric. Thus, evaluation is performed on *ϕ*_obs_(*X* ^(*a*,obs)^) and *ϕ*_pred_(*X* ^(*a*,pred)^). For notational simplicity, we write both maps as *ϕ* when there is no ambiguity.

Representation choice, however, does more than make observed and predicted output shapes compatible. It determines which aspects of the perturbation response are emphasized, which metric families are applicable (Figure 1), and how the resulting metric values should be interpreted. In our taxonomy, we distinguish five major components of the representation map: normalization determines the expression scale (Section 2.1); feature selection restricts the considered genes (Section 2.2); embeddings and enrichment-derived representations define a new feature space (Section 2.3); population summarization determines whether evaluation retains single-cell distributions or collapses them into pseudobulks (Section 2.4); and perturbation effect isolation shifts the comparison from absolute expression profiles to perturbation-specific effects (Section 2.5).

### 2.1 Normalization

UMI counts from scRNA-seq are sparse, high-dimensional count data whose scale depends on both biological expression and technical sampling processes. In particular, observed counts depend on the total number of UMIs captured in each cell, commonly referred to as library size. Differences in library size are usually treated as a technical source of variation. Moreover, gene expression counts exhibit mean-dependent variance, also called heteroskedasticity, such that genes with higher mean expression tend to have higher variance (Supplementary Section C.1). Evaluation protocols that do not account for this mean-dependent variance can therefore overweight highly expressed genes.

To reduce library-size effects and attenuate the mean–variance relationship, evaluation is commonly performed after library-size normalization and variance-stabilizing transformation (Supplementary Section C.2), for example, by scaling cells to a common library size followed by the shifted log transformation log(1 + *x*) [3]. Alternatively, evaluation can be performed in count space using metrics designed for count data, such as Poisson or negative-binomial likelihood-based scores. These metrics include an offset term for library size and explicitly account for mean-dependent variance (Supplementary Sections D.3.1 and D.3.2).

### 2.2 Feature Selection & Weighting

Genes differ in measurement noise, biological relevance, and responsiveness to the perturbation. In particular, for many genetic perturbations, the induced response is often concentrated in a small subset of the transcriptome [69]. Metrics computed over all genes or features can therefore be dominated by weakly informative features. Evaluation protocols can mitigate this by restricting the comparison to selected features or by assigning feature-specific weights, which can be based on expression level, variability, differential expression, or prior biological knowledge.

Feature selection can be applied at different points in the evaluation pipeline. It may be an early preprocessing choice, for example when a selected gene panel is used to construct an embedding or to run differential-expression analysis. Alternatively, it may be a late evaluation choice, for example when a metric is computed only on the top-*K* most differentially expressed genes.

#### 2.2.1 Highly-Expressed Genes

Lowly expressed genes have higher relative sampling noise in scRNA-seq data. A simple feature-selection strategy is therefore to restrict evaluation to highly expressed genes, which are typically less affected by this noise. For example, Ahlmann-Eltze, Huber, and Anders [4] ranked genes by expression in the control condition and computed L2 prediction error on the top-ranked genes across several cutoffs. A related expression-based stratification was used by Li et al. [55], who partitioned non-differentially expressed genes by control expression level into highly expressed and lowly expressed genes, and evaluated these subsets separately using mean absolute error.

The main caveat is that expression level is not equivalent to biological relevance. Regulatory genes, such as transcription factors, can be lowly expressed [103] yet central to the perturbation response. High-expression filters may therefore improve robustness at the cost of excluding low-abundance but biologically important features.

#### 2.2.2 Highly-Variable Genes

In standard scRNA-seq analysis, highly variable gene (HVG) selection is widely used before downstream analyses to focus on genes that capture most biological variation [118] (Supplementary Section C.3). Analogously, many single-cell perturbation benchmarks adopt this strategy [2, 14, 27, 30, 62, 69, 90]. However, the resulting gene panels are not necessarily comparable across studies because studies may use different algorithms to identify HVGs.

Restricting evaluation to HVGs can have several caveats in perturbation settings. HVGs are commonly defined based on the variance across all cells in a dataset. For a fixed effect size, this favors genes affected across many perturbations over genes affected only in a small subset, potentially excluding perturbation-specific response genes. Moreover, because cell-cycle state, shared stress-response programs, and other systematic factors can explain a substantial fraction of the variation in single-cell perturbation data [68, 104], HVG selection may preferentially retain genes associated with these dominant axes of variation.

#### 2.2.3 Differentially Expressed Genes

A model that predicts no change or the average training response may obtain a favorable average error while missing the smaller set of genes that carry the perturbation-specific response [69]. A common strategy is therefore to use differential-expression statistics (see Section 2.5.2 and Supplementary Section C.4) for feature selection or weighting, so that the evaluation focuses on genes with evidence of perturbation-induced change [2, 62, 69, 90, 104, 106]. Selecting DEGs separately for each perturbation means that each perturbation is evaluated in a different feature space, complicating comparisons of scores across perturbations. A shared DEG set, such as the union of DEGs across perturbations [59], restores a common feature space at the cost of including genes that may not be responsive for the perturbation being evaluated. Such union sets may overlap substantially with HVG panels because both favor genes that vary across conditions; however, DEG unions will also select genes that respond only in a small subset of perturbations.

The resulting subset or weighting scheme depends mainly on two choices. The first is the reference contrast used to define perturbation-responsive genes. Most benchmarks either compare each perturbed population to control cells [2, 62, 90, 106] or to all other perturbed cells, which emphasizes genes that distinguish one perturbed state from another [69, 104]. The second is how gene-level evidence is estimated and converted into a retained gene set or weight. This includes the statistical test or model, any covariates included in the differential-expression analysis, and the quantity used to rank genes, such as a *t*-statistic, an adjusted *p*-value, or an effect size estimate. Across benchmarks, these gene-level quantities have been combined with top-*K* cutoffs [62, 90, 104] or significance-defined DEG sets [2, 106]. Alternatively, one can also turn the DEG statistics into continuous weights [68]. Such DE-based weighting changes the relative contribution of genes to the metric, such that errors on genes with stronger differential-expression evidence contribute more than errors on weakly or non-responsive genes (Supplementary Sections D.7 and D.8).

While DEG-based comparisons focus evaluation on responsive genes, they inherit general caveats from the underlying DEG-calling procedure, including sensitivity to the chosen test and selection rule (Supplementary Section C.4). For metrics restricted to observed DEGs, an additional limitation is that errors on non-DE genes are ignored by construction, allowing models to score well despite inducing large spurious changes outside the evaluated gene set. This limitation can be mitigated by defining weights from both observed and predicted responses, as in the weighted cosine term of [71]. However, using predictions to define the weights reduces comparability between predictions from different models, since each is evaluated under a different weighting scheme.

#### 2.2.4 Target Genes

In CRISPRa and CRISPRi experiments, the perturbed gene is expected to exhibit a direct expression change: activation should increase its expression, whereas interference should reduce it. This direct effect is often among the strongest observed expression changes [22, 112] and can therefore have a strong influence on metrics, especially for genetic perturbations with otherwise weak transcriptional effects. Because some models [23] learn this direct response more readily than others [109], including or excluding the target gene from metric calculations can alter relative model performance and, consequently, the conclusions of a benchmark [22].

In CRISPR knockout experiments, the target transcript is not necessarily downregulated. Its abundance depends on whether the induced mutation triggers nonsense-mediated decay, which varies across transcripts and cell types [28]. This editing-dependent response is difficult to predict from perturbation label alone and is not the downstream response of interest. Since target-gene expression largely reflects perturbation efficacy and editing outcome rather than downstream regulation, it is reasonable to exclude the perturbed gene when evaluating model predictions [88, 91]. For CRISPRa and CRISPRi, an alternative is to condition models on the measured post-perturbation expression of the target gene [49].

### 2.3 Embedding

Directly comparing perturbation responses in gene expression space can be difficult because single-cell profiles are high-dimensional, sparse, and noisy. These challenges are especially pronounced for joint distribution metrics such as maximum mean discrepancy or Wasserstein distances (Section 3.3), which compare entire cell populations rather than only average expression profiles. Embeddings address this by mapping each cell to a lower-dimensional representation before metric calculation. This improves computational tractability and may reduce the influence of technical noise by focusing the comparison on biologically meaningful axes of variation.

A useful distinction is between dataset-specific embeddings and fixed pretrained embeddings. Dataset-specific embeddings are fitted directly to the observed cells in the benchmark dataset, for example using principal component analysis (PCA) [51] or variational autoencoders such as scVI [61]. In contrast, fixed pretrained embeddings use an external model whose parameters are not fitted to the benchmark dataset [23]. This mirrors the logic of image-generation metrics such as the Fréchet Inception Distance (FID) [38] or Kernel Inception Distance (KID) [10], where images are first embedded with a pretrained Inception network and then compared. In single-cell transcriptomics, Rizvi et al. [87] adapted this idea by defining a single-cell Fréchet Inception Distance (scFID): cells are first embedded with scGPT [23], and predicted and observed populations are then compared using the Fréchet distance between Gaussian summaries in this embedding space (Supplementary Section D.4.5). However, pretrained embeddings are only useful for evaluation if they preserve perturbation-relevant variation. This remains a concern for current single-cell foundation models, which are mostly trained on observational scRNA-seq data and have not consistently outperformed simpler embeddings such as PCA or scVI [7, 22, 110].

A third class of embeddings is derived from prior biological knowledge. Instead of mapping cells to abstract latent coordinates, enrichment analysis transforms expression profiles into activity scores for predefined biological programs, such as signaling pathways or transcription-factor regulons [95]. These representations denoise expression profiles by aggregating weak signals across biologically related genes [41], and make the evaluated space more interpretable than generic latent embeddings or the original gene space. For example, X-Pert computes MSigDB Hallmark [57] gene-set scores from predicted and observed single-cell profiles for pathway-level evaluation [54], whereas scBIG uses Reactome [83] pathway memberships to transform predicted and observed expression profiles into pathway-activation representations [92]. However, like any prior-knowledge-based analysis, these embeddings inherit the limitations of their underlying knowledge source (Supplementary Section C.5).

### 2.4 Population Summarization

Observed and predicted single-cell profiles for a given condition can be collapsed into aggregate expression profiles, most commonly by summing or averaging expression across cells for each gene to obtain pseudobulk vectors [20]. We denote the observed and predicted pseudobulk profiles for perturbation *a* by ***y***^(*a*,obs)^ and ***y***^(*a*,pred)^, respectively. These vectors can then be compared using centroid-based metrics such as L2 distance or Pearson correlation (Section 3.1). This representation is simple, computationally efficient, and compatible with models that predict only average expression responses. However, pseudobulk comparisons capture only differences in the average response and therefore discard within-perturbation heterogeneity. By retaining single-cell profiles, marginal distribution metrics (Section 3.2) can assess differences beyond the average response, while joint distribution metrics can additionally capture differences in gene–gene dependence patterns (Section 3.3).

Between the two extremes, populations can be summarized within predefined cell groups, such as cell types or cell states. In this case, the representation consists of group proportions together with group-specific pseudobulks. Evaluation can then separately assess whether the model predicts the correct composition of cell groups (Section 3.4) and whether it predicts the correct expression response within each group. However, cell-type or cell-state labels may poorly approximate continuous trajectories, while rare groups can yield unstable pseudobulks and noisy proportion estimates. Local-neighborhood summaries provide an alternative when cellular states are continuous and thus poorly represented by discrete groups. For example, lochNESS measures whether perturbed cells are enriched or depleted in the neighborhood of a query cell relative to their background prevalence in the dataset [40]. Su et al. [100] used this score to evaluate predictions of perturbation-induced changes in cellular composition.

The appropriate level of summarization depends on the experimental system. In relatively homogeneous settings, such as many cancer cell line perturbation screens, one pseudobulk per perturbation may capture much of the response of interest. However, even in cell-line systems, a single perturbation can induce heterogeneous cellular responses. For example, Adamson et al. [1] reported a bifurcated unfolded-protein-response program within HSPA5-perturbed cells, showing that one perturbation can give rise to two distinct single-cell states. This limitation becomes even more important in complex systems such as organoids, immune co-cultures, or *in vivo* screens, where perturbations may affect both the abundance of cell types or states and their state-specific transcriptional programs [30, 100]. In such settings, evaluating only one pseudobulk per perturbation can obscure biologically relevant structure.

### 2.5 Perturbation Effect Isolation

The observed perturbation response *X*^(*a*,obs)^ reflects not only the biological response to the intended intervention, but also pre-existing biological variation, such as cell-type-specific expression programs, delivery-associated signals, such as viral infection or stress responses, and unwanted technical variation, such as batch effects. Isolating the perturbation effect matters because evaluation should reward models for capturing intervention-specific changes rather than for reproducing cell-type-specific expression, batch structure, delivery-associated signal, or other variation that covaries with the perturbation label. This isolation is challenging because scRNA-seq assays usually provide unpaired measurements: the same cells cannot be observed before and after perturbation. Perturbation effect isolation aims to estimate the effect of an intervention by comparison to a reference population, such as control cells. This is commonly done in two related ways: by subtracting a reference pseudobulk or by estimating differential-expression statistics against a reference population. For evaluation, the same transformation is applied to both observed and predicted responses, unless a model directly predicts the resulting perturbation-effect representation.

#### 2.5.1 Reference Centering

Reference centering defines perturbation effects by subtracting a reference pseudobulk **y**^(ref)^ from each observed and predicted perturbation pseudobulk, yielding delta vectors Δ^(*a*,obs)^ = **y**^(*a*,obs)^ − **y**^(ref)^ and Δ^(*a*,pred)^ = **y**^(*a*,pred)^ − **y**^(ref)^. When **y** is represented on a normalized log-expression scale, these delta vectors are commonly referred to as log fold-change (LFC) vectors. This representation is computationally efficient and directly targets average perturbation-induced changes, but discards within-perturbation cellular heterogeneity (Section 2.4).

These delta vectors are then compared using centroid metrics (Section 3.1). Correlation-based similarity metrics are particularly common for delta-vector evaluation [4, 23, 90, 109]. Pearson correlation and cosine similarity assess whether the model recovers the pattern of perturbation-induced gene expression changes, such as whether genes that are up- or down-regulated in the observed response are also predicted to change in the same direction. In contrast, correlations computed on absolute expression pseudobulks can be misleadingly high when perturbations induce no or only weak transcriptional effects, because even a model that predicts no change relative to the reference can obtain high correlation values [23]. Notably, subtracting the same reference pseudobulk from the observed and predicted profiles leaves translation-invariant metrics, such as L2-type distances, unchanged, but can substantially change correlation- and cosine-based similarities.

The choice of reference determines which variation is removed and which perturbation signal is emphasized. Control cells are the most natural reference when the goal is to estimate perturbation-induced change relative to an untreated or non-targeting baseline. More specific references can also be used to reduce unwanted variation. For example, Wenkel et al. [109] use batch-matched control profiles after showing that mean control expression profiles are more similar within batches than across batches, indicating batch effects. However, more specific references are not always preferable: if few reference cells are available per batch, the reference estimate can become noisy, increasing the variance of the resulting delta vectors.

A control reference isolates perturbed cells from controls, but it does not necessarily isolate the signal that distinguishes one perturbation from another. Mejia et al. [68] and Viñas Torné et al. [104] argue that many Perturb-seq datasets contain systematic shifts shared across perturbations, such as generic stress responses. In this setting, a model can obtain high control-referenced delta correlations by predicting this shared perturbed-versus-control shift for all perturbations, without distinguishing the effect of one perturbation from another. To address this, these works consider references based on all perturbed cells in the training partition rather than the control population, shifting the evaluation toward signals that distinguish individual perturbations from other perturbations. However, this also means that the resulting scores depend on the perturbations included in the training partition. If the train–test split changes, the perturbed-cell reference changes as well, which can make scores less comparable across splits [75]. More generally, a strong shared perturbation response can also reflect dataset-specific selection bias, for example when perturbations predominantly target the same pathway [1]. Such datasets may therefore be less suitable for evaluating whether models extrapolate to distinct held-out perturbations, including perturbations of unseen pathways.

One caveat is that reference centering can itself inflate correlation-based performance estimates when the same reference estimate is used to define both the observed and predicted delta vectors [73]. Because the reference pseudobulk **y**^(ref)^ is estimated from a finite set of reference cells, and the same estimate is subtracted from both the observed and predicted responses to form Δ^(*a*,obs)^ = **y**^(*a*,obs)^ −**y**^(ref)^ and Δ^(*a*,pred)^ = **y**^(*a*,pred)^ −**y**^(ref)^, the two delta vectors share its sampling noise. For correlation- or cosine-based comparisons, this shared random component can induce spurious positive similarity, even when the prediction contains little or no perturbation-specific signal, and the bias can be substantial in high-dimensional settings with limited numbers of reference cells. Nicol, Shivakumar, and Irizarry [73] therefore propose estimating the reference from independent samples, for example by splitting the reference cells into two disjoint groups and comparing **y**^(*a*,obs)^ −**y**^(ref,1)^ with **y**^(*a*,pred)^ −**y**^(ref,2)^. Such split-reference estimators remove the upward bias, although they can be conservative and introduce additional variance, which may require averaging over multiple random splits for stability.

#### 2.5.2 Differential Expression Analysis

Like reference-centered delta vectors, differential expression analysis defines perturbation effects by contrasting a perturbed population with a reference population. However, DE analysis tests these gene-wise deltas against a null hypothesis of no change, allowing the analysis to account for sampling noise and, depending on the method, experimental covariates. This makes differential expression analysis a standard way to interpret scRNA-seq perturbation experiments biologically. It is therefore natural to evaluate whether predicted responses recover the observed differential-expression statistics, including the genes called differentially expressed and the direction and magnitude of their changes. For perturbation *a*, this can be assessed by comparing DE statistics computed from *X* ^(*a*,obs)^ versus *X* ^(ref)^ with corresponding statistics computed from *X* ^(*a*,pred)^ versus the same reference. DEG information can enter evaluation in two distinct ways. First, DE analysis can be used to select genes before applying metrics such as Pearson correlation, or to define gene-specific weights for metrics such as weighted MSE (WMSE) (Section 2.2.3). Second, the outputs of DE analysis can themselves be compared, for example by measuring overlap between observed and predicted DEG sets (Section 3.5).

Compared with simple delta-vector metrics, DE-based comparisons are computationally more expensive, since they require a separate differential-expression analysis for each perturbation– reference comparison. They are also sensitive to statistical power: significance-defined DEG sets are difficult to compare across perturbations, because perturbations with more cells can yield larger DEG sets at the same effect size simply through higher power [72]. Moreover, DEG calls can vary substantially with the choice of differential-expression test and DEG selection rule (Supplementary Figure 2). Additional caveats are described in Supplementary Section C.4.

## 3 Metrics

Metric choice and representation choice are tightly linked. The representation determines what information about the perturbation response is captured, whereas the metric determines how differences in that representation are scored. Changing either one can therefore change the meaning of the evaluation. This can be formalized as *d*(*ϕ* (*X* ^(*a*,obs)^, *ϕ X* ^(*a*,pred)^)), where *ϕ* denotes the representation map and *d* denotes the metric applied in that representation.

Different metrics applied to the same representation can reward different properties. For delta vectors, Pearson correlation emphasizes agreement in the pattern of up- and down-regulated genes, whereas L2 distance is sensitive to the absolute size of gene-wise errors and therefore penalizes predictions with incorrect effect magnitude. Conversely, the same metric can have different meanings in different representations. For example, L2 distance between pseudobulks in gene expression space compares average gene expression profiles, whereas L2 distance between pseudobulks in an embedding space compares average transformed coordinates that may encode structure beyond gene-wise means.

We group metrics into five families according to the aspect of the perturbation response they compare: centroid metrics summarize aggregate profiles; marginal and joint distribution metrics compare single-cell distributions; composition metrics compare cell-state proportions; and differential-expression metrics compare gene-wise differential-expression statistics (Figure 2).

**Figure 2:**
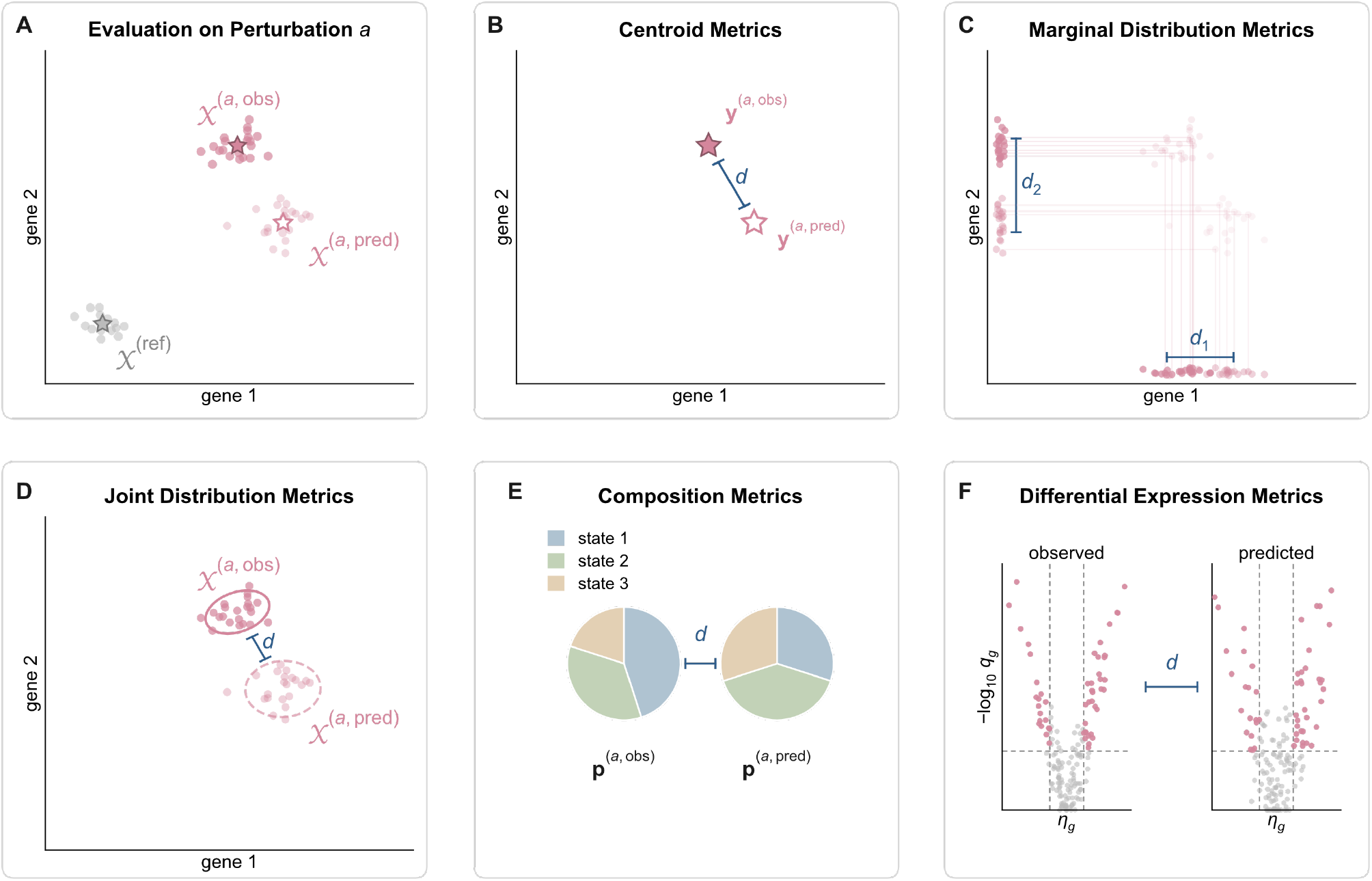
Five metric families compare different aspects of the single-cell perturbation response. The schematic shows evaluation for one perturbation *a*. In panels A–D, axes denote a toy two-gene expression space; dots denote individual cells and stars denote pseudobulks. Grey marks the reference population *X* ^*(ref)*^, *darke*r filled symbols denote observed data, and lighter symbols, open stars, or dashed outlines denote model predictions where shown. **(A)** Evaluation compares the observed population *X* ^*(a,obs)*^ *with* the predicted population *X* ^(*a*,pred)^, optionally relative to the reference. **(B)** *Centroid metrics* compare pseudobulk vectors, *d*(**y**^(*a*,obs)^, **y**^(*a*,pred)^), discarding within-population heterogeneity and gene–gene dependence. **(C)** *Marginal distribution metrics* compare the observed and predicted single-cell distributions feature by feature (*d*_1_ for gene 1, *d*_2_ for gene 2) and aggregate the per-feature scores, capturing distributional differences beyond the mean but ignoring dependence between features. **(D)** *Joint distribution metrics* compare the full joint single-cell distributions in the evaluated space; ellipses indicate covariance structure, so these metrics can also capture gene–gene dependencies. **(E)** *Composition metrics* compare cell-state proportion vectors, **p**^(*a*,obs)^ and **p**^(*a*,pred)^, separately from any state-specific expression comparison. **(F)** *Differential-expression metrics* compare gene-level summaries obtained by contrasting each observed or predicted perturbation response with a reference population, such as effect sizes *η*_*g*_ and adjusted *p*-values *q*_*g*_ shown here as volcano plots.

### 3.1 Centroid Metrics

The simplest class of metrics compares observed and predicted perturbation responses through their centroids, usually mean gene expression vectors (pseudobulks), thereby discarding within-population heterogeneity and gene–gene dependence structure (Section 2.4). Common examples include L1 distance (MAE), L2 distance (MSE, RMSE), *R*^2^, Pearson correlation, Spearman correlation, and cosine similarity (Supplementary Section D.2). As discussed in Section 2.2.3, perturbation responses are often concentrated in a small subset of genes. In such cases, many commonly used centroid metrics fail to separate good from bad predictions when the comparison includes all measured genes (Section 6.3). This can be addressed by restricting the comparison to perturbation-responsive genes or by assigning gene-specific weights. Several weighted centroid metrics have been proposed (Supplementary Section D.8), including weighted mean squared error and weighted cosine similarity [68, 69, 71, 105]. The weights are commonly derived from differential-expression statistics (Supplementary Section D.7). The main advantages of centroid metrics are their interpretability and low computational cost.

Within this family, different metrics emphasize different aspects of prediction quality. L1- and L2-type metrics are sensitive to errors in effect magnitude, whereas Pearson correlation and cosine similarity emphasize directional agreement and are insensitive to global rescaling of the evaluated vectors (Supplementary Figure 3). For reference-centered delta vectors (Section 2.5.1), this directional agreement corresponds to preserving the pattern of up- and down-regulated genes. In this setting, two predictions can have similar L2 error but differ substantially in whether they preserve the direction or sign structure of the observed perturbation effect (Supplementary Figure 4). However, correlation- and cosine-based comparisons of delta vectors become unstable when the observed perturbation effect is close to zero, because random fluctuations around the reference can dominate the estimated direction. Such similarities on delta vectors are most interpretable when the observed response has sufficient magnitude [59], or when evaluation is restricted or weighted toward genes with evidence of perturbation-induced change.

### 3.2 Marginal Distribution Metrics

Rather than reducing each perturbation to a single centroid, marginal distribution metrics compare predicted and observed cells feature by feature and then aggregate the resulting scores across features (Supplementary Section D.3). Common examples include gene-wise count likelihoods, such as Poisson or negative-binomial negative log-likelihoods, and gene-wise two-sample test statistics, such as Welch’s *t*-statistics or Kolmogorov–Smirnov statistics [37, 45]. Because these metrics compare cell-level values for each feature, they can capture differences that are lost when each perturbation is reduced to a pseudobulk centroid. Which differences they capture depends on the metric: for example, a gene-wise *t*-test mainly reflects changes in mean expression, whereas a Kolmogorov–Smirnov statistic can also respond to changes in variance, skewness or multimodality (Supplementary Figure 5). At the same time, marginal distribution metrics remain computationally and statistically simpler than joint distribution metrics because each feature is treated separately. Their main limitation is that they ignore dependence between features: two populations can have identical marginal distributions for every gene while differing in gene–gene covariance or more complex co-expression structure (Supplementary Figure 6).

### 3.3 Joint Distribution Metrics

Joint distribution metrics directly compare the predicted and observed distributions of single-cell profiles in the chosen feature or embedding space [11]. Unlike centroid metrics, which compare only average profiles, and marginal distribution metrics, which compare each feature independently, joint distribution metrics can also capture differences in covariance structure and nonlinear dependencies between features. Common examples include maximum mean discrepancy (MMD), energy distance, and Wasserstein distances (Supplementary Section D.4).

The expressiveness of joint distribution metrics comes with two practical challenges: comparing high-dimensional distributions can be noisy when few cells are available per perturbation, and many joint distribution metrics are expensive to compute because they require pairwise cell–cell comparisons or optimal-transport computations. To address these challenges, joint distribution metrics are commonly computed in lower-dimensional representations, such as PCA or scVI embeddings (Section 2.3), or approximated directly, as in sliced Wasserstein distances that average Wasserstein distances over random one-dimensional projections [52]. When joint distribution metric scores are compared across perturbations, unequal cell counts can confound interpretation because empirical distances may change with sample size. A perturbation with fewer cells may receive a different score simply because its distribution is estimated less precisely, not because the prediction is worse. Peidli et al. [78] observed such a dependence for energy distance, which varied with the number of cells per perturbation. To mitigate this effect, they subsampled each perturbation to the same number of cells before comparing scores across perturbations.

### 3.4 Composition Metrics

Composition-aware evaluation separates each perturbation response into two components: a composition vector containing the proportions of cells assigned to predefined groups, and group-specific cellular representations describing the expression profiles within each group (Section 2.4). The group-specific representations can be evaluated with centroid or distributional metrics, whereas the composition vector requires metrics suited to compositional data. Specifically, composition vectors lie on a simplex: their entries are non-negative and sum to one. As a result, changes in one group proportion necessarily affect the remaining proportions, so the entries cannot be interpreted as independent features. Composition metrics, such as total variation distance, Jensen–Shannon divergence, and Aitchison distance, are designed for such constrained vectors (Supplementary Section D.5). Several recent benchmarks have included cellular composition as one component of their evaluation [30, 51, 58, 100]. For example, the Obesity Machine Learning Competition evaluated predicted proportions of cells assigned to adipogenic programs separately from transcriptome-wide expression accuracy [30].

### 3.5 Differential Expression Metrics

Differential-expression metrics compare predicted and observed perturbation responses through gene-level summaries obtained by contrasting each perturbed population with a reference population, such as control cells (Section 2.5 and Section 2.5.2). Rather than directly comparing expression values across cells or pseudobulks, they compare the outputs of differential-expression analysis, such as adjusted *p*-values, effect-size estimates (e.g. log fold changes), and test statistics (Supplementary Section D.6, Supplementary Figure 7). Because differentially expressed genes are a central output of perturbation experiments, DE-based metrics are often directly interpretable. They evaluate whether a model recovers the gene-level changes that would typically guide biological interpretation and experimental follow-up [24, 91]. As with all DE-based evaluation, however, these metrics inherit the caveats of the underlying DE pipeline (Supplementary Section C.4).

A common strategy is to reduce DE evidence to discrete gene-level labels, such as up-regulated, down-regulated, or unchanged, and then compare these labels between observed and predicted responses. This reflects the practical role of DE analysis: genes with sufficiently strong statistical evidence are often treated as more plausible targets for interpretation, hypothesis generation, or follow-up experiments. For example, genes may be classified as differentially expressed or not based on an adjusted *p*-value threshold, potentially combined with a minimum effect-size requirement. If genes are assigned binary DEG/non-DEG labels, standard binary classification metrics such as precision, recall, and F1 score can be used (Supplementary Figure 7C). Since most genes are typically not differentially expressed for any given perturbation, these comparisons are often highly class-imbalanced; per-class or class-balanced summaries can therefore be more informative than aggregate accuracy alone [101]. If genes are instead partitioned into down-regulated, unchanged, and up-regulated, the comparison becomes a multi-class classification problem. In this setting, macro-F1 is commonly used because it gives equal weight to each class rather than being dominated by the unchanged class (Supplementary Figure 7B). A related direction-aware metric is *directionality agreement* [2, 106]. Rather than asking only whether the same genes are called differentially expressed, it asks whether overlapping DEGs are predicted to change in the same direction as in the observed response.

However, predicted DE evidence may be poorly calibrated. For example, a model may assign systematically too many or too few genes to the DEG class, even if it still ranks the most responsive genes near the top. If the goal is to evaluate this ranking while avoiding direct penalties for such calibration errors, DEG recovery can instead be treated as a ranking problem. Each gene receives a binary label indicating whether it is an observed DEG, while the predicted DE evidence provides a continuous score used to rank genes. AUROC measures how well observed DEGs are ranked ahead of non-DE genes across all possible thresholds, whereas AUPRC emphasizes the recovery of DEGs among the highest-ranked predictions (Supplementary Figure 7D) [67, 86]. Because only a small fraction of genes is typically differentially expressed, Zhu et al. [121] advocate AUPRC as a biologically relevant summary of the precision–recall trade-off. However, class imbalance alone does not inherently make AUPRC preferable to AUROC [67]. Rather, AUPRC is most appropriate when the evaluation should prioritize recovery of a limited number of candidate genes for downstream biological validation.

Whereas AUROC and AUPRC summarize ranking performance across many possible thresholds, a related strategy is to compare the top*-K* most differentially expressed genes. Here, the observed and predicted DE evidence is used to rank genes, and the metric asks how much the two top-ranked sets overlap (Supplementary Figure 7E) [2, 46, 108, 114]. The interpretation depends on how *K* is chosen. If *K* is set to the number of observed DEGs, the metric has a recall-like interpretation: it asks what fraction of observed DEGs is recovered among the top-ranked predicted genes. If *K* is set to the number of predicted DEGs, the metric has a precision-like interpretation: it asks what fraction of predicted DEGs is supported by the observed response.

One can also avoid thresholding altogether and directly compare continuous DE outputs between prediction and ground truth (Supplementary Figure 7F). Here, each gene is represented by a DE-related quantity such as a log fold change, test statistic, adjusted-*p*-value-derived score, or a combination thereof. For example, Szałata et al. [102] represent perturbation effects using − log_10_(*p*) multiplied by the sign of the log fold change and evaluate predictions using centroid metrics such as RMSE. This retains information that is lost after thresholding, but also makes the evaluation sensitive to the tails of the chosen DE statistic. In the updated living benchmark, the authors therefore clipped *p*-values at 10^−4^ so that errors were not dominated by distinctions among genes that were already highly significant [102].

Finally, DEG outputs can be compared after mapping them to prior-knowledge-derived representations (Supplementary Section C.5). This can be useful when DEG sets do not overlap strongly, but the observed and predicted responses nevertheless implicate similar biological processes. Instead of asking whether the same genes are differentially expressed, such metrics ask whether the predicted and observed DE outputs enrich similar pathways, gene sets, or regulatory programs [80, 82]. The gene-set collection should match the biological context of the dataset or process under study: broadly applicable ontologies such as Gene Ontology can capture general cellular processes, whereas specialized collections may be more informative in disease- or cell-type-specific settings [82]. The general caveats of prior-knowledge-based evaluation are discussed in Supplementary Section C.5.

## 4 Cross-Perturbation Evaluation Strategies

Cross-perturbation evaluation strategies move beyond comparisons of individual perturbation responses by considering collections of perturbations. They are particularly relevant for experimental prioritization and lab-in-the-loop workflows, where predictions are used to decide which interventions should be tested next. Beyond prioritization, these strategies can reveal mode collapse, where a model predicts similar responses across conditions, and assess whether predicted effect sizes reflect the relative strength of observed perturbation responses.

### 4.1 Retrieval-Based Evaluation

A promising use of perturbation prediction models is to prioritize candidate perturbations for follow-up experiments [39, 120], such as identifying interventions that activate or repress a specific pathway. In evaluation settings, the candidate set is typically defined as the set of all perturbations tested in a given dataset. For each candidate perturbation, the model prediction is converted into a score that quantifies how strongly the candidate is predicted to induce the desired response. Sorting perturbations by this score produces a model-derived ranking of candidates. Retrieval-based evaluation then quantifies whether perturbations that induce the desired response in the observed data appear near the top of this ranking (see Supplementary Section D.10) [8, 47, 59, 70, 104, 114].

The *retrieval-rank* metric introduced in Wu et al. [114] is a straightforward way to evaluate whether a model prioritizes the correct candidate perturbations (see Supplementary Figure 8A). For a given perturbation *a*, the observed response of *a* is treated as the desired response. The predicted responses for all perturbations in the candidate set are then ranked by their distance to this observed response, with smaller distances corresponding to better ranks. The final score is determined by the rank of the prediction for perturbation *a*. Retrieval rank therefore asks whether a model would prioritize perturbation *a* when the desired outcome is the observed response induced by *a*. Because only the matched perturbation is treated as relevant, retrieval rank can underestimate performance when several perturbations have nearly indistinguishable observed effects.

*Hit-retrieval* metrics relax the single-hit assumption by allowing multiple perturbations to count as correct if they produce the desired response (Supplementary Figure 8B). For example, if the desired response is activation of a biological pathway, enrichment analysis can be used to label observed perturbations as hits when they activate that pathway. The same scoring procedure is then applied to predicted perturbation effects, yielding a score for each candidate perturbation and hence a ranking over candidates. In Littman et al. [59], these rankings are evaluated by AUROC, whereas Bereket and Leskovec [8] consider recall under a fixed experimental budget or at an empirical false-discovery-rate cutoff.

Littman et al. [59] introduced *phenocopy retrieval*, which evaluates whether predicted effects preserve similarity relationships between perturbations (Supplementary Figure 8C). This is useful when the goal is to identify perturbations that mimic the effect of an anchor perturbation, for example the knockdown of an undruggable target. In this setting, the anchor defines the desired response, and the goal is to recover perturbations of other, potentially druggable targets that produce similar observed effects. For a given anchor perturbation *a*, the observed data are first used to define the true phenocopies of *a*, for example as the *K* perturbations whose observed responses are most similar to the observed response of *a*. The predicted data are then used to rank candidate perturbations by their predicted similarity to *a*. The predicted ranking can then be summarized by Recall@*K*, the fraction of the *K* observed phenocopies of *a* recovered among the top-*K* perturbations ranked by predicted similarity to *a*.

In observed perturbation screens, pairwise similarity matrices between perturbation responses are often compared with prior biological knowledge, such as shared pathway membership, protein complexes, or protein-protein interactions, to assess whether the screen recovers expected biological relationships [16]. The same idea could be applied to predicted perturbation responses by asking whether perturbations that are predicted to have similar effects are enriched for known biological relationships [75]. Such evaluations can test biological plausibility, but they should not be the sole criterion for model performance, especially for models that already encode prior biological knowledge, since they would reward recovery of the priors used to construct the model.

The previous retrieval formulations ask whether predictions recover the correct or biologically useful candidate perturbations. The *transposed retrieval-rank* metric of [114] instead evaluates whether model predictions collapse toward similar responses across perturbations (see Supplementary Figure 8A). Transposed retrieval rank tests this by fixing the prediction for perturbation *a* as the query, ranking all observed perturbation responses by their similarity to that prediction, and asking whether the observed response of *a* is ranked ahead of the other observed responses. This formulation is closely related to *centroid accuracy* in Systema [104] and to the *perturbation discrimination score* used in the Virtual Cell Challenge [2, 91].

The behavior of retrieval-based metrics is still not well understood in single-cell prediction bench-marking. Liu et al. [60] show that transposed-retrieval-style metrics based on L1 and L2 distances can be highly sensitive to the scale of the predicted effect vector. In particular, for large scaling factors, the ranking is increasingly determined by directional alignment rather than by agreement in effect magnitude. More broadly, Liu et al. [60] argue that perturbation magnitudes are difficult to estimate robustly, because they can depend on guide efficiency, experimental design, and measurement noise. They therefore suggest defining discrimination-based retrieval with scale-invariant similarities such as cosine similarity.

### 4.2 Effect-Size Evaluation

Commonly used evaluation protocols, such as Pearson correlation on delta vectors, compare the pattern of up- and down-regulated genes, but do not assess whether a model recovers the relative strength of perturbation effects across conditions. Effect-size evaluation addresses this complementary question by summarizing the total effect of each perturbation as a scalar magnitude and asking whether these magnitudes agree across perturbations. This distinction is relevant because different applications emphasize different aspects of the response. For biological interpretation, recovering the pattern and direction of gene-level changes may be most important. In contrast, for applications such as cell-state engineering, where the goal is to select perturbations that push cells toward a desired transcriptomic state, it also matters how far a perturbation is expected to move cells toward that state.

In Littman et al. [59], effect-size evaluation is used as part of a two-stage framework that separates response magnitude from response direction. In the first stage, observed perturbations are labeled according to whether they induce a significant total effect. For each perturbation, this effect is quantified by the Euclidean norm of the control-centered delta vector (Section 2.5.1). Significance is determined against a perturbation-specific empirical null. This null is obtained by repeatedly sampling subsets from all cells in the dataset, with each subset containing the same number of cells as observed for that perturbation, and computing the Euclidean norm of its control-centered pseudobulk. Model predictions are scored analogously using the Euclidean norm of the predicted delta vector, and performance is summarized by AUROC (see Supplementary Figure 9). This formulation is compatible with models that output only a pseudobulk response. In the second stage, they assess response direction using cosine similarity between predicted and observed delta vectors centered on the training set mean response. This second stage is restricted to perturbations with significant total effects, because for near-zero delta vectors small amounts of noise can dominate the estimated direction, yielding cosine similarities close to zero even between technical duplicates [116].

Cell-Eval, introduced in Adduri et al. [2], uses a differential-expression-based notion of perturbation magnitude. For each perturbation, the effect size is quantified by the number of significant differentially expressed genes (Section 2.5.2). Predicted and observed effect sizes are then compared across perturbations using Spearman correlation. This formulation is applicable when the model predicts single-cell populations that can be used for differential-expression testing. However, the number of significant DEGs is sensitive to the number of cells measured for each perturbation (see Supplementary Section C.4).

## 5 Score Transformation, Aggregation, and Reporting

Different combinations of representations and metrics emphasize different properties of the perturbation response. Benchmarks therefore often report several evaluation protocols, yielding scores for each protocol, model, and held-out condition. To make these results interpretable, the resulting scores are transformed, summarized, and reported in ways that address benchmark questions such as which model performs best overall and which held-out conditions are easier or harder to predict.

### 5.1 Baseline Normalization of Metric Scores

Raw metric scores are often difficult to interpret. Their scale depends on the chosen representation, metric, and perturbation effect size. For example, an MSE of 0.05 is difficult to interpret without knowing whether a simple baseline, such as predicting no change, achieves an MSE of 0.50 or 0.06.

Baseline normalization makes raw scores more interpretable across perturbations and metrics by transforming them relative to one or more reference predictions. The transformed score can quantify how much the model improves over a lower reference, such as a no-change baseline or training-mean baseline. A common example is reporting the MSE relative to the MSE of a no-change baseline [6, 59, 81, 90]. Some protocols additionally quantify how much of the gap to a positive reference is closed [105]. This positive reference approximates the performance of an experimental replicate and is often constructed as a *technical duplicate*, obtained by splitting the observed cells for a perturbation into two disjoint subsets and treating one subset as prediction and the other one as evaluation target. However, normalization between lower and positive references is only interpretable when the evaluation protocol consistently assigns better scores to positive references than to lower references (Section 6.3).

### 5.2 Aggregation & Stratification of Metric Scores

Selecting one model over another in perturbation-response benchmarks requires comparing scores across many metrics, held-out perturbations, or biological contexts. Benchmarks therefore often aggregate metric scores into summaries for model comparison. However, such summaries can hide relevant heterogeneity: a model may perform well on average while failing systematically for particular perturbation classes or generalization regimes. Stratified reporting complements aggregation by showing performance separately across biologically or experimentally relevant subsets of the test set.

#### 5.2.1 Aggregation and Stratification Across Perturbations and Contexts

The most common aggregation strategy is an unweighted mean over held-out conditions. While simple, it gives equal weight to each test instance, irrespective of biological relevance, effect size, measurement uncertainty, or difficulty. It also summarizes only average performance, even though the variability of performance across perturbations or contexts can be important. For example, a model with slightly worse mean performance but fewer severe failures may be preferable in applications where robust behavior across test cases matters. Depending on the metric, mean aggregation can also give disproportionate influence to particular perturbations. For example, Littman et al. [59] reported that RMSE per perturbation is strongly correlated with perturbation effect size, implying that large-effect perturbations have disproportionate influence.

Weighted aggregation is useful when the benchmark should emphasize particular perturbations or balance unequal groups. For example, one may group perturbations according to biological categories, such as pathways for genetic perturbations or mechanisms-of-action annotations for small molecules, and then average category-level scores so that each category contributes equally. Alternatively, weights may reflect the reliability of individual test cases, for example by accounting for the number of observed cells. A complementary strategy is baseline normalization (Section 5.1), which expresses performance relative to reference predictions, thereby placing perturbation-level scores on a more comparable scale before aggregation.

Stratified evaluation provides a complementary view by reporting performance separately across relevant subsets of the test set. In combinatorial perturbation prediction, for example, performance is commonly reported according to whether zero, one, or both single perturbations were observed during training [90]. Similarly, Wenkel et al. [109] stratified performance by the amount of prior knowledge available for different genetic perturbations and additionally showed that performance varied with perturbation effect size. More generally, useful strata depend on the experimental design and biological context. These may include perturbation families, such as pathways or drug mechanisms of action, as well as cell type, tissue, or batch. Such breakdowns can reveal systematic model failures that are hidden by a global score, thereby identifying opportunities for improvement and informing how much confidence should be placed in real-world applications.

#### 5.2.2 Aggregation Across Metrics

Because no single metric captures all relevant aspects of perturbation-response prediction, benchmarks commonly report multiple metrics, often combined into a single composite score. This introduces an additional aggregation problem: scores must be combined not only across perturbations, but also across metrics.

A common strategy is rank aggregation. Models are ranked separately under each metric, and the resulting ranks are then averaged or otherwise combined. This places heterogeneous metrics on a common scale and avoids directly averaging quantities with different units or ranges. For example, Wei et al. [108] use rank-based aggregation across multiple metrics. However, rank aggregation can exaggerate negligible differences between nearly tied models, and the results may depend strongly on the chosen aggregation rule [65].

Baseline-normalization provides an alternative to transform raw metric values onto a shared scale (Section 5.1). The Virtual Cell Challenge used this strategy by normalizing three metrics relative to a training-perturbation-mean baseline before averaging across the metrics [5, 91]. While equal weighting of different metrics is simple, it may not account for redundancy among metrics or differences in their relevance. Unequal weighting can encode benchmark priorities. For example, in single-cell integration benchmarking, the scIB metric suite assigns larger weight to biological conservation than to batch correction [63]. Similarly, a perturbation benchmark might weight retrieval performance or distributional fidelity differently depending on whether the intended application is hit prioritization or realistic data generation.

## 6 Assessing Evaluation Protocols

The preceding sections illustrate that perturbation-response evaluation has a large design space: choices about representation, metrics, score transformations, and cross-perturbation evaluation can be combined in many ways. This flexibility is necessary, since no single protocol suits every biological context or prediction task, but it raises a practical question: how should an evaluation protocol be chosen? We organize this choice along five axes: benchmark validity, task alignment & interpretability, signal recovery, complementarity, and scalability & estimation error (Table 1).

### 6.1 Benchmark Validity

A benchmark is only valid if its train–test split withholds the information required by the intended generalization task. When it does not, performance estimates become more optimistic than the task warrants, a problem commonly referred to as leakage [21, 48]. This can happen, for example, when held-out perturbations and contexts are closely related to the training data, so that the model is not actually required to generalize in the way the task intends. Test perturbations may share pathways in genetic screens or chemical scaffolds in small-molecule screens, and held-out contexts may resemble training data through related cell lines, tissues, donors, or batches [9, 77, 96, 110, 117].

Assessing leakage is particularly difficult for foundation models pretrained on large, heterogeneous corpora, especially proprietary ones, because benchmark data may have entered pretraining [21]. Even when response profiles are genuinely withheld, advance knowledge of the evaluation set can create benchmark-specific shortcuts: a model tuned for a fixed list of targets need not solve the harder task of predicting arbitrary perturbations. Competitions can mitigate this by keeping the test conditions non-public until late in the process; in the Virtual Cell Challenge, for example, the gene names of the 100 held-out perturbations used for final scoring were released one week before the submission deadline [91].

Leakage can also arise from the evaluation protocol itself. Learned embeddings are a common source of this problem. If an embedding is fitted on both training and evaluation cells and then used for model training, the model is indirectly shaped by data it will later be scored against, even if it never sees the held-out perturbations or contexts directly. CellFlow avoids this by fitting the PCA space used to train the model on the training partition alone, whereas the predictions are scored in a PCA space fitted on the full dataset [51]. A related issue arises with reference centering: when the same estimated reference is subtracted from both the observed and predicted responses, the two delta vectors share its estimation noise, which can inflate correlation- or cosine-based similarities even when the prediction carries little task-relevant signal (Section 2.5.1) [73].

### 6.2 Task Alignment & Interpretability

Different goals favor different evaluation protocols. If the goal is experimental prioritization, retrieval-based evaluation (Section 4.1), top-*K* overlap / precision@*K* (Section 3.5), and enrichment-based scores are most actionable, because they ask whether the biologically relevant perturbations appear near the top of a candidate list. If the goal is realistic data generation, joint distributional metrics (Section 3.3) that compare full single-cell populations are more relevant, and evaluation should avoid overly restrictive gene subsets. If the goal is mechanistic interpretation, DEG directionality and pathway-level agreement may matter more than overall expression error. If the goal is screen design, the calibration of uncertainty estimates may be more important, though this is rarely evaluated at present.

A score is most interpretable when it conveys how much signal a model recovers, not merely whether one prediction beats another. A bounded scale can help here, as in correlation-based metrics, but it does not by itself make a score interpretable. A Pearson correlation of 0.95 on absolute expression profiles looks near-perfect, yet it can be unimpressive, because a no-change baseline already reaches comparable values (Section 2.5). One way to make such a score interpretable is baseline normalization (Section 5.1), which provides reference points for what good and bad performance looks like. Furthermore, some evaluation protocols are inherently easier to translate into a statement about prediction quality. In the DEG setting, AUROC can be interpreted as the probability that a randomly chosen observed DEG receives a higher predicted DE score than a randomly chosen observed non-DE gene. Such interpretations make it easier to state which aspect of prediction quality is being measured. By contrast, more abstract distributional discrepancies such as MMD or Wasserstein distances are often harder to translate into a direct biological reading.

### 6.3 Signal Recovery

Signal recovery asks whether an evaluation protocol is sensitive to the perturbation-response it is meant to detect. This is not trivial in single-cell perturbation data, which are high-dimensional, noisy, and often sparse. Metric scores can be overly sensitive to technical variation and systematic shifts, or insufficiently sensitive to real perturbation-specific effects.

In practice, signal recovery cannot be assessed against an absolute ground truth. Defining the signal to be recovered—for example, deciding which of two predictions should be considered better or which perturbations should be considered similar—already requires a comparison rule. Instead, we rely on assumptions about which relationships a useful protocol should capture (Figure 3), such as separating perturbed cells from controls or preserving expected similarities between perturbations. This contrasts with other generative-modeling domains, such as image generation, where evaluation metrics can be compared against human perceptual judgments [44, 99].

**Figure 3:**
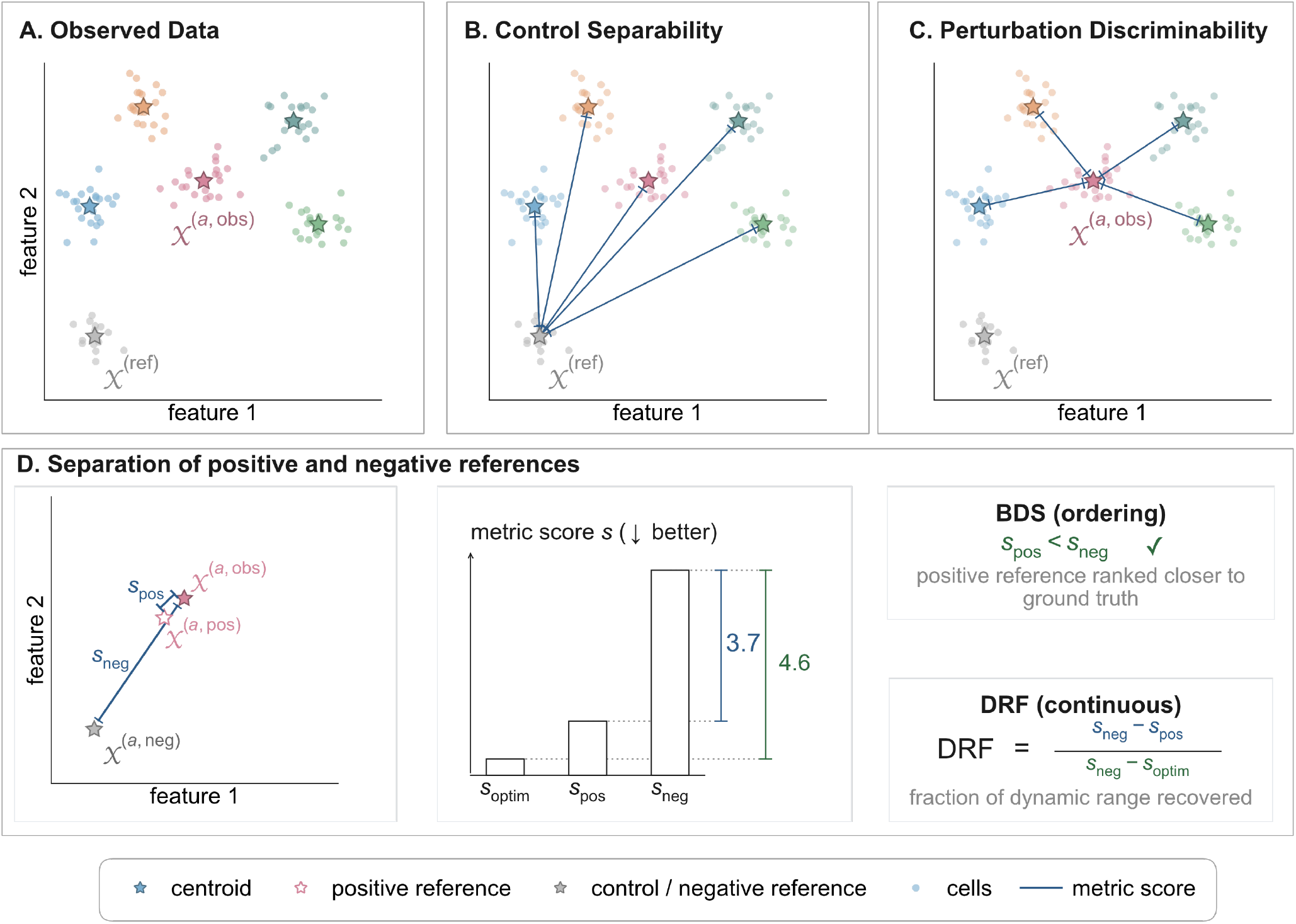
Schematic checks for signal recovery by an evaluation protocol. **(A)** Observed data contain multiple perturbation populations and a shared reference population; stars denote population centroids. **(B)** Control separability asks whether each perturbed population is distinguishable from the reference population. **(C)** Perturbation discriminability asks whether responses to different perturbations are distinguishable from one another. **(D)** Positive– negative reference separation compares a ground-truth perturbation response with a positive reference (e.g. technical duplicate) intended to preserve perturbation-specific signal and a negative, uninformative reference (e.g. control cells). The bound discrimination score (BDS) records whether the positive reference receives a better score than the negative reference. The dynamic range fraction (DRF) additionally quantifies the improvement of the positive-reference score *s*_pos_ over the negative-reference score *s*_neg_ relative to the full available improvement from *s*_neg_ to the optimal score *s*_optim_. The illustrated example gives DRF = 3.7*/*4.6 = 0.8.

#### 6.3.1 Control Separability

The simplest check is whether an evaluation protocol separates perturbed populations from controls (Figure 3B) [45, 84, 96], relying on the assumption that perturbations induce measurable deviations from the control state. Ji et al. [45] formalize this as the *control-rank percentile* (CRP), which asks whether perturbed populations are farther from the controls than control subsets are from one another. They also summarize how stable this separation is across control subsamples as a robustness criterion: stable separation indicates a reliable protocol, whereas high variability across subsamples indicates sensitivity to sampling noise or within-control heterogeneity.

#### 6.3.2 Perturbation Discriminability

Beyond separating perturbations from controls, an evaluation protocol should also distinguish perturbations from one another (Figure 3C) [31, 47, 84, 96, 102, 109]. At the most basic level, replicates of the same perturbation should be more similar to each other than to other perturbations. More broadly, related perturbations, such as compounds sharing a mechanism of action, should be more similar than unrelated ones.

In high-content imaging, Reisen et al. [84] quantified this by AUROC, treating replicate pairs as positives and randomly chosen pairs of distinct compounds as negatives, with equal numbers of each. In single-cell RNA-seq, true replicates are often unavailable because of experimental cost. In such cases, technical duplicates (Section 5.1) can serve as surrogate replicates. For example, Wenkel et al. [109] used technical duplicates to test whether pseudobulk profiles corresponding to the same perturbation retrieve one another more strongly than unrelated perturbations under a given representation and metric. Similarly, the SBB framework [105] repurposes the perturbation discrimination score (PDS; Supplementary Section D.10) as a meta-metric for signal recovery. Instead of scoring model predictions, PDS is computed only on technical duplicates from the observed data and asks whether each perturbation retrieves its matched duplicate ahead of duplicates from other perturbations. Additionally, SBB estimates statistical significance by comparing the observed self-retrieval ranks to a null distribution obtained by permuting perturbation labels across the technical-duplicate bags, thereby testing whether perturbations retrieve their matched duplicates more strongly than expected by chance.

#### 6.3.3 Separation of Positive and Negative References

Another diagnostic is whether the protocol preserves expected performance rankings (Figure 3D). Predictions expected a priori to be good, such as technical duplicates, should score better than uninformative baselines, such as predicting no change or the training mean for all perturbations. Two diagnostics formalize this idea: the *bound discrimination score* (BDS) [105] and the *dynamic range fraction* (DRF) [69]. Both compare three objects for a fixed perturbation: the ground truth, a positive reference (a technical duplicate; Section 5.1), and a negative reference. They differ in how this comparison is summarized.

BDS is the simpler diagnostic. For each perturbation it asks whether the protocol ranks the positive reference better than the negative reference, and reports the fraction of perturbations where it does. Vollenweider and Bühlmann [105] pairs this with a permutation test that flags which perturbations show a separation beyond chance (Supplementary Section D.11). DRF is the continuous analog of BDS. It measures how much of the available separation between the negative reference and the ground truth is recovered by the positive reference under the evaluation protocol (Supplementary Section D.12, Supplementary Figure 10). In the original formulation of Miller et al. [69], DRF is defined only for centroid-based metrics, with the technical duplicate as the positive reference. Because technical duplicates can be unstable when few cells are available, they also define a perturbation-specific *interpolated technical duplicate*. For each perturbation, genes with weak differential-expression evidence are shrunk toward the training mean, whereas genes with stronger perturbation-specific evidence remain closer to the corresponding technical duplicate.

To illustrate how metric and representation choices affect positive–negative reference separation, we computed the DRF for commonly used evaluation protocols across seven datasets (Table 2). Commonly used centroid metrics, such as Pearson correlation on full-gene delta vectors and MSE on full-gene pseudobulks, often achieved only limited dynamic range, indicating that these protocols do not consistently distinguish the interpolated technical duplicate from the negative reference.

**Table 2:** Mean dynamic range fraction (DRF) across evaluation-protocol configurations and datasets. Rows specify the base metric and, where applicable, reference centering, representation, positive and negative references, feature space, and weighting exponent. Columns report the mean DRF for Wessels23 [111], Replogle22 RPE1 [85], Replogle22 K562 [85], Nadig25 Jurkat [72], Nadig25 HepG2 [72], ArcH1 [91], and Kaden25 RPE1 [97]. Darker blue cells indicate larger positive mean DRF values and orange cells indicate negative values. Empty configuration fields denote choices that are not applicable to the corresponding protocol.

| Base Metric | Reference Centering | Representation | Positive Reference | Negative Reference | Feature Space | Weight Exponent | Wessels23 | Replogle22 RPE1 | Replogle22 K562 | Nadig25 Jurkat | Nadig25 Hepg2 | ArcH1 | Kaden25 RPE1 |
| --- | --- | --- | --- | --- | --- | --- | --- | --- | --- | --- | --- | --- | --- |
| Pearson | — | Pseudobulk | Interpolated TD | All-pert. (mean) | — | — | +0.33 | +0.17 | +0.11 | +0.04 | +0.08 | +0.61 | -0.20 |
| Pearson | Ctrl | Pseudobulk | Interpolated TD | All-pert. (mean) | — | — | +0.39 | +0.18 | +0.22 | +0.15 | +0.16 | +0.70 | +0.02 |
| Pearson | Pert | Pseudobulk | Interpolated TD | Control (mean) | — | — | +0.49 | +0.07 | +0.17 | +0.14 | +0.07 | +0.56 | +0.10 |
| Pearson | Pert | Pseudobulk | Interpolated TD | Control (mean) | Sig. DEGs | — | +0.74 | +0.37 | +0.26 | +0.31 | +0.15 | +0.83 | +0.42 |
| Pearson | Pert | Pseudobulk | Interpolated TD | Control (mean) | Top-50 DEGs | — | +0.24 | +0.36 | +0.30 | +0.33 | +0.26 | +0.96 | +0.14 |
| MSE | — | Pseudobulk | Interpolated TD | All-pert. (mean) | — | — | +0.33 | +0.17 | +0.12 | +0.05 | +0.08 | +0.61 | -0.19 |
| MSE | — | Pseudobulk | Interpolated TD | All-pert. (mean) | Sig. DEGs | — | +0.68 | +0.54 | +0.49 | +0.46 | +0.41 | +0.84 | +0.24 |
| MSE | — | Pseudobulk | Interpolated TD | All-pert. (mean) | Top-50 DEGs | — | +0.51 | +0.54 | +0.53 | +0.47 | +0.38 | +0.97 | +0.20 |
| WMSE | — | Pseudobulk | Interpolated TD | All-pert. (mean) | — | 1.0 | +0.66 | +0.41 | +0.31 | +0.26 | +0.27 | +0.93 | +0.02 |
| WMSE | — | Pseudobulk | Interpolated TD | All-pert. (mean) | — | 2.0 | +0.77 | +0.54 | +0.47 | +0.42 | +0.39 | +0.99 | +0.15 |
| WMSE | — | Pseudobulk | Interpolated TD | All-pert. (mean) | — | 4.0 | +0.75 | +0.58 | +0.56 | +0.49 | +0.40 | +0.99 | +0.28 |
| TransposedRank | — | Pseudobulk | Interpolated TD | All-pert. (mean) | — | — | +0.93 | +0.67 | +0.48 | +0.46 | +0.50 | +0.96 | +0.30 |
| Rank | — | Pseudobulk | Interpolated TD | All-pert. (mean) | — | — | +0.97 | +0.80 | +0.75 | +0.69 | +0.69 | +0.99 | +0.30 |
| DE Overlap @ 50 | — | Single-cell | Technical duplicate | All-pert. {8192 cells} | — | — | +0.08 | +0.11 | +0.14 | +0.10 | +0.07 | +0.51 | +0.05 |
| DE AUPRC | — | Single-cell | Technical duplicate | All-pert. {8192 cells} | — | — | +0.22 | +0.03 | +0.09 | +0.06 | +0.04 | +0.33 | +0.22 |
| DE AUROC | — | Single-cell | Technical duplicate | All-pert. {8192 cells} | — | — | +0.42 | +0.10 | +0.11 | +0.17 | +0.13 | +0.42 | +0.40 |
| MMD | — | Single-cell | Technical duplicate | All-pert. {8192 cells} | Top-50 DEGs | — | -0.11 | +0.18 | +0.44 | +0.12 | -0.30 | +0.97 | +0.38 |
| MMD | — | Single-cell | Technical duplicate | All-pert. {8192 cells} | PCA-50 | — | +0.95 | +0.89 | +0.90 | +0.83 | +0.82 | +0.98 | +0.66 |
| ED | — | Single-cell | Technical duplicate | All-pert. {8192 cells} | Top-50 DEGs | — | +0.51 | +0.61 | +0.63 | +0.49 | +0.35 | +0.98 | +0.18 |
| ED | — | Single-cell | Technical duplicate | All-pert. {8192 cells} | PCA-50 | — | +0.96 | +0.90 | +0.90 | +0.84 | +0.84 | +0.98 | +0.66 |
| W<sub>2</sub> (Sinkhorn) | — | Single-cell | Technical duplicate | All-pert. {8192 cells} | Top-50 DEGs | — | +0.32 | +0.25 | +0.16 | +0.14 | +0.16 | +0.40 | +0.05 |
| W<sub>2</sub> (Sinkhorn) | — | Single-cell | Technical duplicate | All-pert. {8192 cells} | PCA-50 | — | +0.16 | +0.17 | +0.11 | +0.07 | +0.11 | +0.11 | -0.03 |

Likewise, DEG overlap and DE AUPRC showed limited dynamic range, raising questions about the suitability of these direct differential-expression metrics as primary evaluation metrics. In contrast, retrieval-based metrics achieved some of the strongest and most consistent separation across datasets. Among joint distribution metrics, MMD and energy distance achieved substantially higher DRF scores in a 50-dimensional PCA space than when computed directly on the top 50 DEGs, whereas the Wasserstein-2 distance remained comparatively weak in both representations. For centroid metrics, restricting the comparison to significant or top-ranked DEGs, or weighting genes according to their differential-expression evidence (Section 2.2.3), generally increased the DRF. However, this improvement also exposes a limitation of the DRF: protocols can obtain high values by concentrating on a small set of strongly responding genes, while errors on weakly responsive or non-responsive genes contribute little to the score. The DRF is therefore useful for diagnosing whether an evaluation protocol separates positive from negative references, but should not serve as the sole criterion for protocol selection.

#### 6.3.4 Sensitivity to Graded Perturbation Strength

Ji et al. [45] introduced *biological reproducibility* (BioRep) to assess whether an evaluation protocol reflects a continuous covariate associated with perturbation strength, such as drug dosage, developmental time, or the number of guide RNAs. A good protocol should assign larger scores as the covariate increases. In practice, BioRep is defined as the Spearman correlation between scores and the corresponding covariate. Unlike the other criteria, BioRep evaluates whether a protocol is sensitive to graded biological variation. This is most informative for magnitude-sensitive metrics, whereas scale-insensitive metrics such as Pearson correlation can remain similar as perturbation magnitude increases. A further caveat is that the covariate may not track perturbation strength monotonically, for instance when dose responses saturate.

### 6.4 Complementarity

No single evaluation protocol captures every aspect of perturbation-response prediction, so benchmarks often report a panel of protocols. For this panel to be informative, the protocols must be complementary rather than redundant. For example, a metric sensitive to the magnitude of the response, such as L2 distance, can be paired with one sensitive to the relative pattern of change, such as Pearson correlation on delta vectors. A common empirical diagnostic is to compute pairwise correlations between protocol scores across a set of model predictions, where highly correlated scores suggest partly redundant protocols [66, 82]. However, these correlations depend on the models evaluated and may reflect shared failure modes rather than an intrinsic relationship between the protocols.

### 6.5 Scalability & Estimation Error

Evaluation protocols differ not only in what they measure but also in how expensive they are to compute and how reliably their scores can be estimated. Both matter in single-cell settings, where each condition may be represented by relatively few cells in a high-dimensional gene space, and modern datasets can contain thousands of conditions. The computational burden depends on how a protocol compares populations. Pseudobulk-based protocols are relatively cheap: they reduce each population to a single vector and avoid pairwise cell–cell comparisons. Protocols that operate on the full distribution of cells, such as distributional metrics and differential-expression pipelines, are substantially more expensive (Supplementary Table 1).

Estimation error in single-cell settings has two main drivers: the number of cells per population and the feature dimension. These are especially problematic in combination, as in single-cell data, where high dimensionality coincides with limited sample size, leaving even a conceptually appealing distributional metric with a noisy or biased estimate. Wasserstein distances are a clear example: the empirical estimator converges slowly in high dimensions and can show substantial upward bias even when the two distributions are identical [32, 36, 107]. Cost and estimation error tend to go together: comparing full cell distributions is expensive and hard to estimate, whereas reducing each population to a pseudobulk or centroid lowers both.

## 7 Outlook & Discussion

Here, we present a taxonomy of evaluation protocols for single-cell perturbation prediction models, as well as criteria for assessing these protocols. The taxonomy highlights that evaluation protocols comprise more than a single metric: they involve many design choices, from feature selection, embeddings, and isolation of perturbation effects to score transformation and aggregation. Making these choices explicit reveals shared structure across existing approaches and clarifies the assumptions, trade-offs, and failure modes that arise throughout the evaluation pipeline.

We focus on the most common evaluation setting: directly comparing observed and predicted perturbation responses. Complementary work evaluates predictions not by comparison to matched observed responses, but by asking whether predicted effects recover external biological or pharmacological structure, such as grouping small molecules by mechanism of action [29] or reflecting orthogonal information about perturbations, such as the sensitivity of cancer cell lines to small molecule treatments [12]. Another line of work asks whether models can infer causal relationships that explain how biological systems respond to interventions [15, 19, 98]. In biological systems, however, ground-truth causal relationships are rarely available, so these benchmarks often rely on indirect evaluation, including whether inferred networks can predict perturbation-induced gene expression changes [88, 98].

Although we focus on transcriptomic readouts, many of the evaluation principles discussed here extend to other high-dimensional single-cell perturbation-response measurements. These include established imaging readouts such as Cell Painting [17], as well as emerging readouts such as spatial transcriptomics [18, 53, 94] and multi-modal assays, including CITE-seq [33] and paired RNA–ATAC measurements [34]. The specific representations will differ across modalities, but the core questions remain the same: which aspects of the perturbation response are emphasized by a given representation, which combinations of representations, metrics, and cross-perturbation evaluations are aligned with the intended use case, and which score aggregation scheme supports meaningful model selection.

At present, the field of single-cell perturbation prediction lacks consensus on which tasks should be prioritized and which evaluation protocols are appropriate for each task. This fragmentation limits comparability across studies and makes it difficult to determine whether new models represent genuine progress. Similar concerns in biomedical image analysis motivated Metrics Reloaded, a community-driven effort that resulted from a multistage Delphi process, workshops, surveys, expert meetings, and crowdsourced feedback, and produced a problem-aware framework for metric selection [64]. We envision an analogous process for the single-cell perturbation modeling community. By organizing existing evaluation choices into a taxonomy and outlining criteria for assessing them, this work provides a starting point to discuss the development of task-aware evaluation standards in the field. Developing such standards will require input from across the community: experimental biologists who understand the intricacies of perturbation screens, statisticians and mathematicians who study metric behavior, and computational biologists and machine-learning researchers who develop and benchmark predictive models. We hope that this work helps catalyze a broad and inclusive discussion, and we invite the community to join forces to help shape the next generation of evaluation standards for single-cell perturbation modeling.

## Code availability

Reference implementations of selected evaluation protocols are provided in the accompanying Python package scPertEval, available at https://github.com/Virtual-Cell-Research-Community/scPertEval. Code for reproducing the analyses and figures presented in this manuscript is available at https://github.com/Virtual-Cell-Research-Community/scPertEval-figures.

## Data availability

The datasets analyzed in this study are publicly available. Dataset names in italics correspond to the identifiers used throughout the manuscript.

***Wessels23*** The dataset from Wessels et al. [111] was obtained through scPerturb [78] and is available at https://zenodo.org/records/13350497/files/WesselsSatija2023.h5ad?download=1.

***Replogle22 RPE1*** The RPE1 dataset from Replogle et al. [85] was obtained through scPerturb [78] and is available at https://zenodo.org/records/13350497/files/ReplogleWeissman2022_rpe1.h5ad?download=1.

***Replogle22 K562*** The K562 genome-wide Perturb-seq dataset from Replogle et al. [85] was obtained through scPerturb [78] and is available at https://zenodo.org/records/13350497/files/ReplogleWeissman2022_K562_gwps.h5ad?download=1.

***Nadig25 Jurkat*** The Jurkat dataset from Nadig et al. [72] was obtained through scPerturb [78] and is available at https://zenodo.org/records/13350497/files/NadigOConner2024_jurkat.h5ad?download=1.

***Nadig25 HepG2*** The HepG2 dataset from Nadig et al. [72] was obtained through scPerturb [78] and is available at https://zenodo.org/records/13350497/files/NadigOConner2024_hepg2.h5ad?download=1.

***ArcH1*** The dataset from the Virtual Cell Challenge [91] is available at https://virtualcellchallenge.org/datasets.

***Kaden25 RPE1*** The RPE1 CRISPRa dataset from Southard et al. [97] is available at https://zenodo.org/records/15200179/files/fibroblast_CRISPRa_final_pop_singlets_normalized_log1p.h5ad?download=1.

## Contributions

PSLS performed the literature review, developed the initial manuscript structure, wrote the first draft, and designed the schematic figures. KAR initiated the project, assembled the research team, managed the collaborative infrastructure, and provided critical review throughout manuscript revisions. KAR and PSLS jointly formulated the core research problem. ZB curated the project data, led the development of the accompanying software package, performed the computational analyses, and generated the computational figures. HH made substantial contributions to the implementation and documentation of the software package. All authors critically reviewed the manuscript, contributed to the final taxonomy, and approved the final version. JSR supervised the work.

## Acknowledgments

The authors thank Pablo Rodriguez-Mier, Katharina Mikulik, Daniel Dimitrov, Constantin Ahlmann-Eltze, Eric Kernfeld, Bence Szalai, Adam Krejci, Anaïs Baudot, and Jean Radig for their helpful comments and suggestions.

The authors also thank Nathan LaPierre and Abid Rehman for early discussions on the topic of this work.

## Supplementary Material

## A Supplementary Figures

**Supplementary Figure 1:**
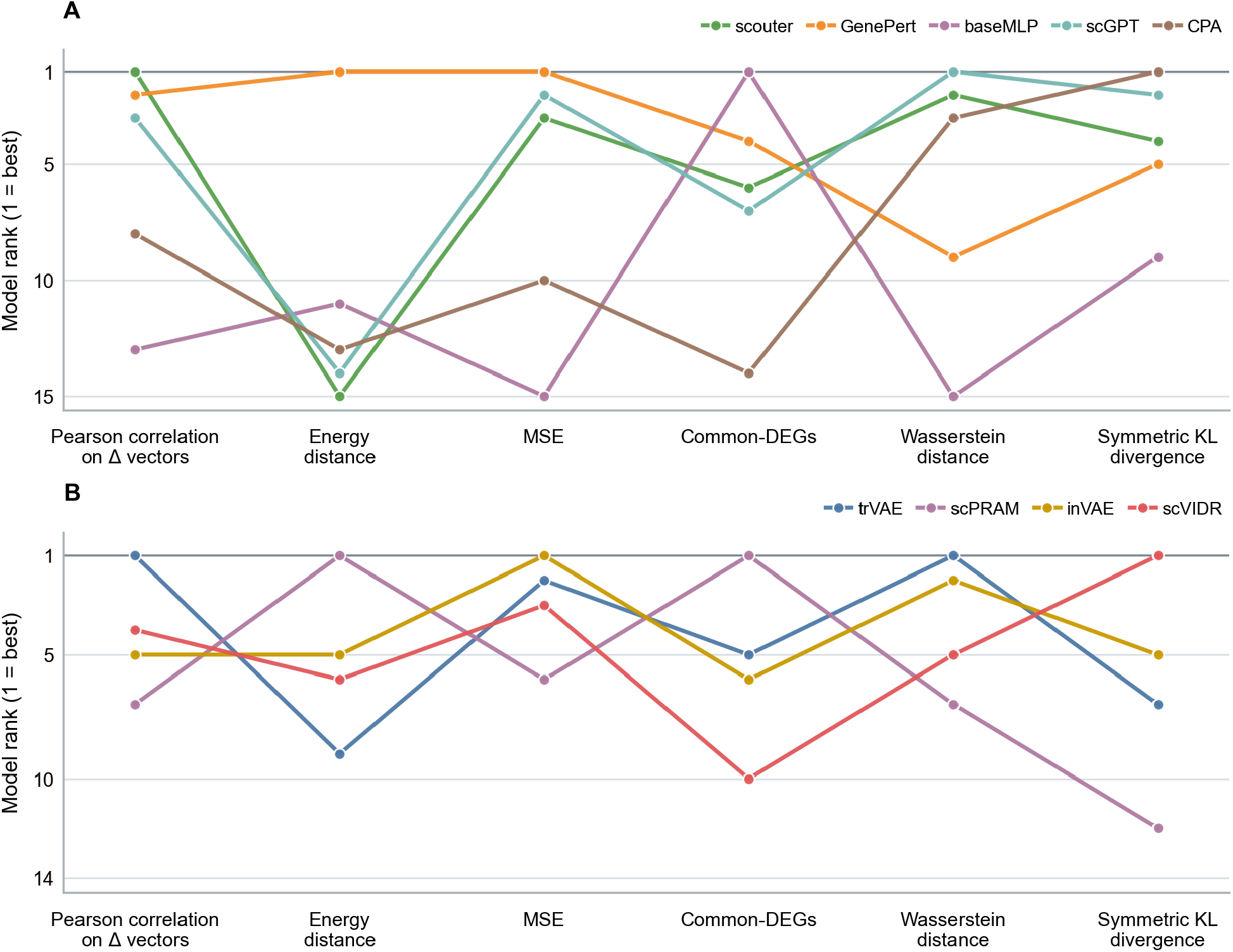
Model rankings depend on the evaluation metric. **(A)** Model rankings were derived from the single-gene perturbation prediction benchmark of Wei et al. [71], which evaluated 15 models on nine datasets. We used the results computed on the top 100 most differentially expressed genes per perturbation. Dataset-level ranks for each model and metric were averaged across datasets with equal weight and re-ranked to obtain the ordinal ranks shown. Lines connect the ranks of the same model across the six metrics; lower ranks indicate better performance. For clarity, only models that rank first under at least one metric are shown, although all 15 models were included in the rank calculations. Across the six metrics, five distinct models attain the top rank. **(B)** The corresponding analysis for out-of-distribution cellular-context generalization used the same gene-selection and rank-aggregation procedures and included 12 datasets and 14 models. Across the six metrics, four distinct models attain the top rank.

**Supplementary Figure 2:**
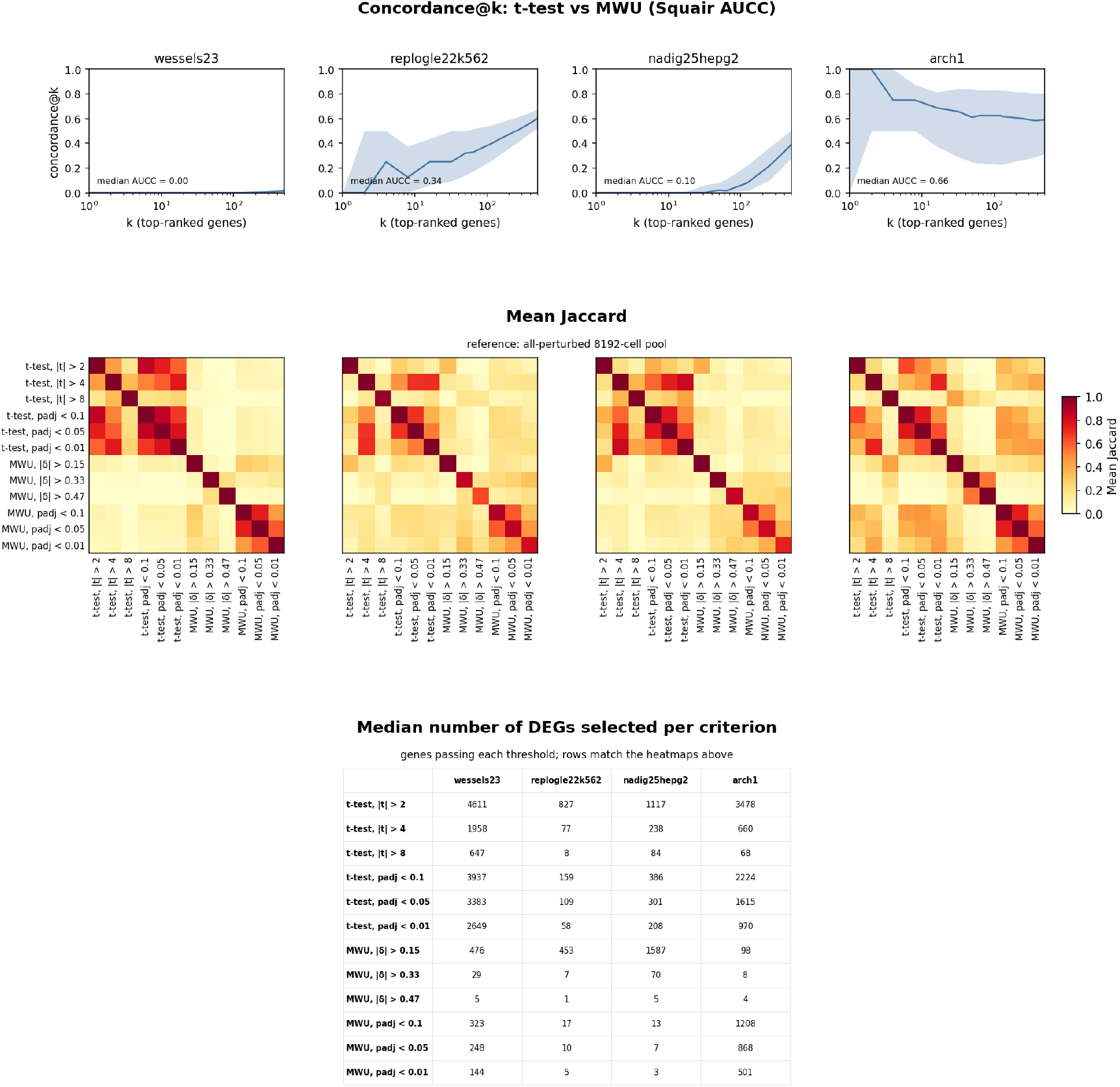
Differential-expression methodology substantially affects DEG calls. For each dataset, differential-expression statistics were computed using a *t*-test and Mann–Whitney U test (MWU) against an all-perturbed reference pool of 8,192 cells. The upper panels show concordance at *k* between the *t*-test and MWU rankings, defined as the fraction of shared genes among the top-*k* genes after ranking by adjusted *p*-value with absolute test statistic used as a tie-breaker. Lines show the median across perturbations, shaded bands show the interquartile range, and each panel reports the median area under the concordance curve (AUCC), computed as the mean concordance over the evaluated *k* grid. The middle panels show the mean Jaccard index between DEG sets defined by threshold-based selection rules for the same two DE procedures, including *t*-test thresholds on | *t* | and adjusted *p*-values and MWU thresholds on |*δ*| and adjusted *p*-values. Higher Jaccard values indicate stronger agreement between the corresponding DEG sets. The bottom table reports the median number of genes selected by each criterion. Overall, agreement between DE procedures and DEG-selection rules varies substantially across datasets and depends both on the statistical test and on the stringency of the selection criterion.

**Supplementary Figure 3:**
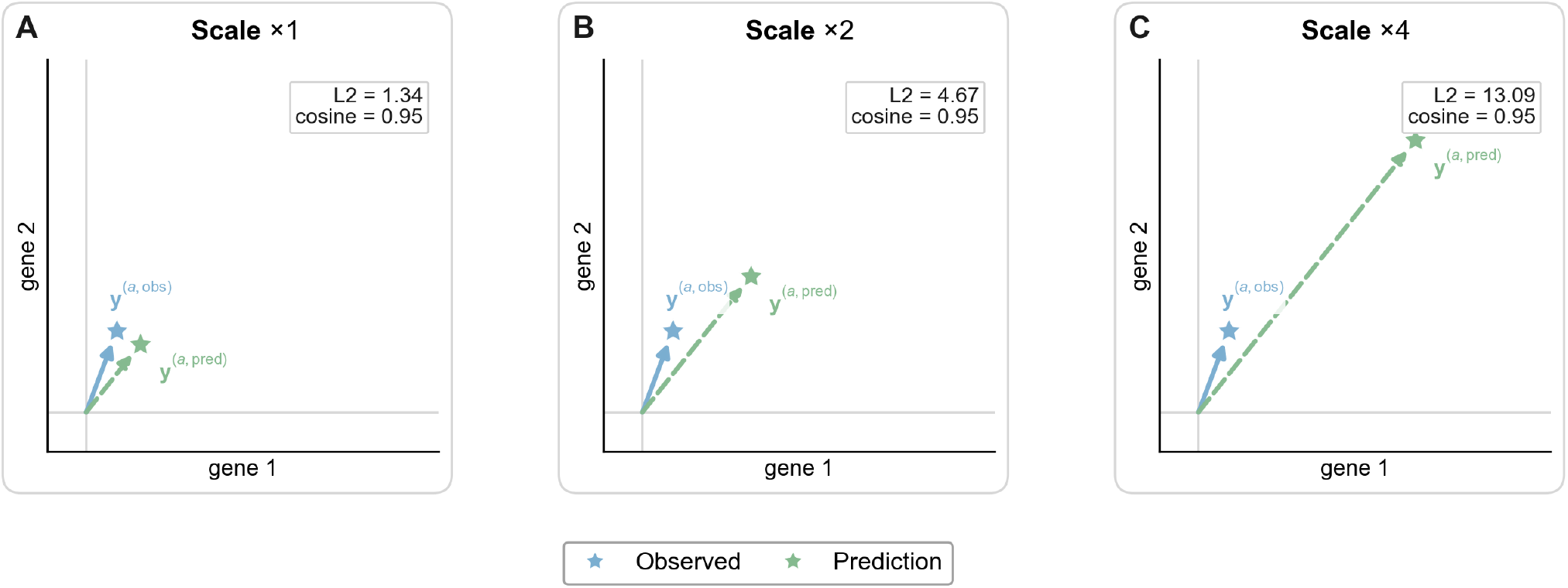
Scale invariance of similarity metrics. A ground-truth perturbation vector (blue) is compared with corresponding prediction vectors (green) that preserve the same overall direction but increase in magnitude from panels A to C. As the prediction is scaled up, the L2 distance grows substantially, whereas cosine similarity stays constant because the angular relationship between the vectors is preserved. This illustrates that L2 is sensitive to absolute scale, while cosine similarity only reflects directional agreement.

**Supplementary Figure 4:**
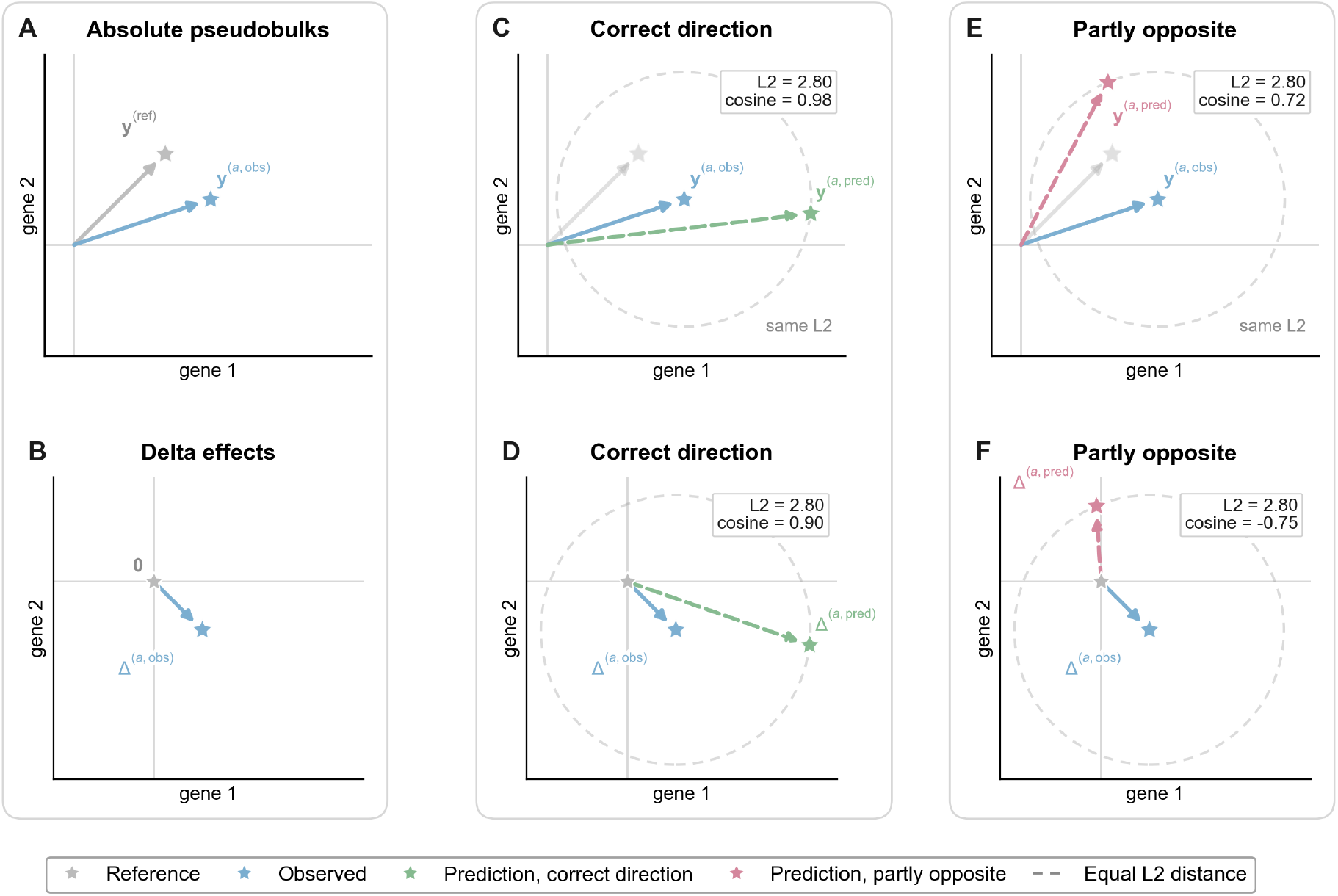
Sensitivity to perturbation direction in absolute-expression and control-centered delta space. Each column shows one scenario in absolute coordinates (top) and the corresponding control-centered delta (bottom). Panel A shows the control (gray) and ground-truth perturbation (blue) in absolute coordinates, and panel B the matching ground-truth delta vector. Panels C and E add two predictions that are equidistant from the ground truth according to L2 distance, as indicated by the dashed circle: the prediction in panel C follows the same direction of change from the control, whereas the prediction in panel E reverses it. Panels D and F show the corresponding control-centered delta predictions, sign-consistent in panel D and sign-inconsistent in panel F. In both rows, the paired predictions have the same L2 error, but Pearson correlation and cosine similarity distinguish preserved from reversed perturbation direction, with the contrast becoming especially explicit in delta space.

**Supplementary Figure 5:**
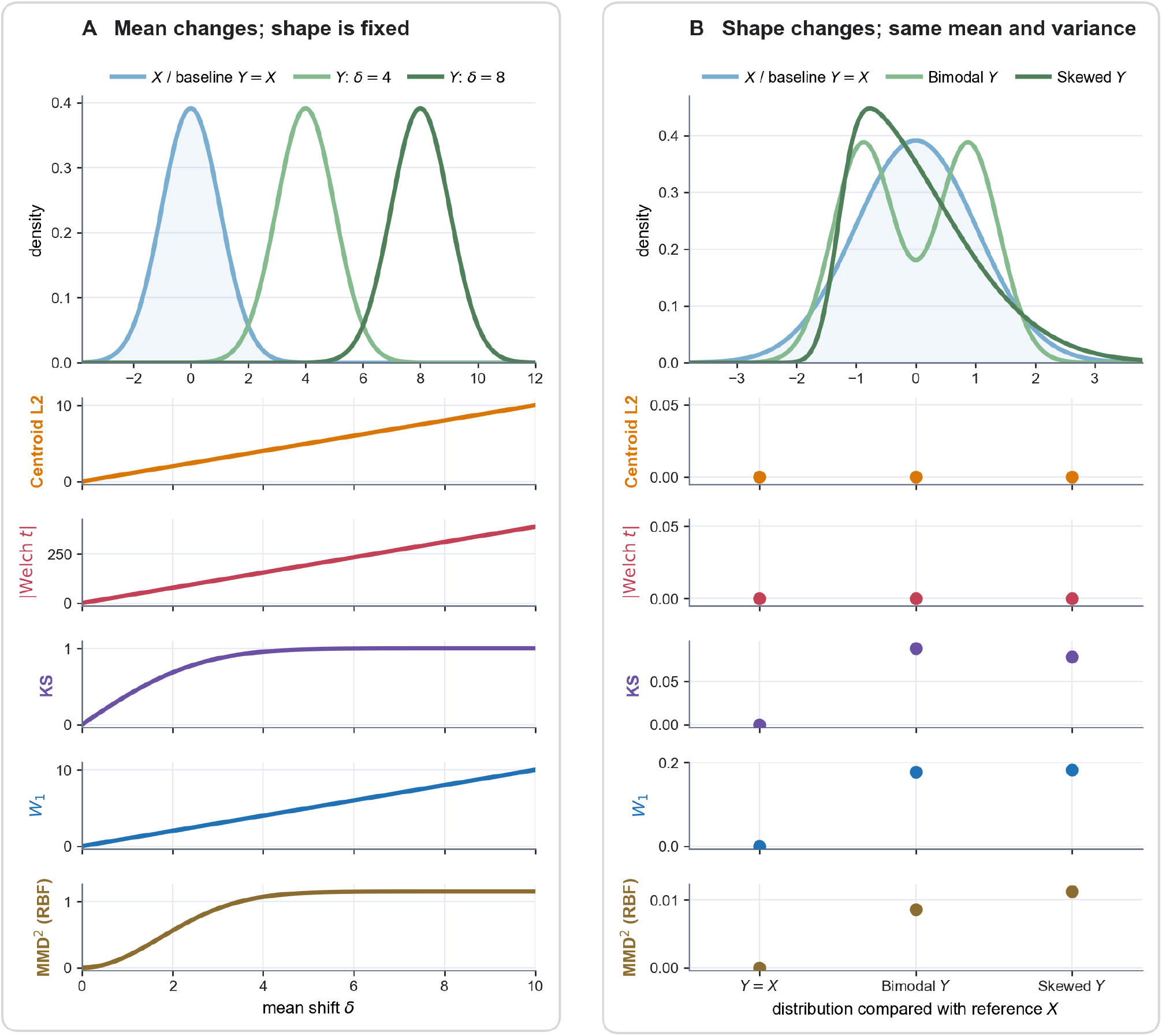
Distribution-sensitive metrics capture population changes missed by mean-based comparisons. **(A)** A reference distribution *X* is compared with *Y* = *X* + *δ* across increasing mean shifts *δ*. Centroid L2, the absolute Welch *t*-statistic, and Wasserstein-1 (*W*_1_) increase with the shift, whereas the bounded KS statistic and RBF MMD^2^ saturate. **(B)** The baseline *Y* = *X* is compared with bimodal and right-skewed alternatives matched to *X* in empirical mean and variance. Consequently, Centroid L2 and the Welch *t*-statistic remain zero, while KS, *W*_1_, and MMD^2^ detect changes in distribution shape. Metrics are shown on their native scales. MMD^2^ denotes the biased empirical maximum mean discrepancy using a Gaussian RBF kernel with bandwidth *σ* = 1.

**Supplementary Figure 6:**
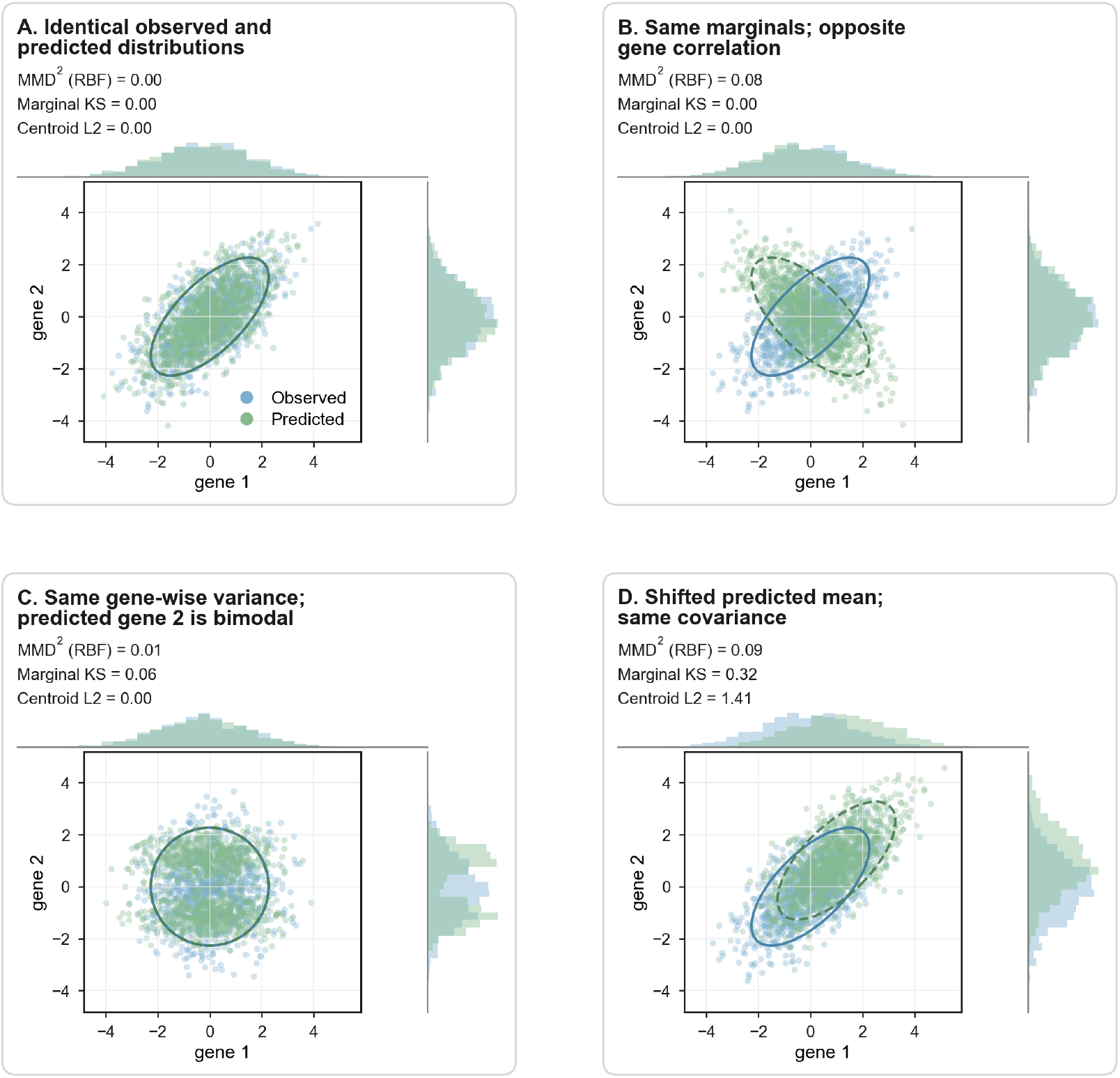
Centroid, marginal, and joint distribution metrics comparing synthetic observed and predicted two-gene populations. **(A)** The observed and predicted populations have identical means and covariance structures, so all metrics are zero. **(B)** The populations have identical one-dimensional marginals but opposite cross-gene correlations. Consequently, the marginal KS and Centroid L2 metrics remain zero, whereas MMD^2^ detects the altered joint dependence structure. **(C)** The populations have the same centroid and gene-wise variances, but predicted gene 2 is bimodal. Marginal KS and MMD^2^ detect this distributional difference, while Centroid L2 remains zero. **(D)** The predicted population has a shifted mean but the same covariance as the observed population, producing discrepancies in the centroid, marginal, and joint distribution metrics. Together, these examples illustrate that centroid metrics cannot detect changes in distributional shape and that both marginal distribution metrics and centroid metrics cannot detect changes in gene–gene dependencies.

**Supplementary Figure 7:**
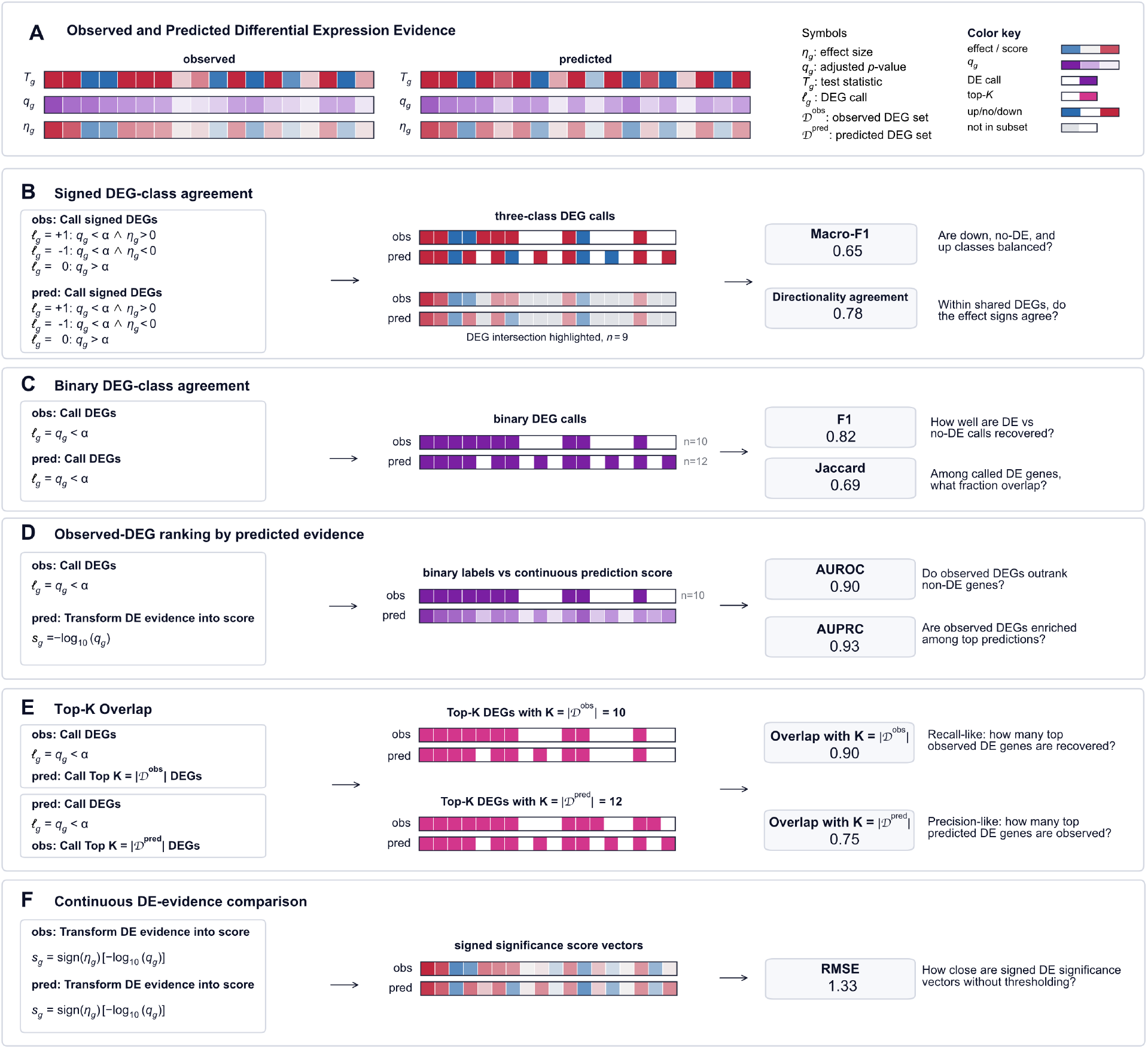
Differential-expression metric. **(A)** Observed and predicted differential-expression analyses provide gene-wise evidence, including effect sizes (*η*_*g*_), adjusted *p*-values (*q*_*g*_), test statistics (*T*_*g*_), and DEG-call labels (*ℓ*_*g*_). The remaining panels show representative transformations of these outputs into signed DEG classes, binary DEG calls, observed-DEG ranking labels, top-*K* DEG lists, or continuous signed scores, followed by the evaluation metric applied to each representation. These examples reflect transformation choices used in the literature for calling DEGs or scoring DEG evidence for ranking, but they are not exhaustive; other thresholds, scores, and class definitions are possible. **(B)** Observed and predicted genes are assigned signed DEG labels using *q*_*g*_ *< α* and the sign of *η*_*g*_, then compared with macro-F1 and directionality agreement within shared DEGs. **(C)** Observed and predicted outputs are thresholded into binary DEG calls and compared with binary F1 and Jaccard similarity. **(D)** Observed DEG calls are treated as binary labels and predicted evidence is retained as a continuous ranking score, evaluated with AUROC and AUPRC. **(E)** Observed and predicted evidence are converted into top-*K* DEG lists, with *K* defined by either the number of observed or predicted DEGs, and compared using top-*K* overlap. **(F)** Continuous signed DE evidence is compared directly after transforming adjusted *p*-values into signed significance scores, evaluated with RMSE.

**Supplementary Figure 8:**
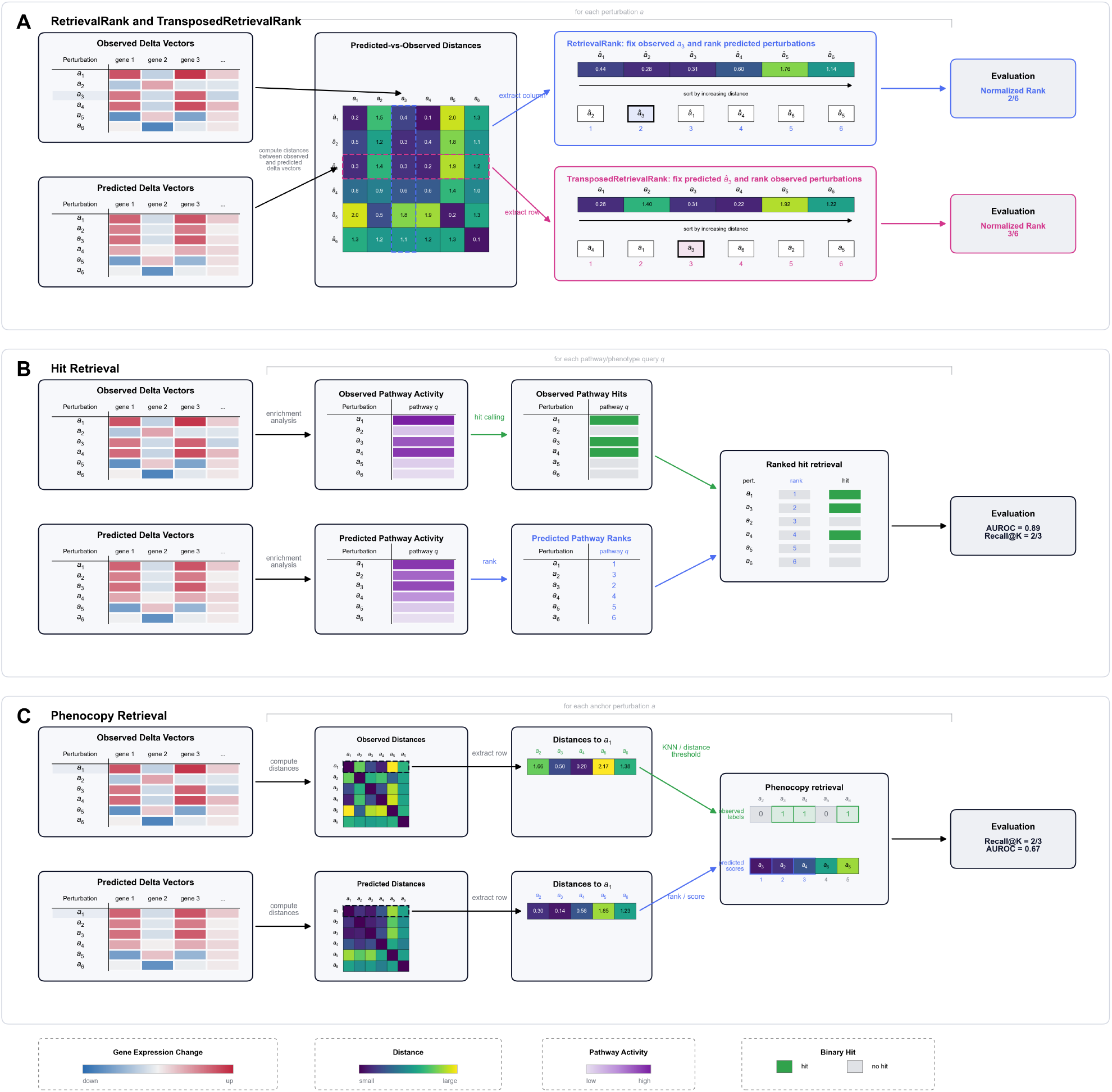
Retrieval-based evaluation of perturbation-response predictions. **(A)** Retrieval rank and transposed retrieval rank are derived from the matrix of distances between predicted and observed perturbation delta vectors. Retrieval rank fixes an observed response as the query and ranks all predicted responses, whereas transposed retrieval rank fixes a predicted response and ranks all observed responses. The matched perturbation’s normalized rank is the evaluation score. **(B)** In hit retrieval, observed perturbation responses are converted into binary hit labels for a pathway or phenotype, while predicted responses are converted into continuous activity scores and ranked. The resulting ranking is compared with the observed hit labels using metrics such as AUROC or Recall@*K*. **(C)** Phenocopy retrieval first defines observed phenocopies of an anchor perturbation using observed-response distances, for example through a nearest-neighbor or distance-threshold rule. Candidate perturbations are then ranked by their predicted similarity to the anchor, and agreement with the observed phenocopy labels is summarized using Recall@*K*, AUROC, or related ranking metrics.

**Supplementary Figure 9:**
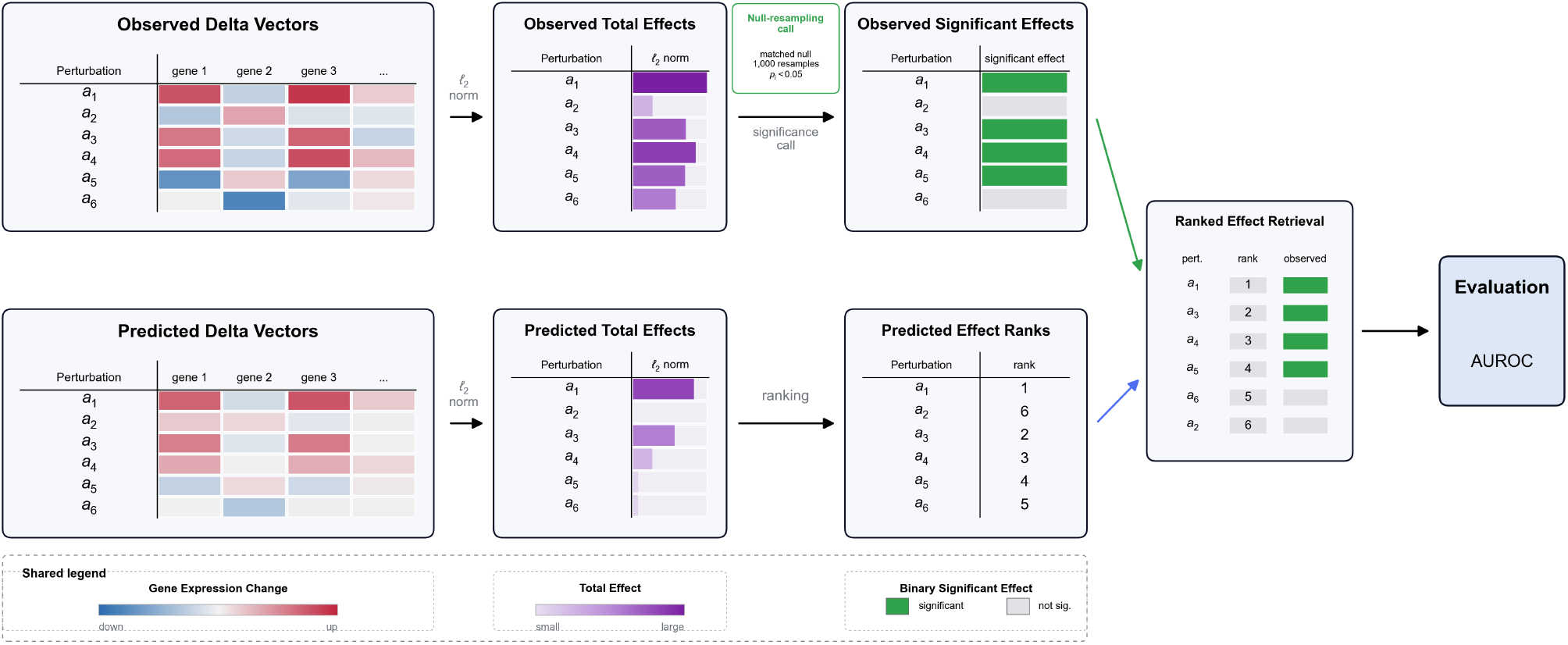
Effect-size evaluation of perturbation-response predictions introduced in Littman et al. [40]. The *ℓ*_2_ norm of the control-centered pseudobulk delta vector for condition *a* (with *N*_*a*_ cells) is used to define the observed total effect. To account for the dependence of pseudobulk noise on cell number, a matched null distribution is generated by repeatedly sampling *N*_*a*_ cells without replacement from all perturbed cells in the dataset, constructing control-centered pseudobulk vectors, and computing their *ℓ*_2_ norms. The one-sided empirical *p*-value is the fraction of null norms exceeding the observed norm, and perturbations with nominal *p*_*i*_ *<* 0.05 receive positive observed-effect labels. Predicted delta vectors are scored by their *ℓ*_2_ norms and ranked by predicted total effect. Comparing this ranking with the binary observed labels yields an effect-size classification task summarized by AUROC.

**Supplementary Figure 10:**
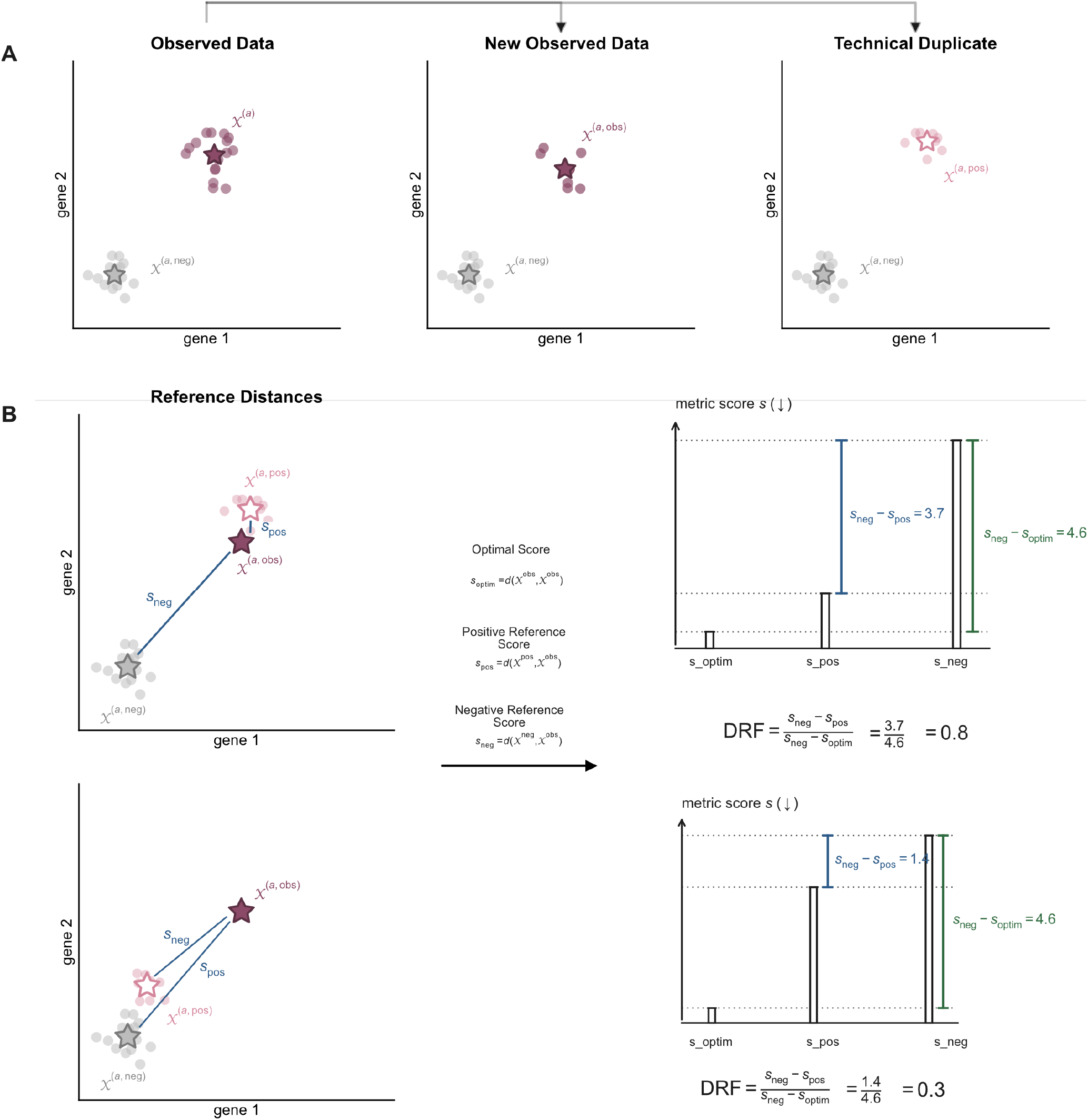
Construction of a technical duplicate and interpretation of the dynamic range fraction. **(A)** Cells from a perturbation are split into two disjoint subsets: one subset defines the observed ground truth *X*^(*a*,obs)^ and the other defines the technical duplicate used as the positive reference *X*^(*a*,pos)^. *An* uninformative population *X*^(*a*,neg)^ serves as the negative reference. **(B)** For a benchmark metric in which lower values are better, *s*_optim_ is the score obtained by comparing the ground truth with itself, *s*_pos_ compares the positive reference with the ground truth, and *s*_neg_ compares the negative reference with the ground truth. The DRF is the fraction of the available score range recovered by the positive reference, shown schematically as (*s*_neg_ − *s*_pos_)*/*(*s*_neg_ − *s*_optim_). A positive reference close to the ground truth produces a high DRF, illustrated by 3.7*/*4.6 = 0.8, whereas a positive reference close to the negative reference produces a low DRF, illustrated by 1.4*/*4.6 = 0.3.

## B Supplementary Tables

**Supplementary Table 1:** Runtime, in seconds, of evaluation-protocol configurations across datasets. Rows specify the base metric and associated choices for reference centering, representation, controls, feature space, and weighting; dataset columns report individual runtimes and the final column reports their sum. Darker cells indicate longer runtimes. **Results**. Pseudobulk MSE and WMSE are among the least expensive protocols (approximately 9–10 seconds in total), and pseudobulk rank-based protocols remain inexpensive (approximately 32 seconds). Single-cell distributional comparisons are substantially more costly: energy distance requires approximately 900 seconds, MMD approximately 2,400 seconds, and Sinkhorn *W*_2_ approximately 3,700–4,400 seconds. Perturbation-centered Pearson correlation is also comparatively expensive in this implementation (approximately 650 seconds). Reported runtimes exclude upstream differential-expression computation; configurations that use DE results to restrict the feature space or construct WMSE weights therefore incur additional cost not shown here.

| Base Metric | Reference Centering | Representation | Positive Control | Negative Control | Feature Space | Weight Exponent | Wessels23 | Replogle22 RPE1 | Replogle22 K562 | Nadig25 Jurkat | Nadig25 Hepg2 | ArcH1 | Kaden25 RPE1 | TOTAL (s) |
| --- | --- | --- | --- | --- | --- | --- | --- | --- | --- | --- | --- | --- | --- | --- |
| Pearson | — | Pseudobulk | Interpolated TD | All-pert. (mean) | — | — | 0.3 | 2.2 | 2.2 | 2.2 | 1.9 | 1.8 | 7.2 | 17.9 |
| Pearson | Ctrl | Pseudobulk | Interpolated TD | All-pert. (mean) | — | — | 0.2 | 0.8 | 0.7 | 0.8 | 0.7 | 1.0 | 5.5 | 9.6 |
| Pearson | Pert | Pseudobulk | Interpolated TD | Control (mean) | — | — | 0.6 | 96.1 | 93.5 | 108.1 | 47.1 | 15.5 | 289.3 | 650.2 |
| Pearson | Pert | Pseudobulk | Interpolated TD | Control (mean) | Sig. DEGs | — | 0.7 | 97.4 | 96.1 | 109.9 | 47.6 | 15.7 | 292.0 | 659.4 |
| Pearson | Pert | Pseudobulk | Interpolated TD | Control (mean) | Top-50 DEGs | — | 0.6 | 96.0 | 94.5 | 107.7 | 47.1 | 15.7 | 287.8 | 649.5 |
| MSE | — | Pseudobulk | Interpolated TD | All-pert. (mean) | — | — | 0.1 | 0.7 | 0.7 | 0.7 | 0.5 | 1.1 | 5.3 | 9.1 |
| MSE | — | Pseudobulk | Interpolated TD | All-pert. (mean) | Sig. DEGs | — | 0.2 | 0.7 | 0.7 | 0.8 | 0.6 | 1.0 | 5.3 | 9.4 |
| MSE | — | Pseudobulk | Interpolated TD | All-pert. (mean) | Top-50 DEGs | — | 0.2 | 0.8 | 0.8 | 0.8 | 0.7 | 1.0 | 5.5 | 9.8 |
| WMSE | — | Pseudobulk | Interpolated TD | All-pert. (mean) | — | 1.0 | 0.2 | 0.9 | 0.9 | 1.0 | 0.8 | 1.0 | 5.5 | 10.3 |
| WMSE | — | Pseudobulk | Interpolated TD | All-pert. (mean) | — | 2.0 | 0.1 | 0.8 | 0.7 | 0.8 | 0.6 | 1.0 | 5.3 | 9.4 |
| WMSE | — | Pseudobulk | Interpolated TD | All-pert. (mean) | — | 4.0 | 0.1 | 0.8 | 0.8 | 0.8 | 0.6 | 1.0 | 5.4 | 9.4 |
| TransposedRank | — | Pseudobulk | Interpolated TD | All-pert. (mean) | — | — | 0.3 | 2.0 | 2.3 | 2.2 | 1.4 | 3.7 | 19.6 | 31.6 |
| Rank | — | Pseudobulk | Interpolated TD | All-pert. (mean) | — | — | 0.3 | 2.2 | 2.4 | 2.4 | 1.6 | 3.7 | 19.8 | 32.5 |
| DE Overlap @ 50 | — | Single-cell | Technical duplicate | All-pert. {8192 cells} | — | — | 0.0 | 0.3 | 0.3 | 0.3 | 0.3 | 0.0 | 0.3 | 1.5 |
| DE AUPRC | — | Single-cell | Technical duplicate | All-pert. {8192 cells} | — | — | 0.3 | 2.7 | 2.8 | 2.9 | 2.8 | 0.3 | 3.3 | 15.0 |
| DE AUROC | — | Single-cell | Technical duplicate | All-pert. {8192 cells} | — | — | 0.3 | 3.1 | 3.1 | 3.2 | 2.9 | 0.3 | 3.4 | 16.4 |
| MMD | — | Single-cell | Technical duplicate | All-pert. {8192 cells} | Top-50 DEGs | — | 33.1 | 498.4 | 488.0 | 528.0 | 432.4 | 36.0 | 409.7 | 2425.5 |
| MMD | — | Single-cell | Technical duplicate | All-pert. {8192 cells} | PCA-50 | — | 33.6 | 496.8 | 476.3 | 523.1 | 421.6 | 35.2 | 467.7 | 2454.3 |
| ED | — | Single-cell | Technical duplicate | All-pert. {8192 cells} | Top-50 DEGs | — | 12.6 | 184.1 | 183.0 | 197.3 | 162.4 | 12.7 | 157.9 | 909.9 |
| ED | — | Single-cell | Technical duplicate | All-pert. {8192 cells} | PCA-50 | — | 12.4 | 180.1 | 177.7 | 186.8 | 157.0 | 13.4 | 165.7 | 893.2 |
| $W_2$ (Sinkhorn) | — | Single-cell | Technical duplicate | All-pert. {8192 cells} | Top-50 DEGs | — | 50.3 | 762.0 | 760.9 | 803.8 | 662.5 | 53.9 | 650.8 | 3744.2 |
| $W_2$ (Sinkhorn) | — | Single-cell | Technical duplicate | All-pert. {8192 cells} | PCA-50 | — | 59.6 | 891.8 | 884.7 | 931.5 | 802.9 | 63.6 | 804.2 | 4438.2 |
| **TOTAL (s)** |  |  |  |  |  |  | **206** | **3321** | **3273** | **3515** | **2796** | **279** | **3617** | **17006** |

## C Supplementary Primers

### C.1 Primer on Statistical Models for Single-Cell RNA-seq Data

Throughout this primer, we take unique molecular identifier (UMI)-based scRNA-seq as the default setting. Let *n* index cells and *g* index genes. The observed measurement for gene *g* in cell *n* is a non-negative integer count *X*_*ng*_. A useful starting point is to separate a cell-specific exposure or size factor *l*_*n*_ from a gene-specific relative expression level *λ*_*g*_ with ∑_*g*_ *λ*_*g*_ = 1 and to define the mean count as

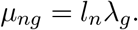

A simple sampling model is then

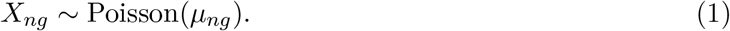

The term *l*_*n*_ is treated as part of the sampling model rather than as a biological effect. Under the Poisson model, the variance equals the mean,

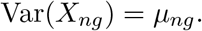

Real scRNA-seq data often show extra variability beyond this Poisson baseline. In the simple mean model above, all cells share the same feature-specific relative expression level *λ*_*g*_ after accounting for exposure *l*_*n*_, so residual cell-to-cell heterogeneity and technical variation appear as overdispersion around *µ*_*ng*_. A common way to model this overdispersion is to assume that the Poisson rate itself varies according to a gamma distribution. To avoid confusion with the relative expression level *λ*_*g*_, we denote this random latent Poisson rate by Θ_*ng*_. In the Gamma–Poisson model,

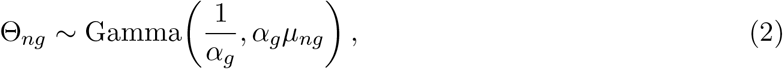

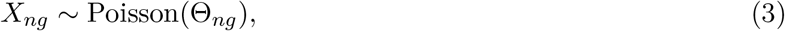

where the gamma distribution is parameterized by shape and scale. This parameterization gives

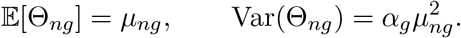

After marginalizing over the latent rate Θ_*ng*_, the count has mean

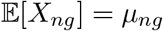

and variance

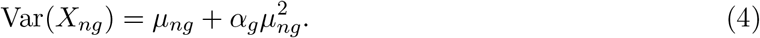

Here, *α*_*g*_ *>* 0 quantifies feature-specific overdispersion, and the Poisson model is recovered in the limit *α*_*g*_ → 0.

The same marginal count distribution is commonly referred to as a negative-binomial distribution. To avoid ambiguity from different negative-binomial parameterizations, it is useful to write the model in Gamma–Poisson form and then note the equivalence. Under a common parameterization of the negative-binomial distribution with shape parameter *r* and success probability *p*, chosen so that the mean is *r*(1 − *p*)*/p*, the Gamma–Poisson model above corresponds to

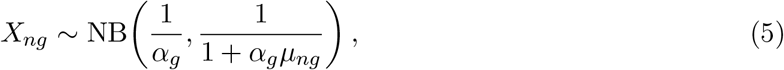

which has mean *µ*_*ng*_ and variance 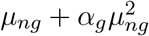.

This Gamma–Poisson, or negative-binomial, model is the standard count-modeling perspective behind many likelihood-based single-cell methods. Gamma–Poisson models also underlie residual-based transformations, including regularized negative-binomial regression in SCTransform [26] and analytic Pearson-residual approaches based on parsimonious offset models [37].

Readers may also encounter zero-inflated variants such as zero-inflated Poisson (ZIP) or zero-inflated negative binomial (ZINB), which add an explicit extra-zero component to the count model. However, it has been argued that explicit zero inflation is often unnecessary for UMI-count data [9, 33, 64].

These modeling choices matter for metric interpretation throughout the review. Count likelihoods such as Poisson or negative-binomial negative log-likelihoods are naturally defined on raw counts and should include appropriate depth or exposure handling, whereas correlation or L1 */* L2 distances should be applied after some preprocessing or transformation step (Supplementary Section C.2).

For further reading, Choudhary and Satija [11] compare and evaluate scRNA-seq error models in detail, and Heumos et al. [27] provide a broader practical overview. Assumptions about count distributions, library-size effects, and gene–gene dependence are also encoded in simulators for single-cell data [39, 62, 75].

### C.2 Primer on Normalization of Single-Cell RNA-sequencing Data

If one analyzes raw UMI counts with a count-based likelihood, then the cell-specific library size *l*_*n*_ is handled inside the model, typically through an offset or exposure term, rather than by creating a separately normalized count matrix. This is natural for methods whose likelihood already matches count data, such as negative-binomial latent-variable models like scVI or multinomial/Poisson generalized PCA variants such as GLM-PCA [11, 41, 66].

If instead one wants a transformed feature matrix for methods that expect homoskedastic data such as PCA, then depth correction and variance-stabilizing transformation become necessary preprocessing steps. Common choices include simple library-size normalization followed by log(1 + *x*), deconvolution-based size factors [44], and residual-based approaches such as regularized negative-binomial regression or analytic Pearson residuals [26, 37]. The goal is usually not to make the data exactly Gaussian, but to reduce mean-variance dependence enough that methods based on Gaussian assumption become more appropriate [3].

For further reading, Ahlmann-Eltze and Huber [3] compare several transformation families in detail, while [27] provides general guidance on normalization.

### C.3 Primer on Highly Variable Gene Selection

Highly variable genes (HVGs) are genes whose variability is larger than expected given their average expression level. This qualification matters because under the count models discussed in Supplementary Section C.1, variance increases with the mean. If one simply ranks genes by raw variance, the result is dominated by the most highly expressed genes rather than by the genes that are most informative about differences between cell states or perturbations.

Common HVG workflows therefore try to estimate the mean-variance trend, and then rank genes by excess variability or overdispersion relative to that baseline. In practice, many single-cell pipelines use Seurat- or Scanpy-style HVG procedures [27, 76]. In multi-sample or multi-batch datasets, the choice between global and batch-aware HVG selection can affect the resulting gene set. Pooled selection will prioritize genes that distinguish samples or batches, whereas batch-aware variants aim to identify genes that show high within-sample variability across multiple samples.

For further reading, Zappia et al. [76] benchmark feature-selection strategies for single-cell integration and querying, while [27] provides broader workflow guidance.

### C.4 Primer on Differential Gene Expression in Single-Cell Perturbation Data

Differential expression (DE) analysis compares a perturbed cell population to a reference population, such as control cells, to identify genes whose expression changes significantly in response to the perturbation. For each gene, a DE procedure typically returns a test statistic, a raw and an adjusted *p*-value, and an effect-size estimate such as a log-fold change. These gene-level outputs are central to how perturbation experiments are interpreted, and they enter model evaluation in two ways: as a basis for feature selection or weighting (Section 2.2) and as the object directly compared by differential-expression metrics (Section 3.5). General overviews of DE analysis in single-cell data are given in [27, 63, 67].

A wide range of DE methods is used in single-cell perturbation studies, and the choice of method varies considerably across the field. Simple two-sample tests include the Wilcoxon rank-sum test [47, 51, 59, 61, 72] and the *t*-test [45, 46, 73], both available in standard toolkits such as Scanpy and Seurat, as well as the Kolmogorov–Smirnov test [1, 54]. More specialized models for scRNA-seq include MAST [21] and Memento [34]. Furthermore, several methods have been designed for pooled CRISPR screens, including MIMOSCA [16], used in [22, 24, 36, 68], SCEPTRE [4, 5], and scMAGeCK [74] (reviewed in [12]). When biological replicates are available, DE is often computed after sample-level pseudobulk aggregation [63] using bulk RNA-seq methods such as DESeq2, edgeR, and limma [10, 43, 55]. Notably, classifier-based approaches are also used to identify perturbation-associated genes, where a model is trained to predict perturbation identity from expression and genes are ranked by their contribution to predictive performance [1, 54].

Another important design choice is which covariates are adjusted for when running DE analyses. These adjustments can change the inferred DEG set, and therefore any downstream DEG-based evaluation metric, by absorbing technical heterogeneity and baseline cellular-state variation before testing. For example, Feng et al. [19] estimate perturbation effects with a linear model that adjusts for cell line, inlet/batch, mitochondrial RNA percentage, cell-cycle scores, and total UMI count per cell. Different choices of adjustment variables can shift DEG calls by removing different combinations of technical variation, baseline biological structure, and perturbation-efficiency differences. Adjustment can also remove real signal: if a covariate such as cell-cycle state is itself altered by the perturbation, controlling for it can mask part of the true response, so that a perturbation whose main effect is to change cell-cycle progression may show a much weaker DE signal after adjustment. The choice of covariates therefore affects not only calibration, but also which biological effects the analysis is able to detect at all.

Several caveats apply to all DEG-based evaluation strategies. First, there is no consensus on which DE testing framework should be used in typical single-cell perturbation assays, and different frameworks can yield very different results (Supplementary Figure 2). In standard single-cell data, DE is typically performed after sample-level pseudobulk aggregation [63] using bulk RNA-seq methods [10, 43, 55]. However, biological replicates are rarely available in current single-cell perturbation studies, so analyses commonly rely on classical two-sample tests at the single-cell level, such as the Wilcoxon rank-sum test [59] or *t*-test [46], which are frequently miscalibrated at this level. For example, Barry et al. [4] compared cells carrying one non-targeting guide with cells carrying another non-targeting guide, a setting in which no true differential expression is expected; well-calibrated methods should yield few significant discoveries after multiple-testing correction, but many commonly used methods produced substantial numbers of false positives.

Second, DEG-based metrics can be difficult to compare across perturbations when DEG sets are defined by statistical significance alone. Perturbations with more cells may yield larger DEG sets at the same effect size simply because they have higher power [49]. Conversely, perturbations with weak effects or few cells may yield very small or even empty DEG sets, which can make DEG-restricted metrics highly variable or ill-defined for those perturbations. Both effects are less problematic when interpreting a single perturbation in isolation, but they can bias aggregate benchmark summaries across perturbations. Possible mitigation strategies include defining DE targets under a fixed effective sample size, for example by subsampling, or using effect-size criteria in addition to or instead of significance thresholds; the latter reduces dependence on sample size but moves DEG-based evaluation closer to delta-based metrics.

Finally, DEG-based metrics depend on the thresholding or ranking strategy used to define candidate genes. Common choices include calling DEGs by an adjusted *p*-value threshold and ranking significant genes by absolute log-fold change [2], or combining a significance threshold with a minimum absolute log-fold-change threshold [78]. Because these choices can substantially alter the resulting scores, we recommend that benchmark reports state the DE method, reference population, covariates, multiple-testing correction, thresholds, ranking score, and resulting DEG-set sizes.

### C.5 Primer on Incorporating Prior Knowledge

Prior biological knowledge, such as curated pathways, gene sets, or gene regulatory networks, can enter evaluation in several ways: as a feature-selection or weighting criterion, as an enrichment-derived embedding (Section 2.3), or as a space in which differential-expression outputs are compared (Section 3.5). Such representations shift the focus from individual genes to interpretable biological modules and can denoise expression profiles by aggregating weak signals across related genes [30]. However, they inherit important limitations of the underlying gene-set collections.

First, biological databases are subject to strong research bias, with a disproportionate fraction of annotations concentrated on a relatively small set of well-studied genes and pathways. Many pathway annotations and expression signatures are also derived from heterogeneous tissues and experimental contexts, and may therefore not reflect the specific cell type or perturbation setting under study [60]. As a result, enrichment-based evaluations may preferentially reward agreement with well-characterized biology while failing to capture context-specific or previously uncharacterized mechanisms.

Second, gene sets are often highly overlapping and hierarchically related, so enrichment analyses may yield multiple closely related terms driven by largely identical gene subsets [53]. If overlap-based metrics are computed directly on such redundant collections, agreement between predicted and observed enrichments can be artificially inflated. Identifying or collapsing redundant gene sets prior to evaluation can help obtain less biased performance estimates.

## D Supplementary Methods and Metric Definitions

### D.1 Notation

Let *A* denote the set of perturbations, with *N*_*A*_ := |A| the total number of perturbations. For each perturbation *a* ∈ *A*, we consider an observed cell population *X*^(*a*,obs)^, a predicted cell population *X*^(*a*,pred)^, and a shared reference population *X* ^(ref)^:

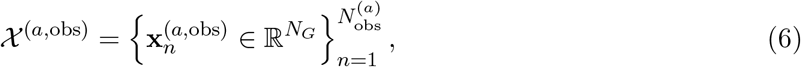

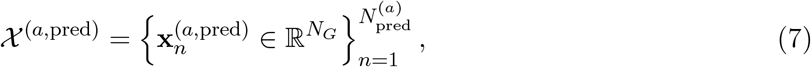

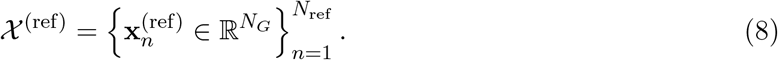

Each cell is represented by an *N*_*G*_-dimensional vector, indexed by features *g* = 1, …, *N*_*G*_. The feature set is denoted by *G* := {1, …, *N*_*G*_}. The value of feature *g* in cell *n* from population *X*^(*a*,·)^ is denoted by 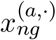.

Unless multiple cell populations appear in the same expression, we use *n* as the cell index. When two populations appear together, we use *n* and *n*^′^ to distinguish them.

Define the centroid (or pseudobulk) vector by

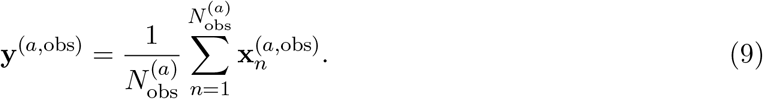

Analogous definitions apply to **y**^(*a*,pred)^ and **y**^(ref)^.

The feature-level components of the centroid are denoted by 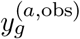, with analogous definitions for 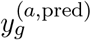 and 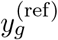. The mean across features is

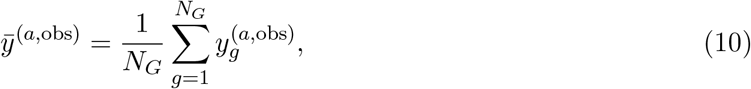

with analogous definitions for 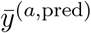 and 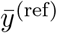.

We further define the empirical probability measure

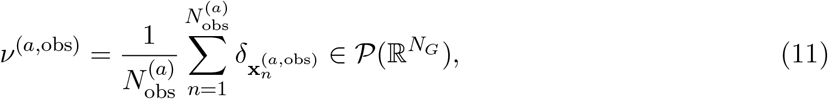

where 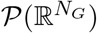 denotes the set of probability measures on 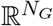. Analogous definitions apply to _*v*_(*a*,pred) _and *v*_(ref).

Reference-based centroid shifts are written as

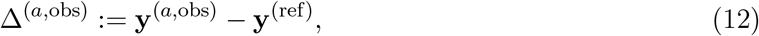

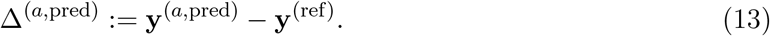

When a perturbation *a* is fixed, we use the local shorthand, dropping the superscript *a*:

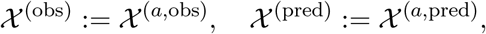

and analogously for **y**^(obs)^, **y**^(pred)^, *v*^(obs)^, *v*^(pred)^, Δ^(obs)^, Δ^(pred)^, and feature-level quantities such as 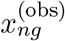 and 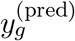. In this fixed-*a* setting, we write *N*_obs_: = |*X* ^(obs)^| and *N*_pred:_ = |*X*^(pred)^|.

We use this shorthand only in sections that define a metric for one fixed perturbation; perturbation-indexed objects and cross-perturbation comparisons retain explicit superscript *a*.

The indicator function is denoted by **1**[·], whereas **1** without an argument denotes an all-ones vector whose dimension is inferred from context.

We use *ϕ* to denote a benchmark-dependent representation map that sends the available object (e.g. single-cell data) to the representation actually compared, for example a centroid (pseudobulk or average-expression) vector, a population of single cells, or an embedding-derived representation. We use d to denote a metric applied to such representations. For notational convenience, we define the composition

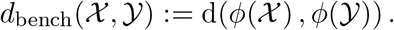

Here, *d*_bench_ denotes the benchmark-specific metric induced by the chosen representation map *ϕ* and metric d.

For notational convenience, we write*X*^(*a*,pred)^ as if models produced a predicted cell population. This should be understood as an abstraction: in practice, model outputs may take different forms, including generated cells, direct centroid predictions, or lower-dimensional representations. The relevant object for evaluation is therefore *ϕ*(*X*^(*a*,pred)^), interpreted as the benchmark-specific representation induced by the model output, regardless of whether a full predicted population exists.

We use the term metric loosely to include distances, divergences, similarities, and test statistics.

Unless stated otherwise, all metric definitions are written in the ambient feature space 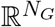. Some metrics may be computed in a lower-dimensional embedding with effective dimension *N*_emb_ ≪ *N*_*G*_, as discussed in Section 2.3, but we retain the *N*_*G*_-based notation throughout for consistency.

General notation used throughout the manuscript.

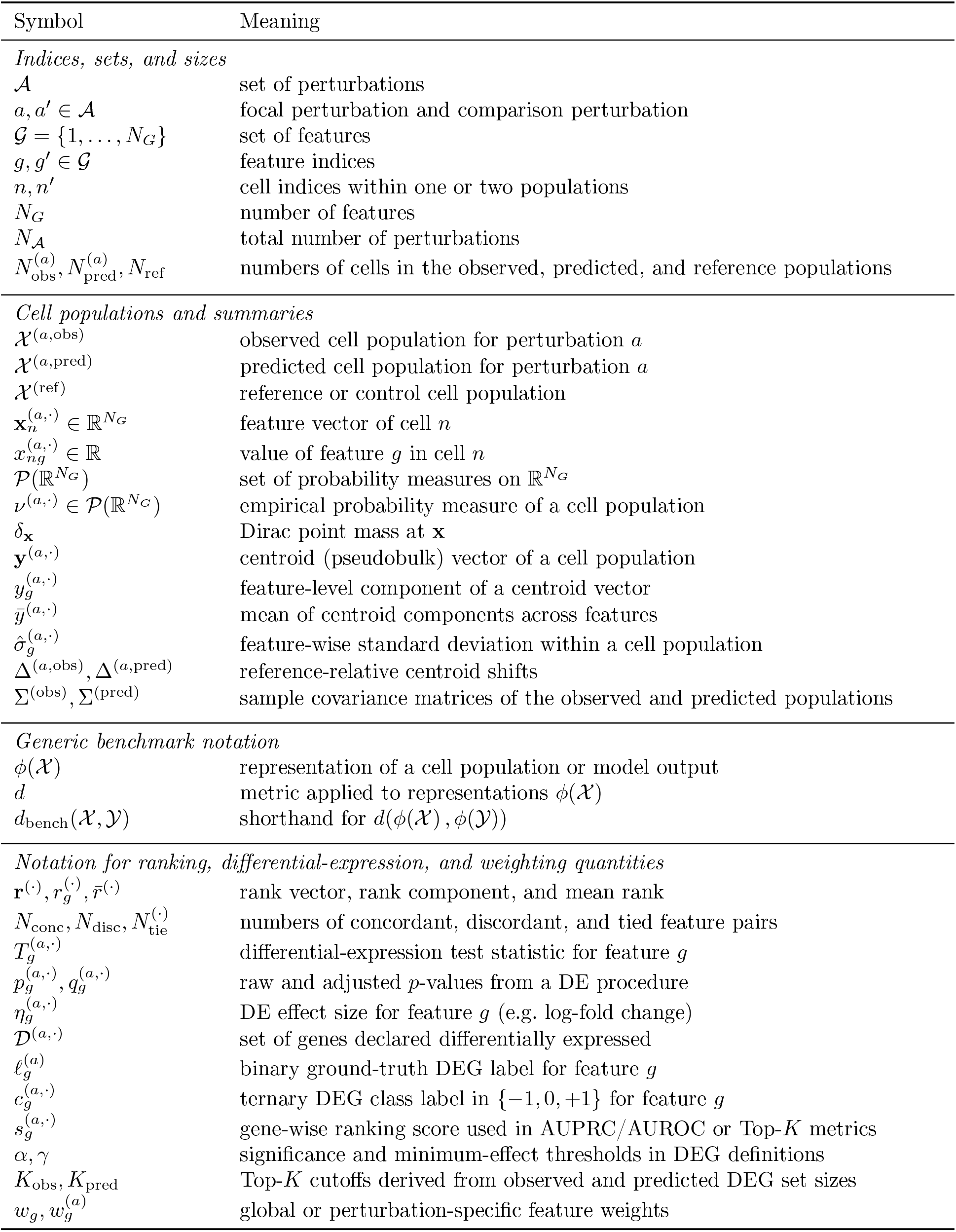

### D.2 Centroid Metrics

Centroid metrics compare observed and predicted perturbation responses after each cell population has been summarized by a centroid, or pseudobulk, vector. The same definitions can be applied to reference-centered centroid shifts by replacing **y**^(obs)^ and **y**^(pred)^ with Δ^(obs)^ and Δ^(pred)^. In this section, all metrics are defined for a fixed perturbation, using the local shorthand **y**^(obs)^ and **y**^(pred)^ introduced in Supplementary Section D.1.

#### D.2.1 Pearson Correlation

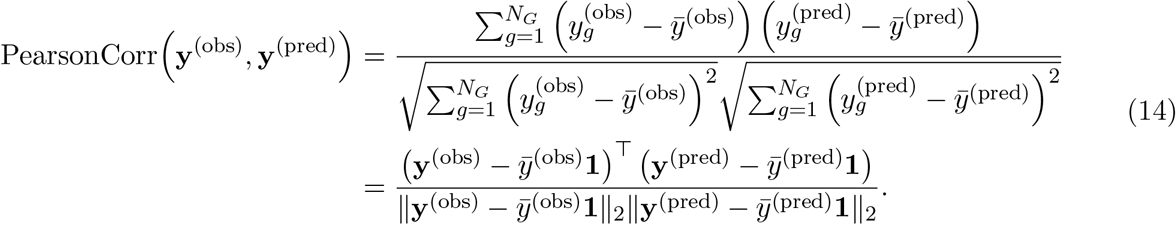

The corresponding Pearson distance is defined as

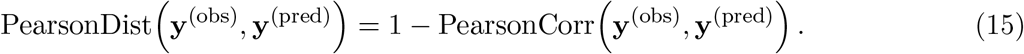

Pearson correlation is undefined if either centroid vector is constant across features.

#### D.2.2 Cosine Similarity

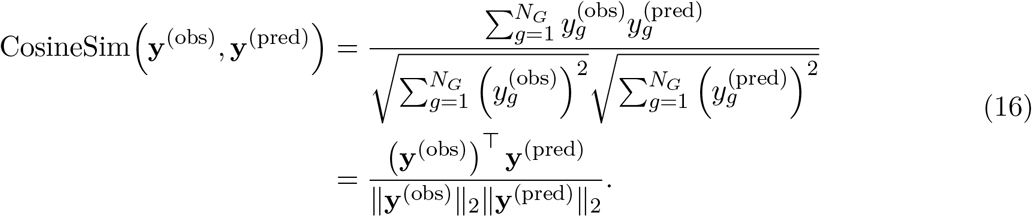

The corresponding cosine distance is defined as

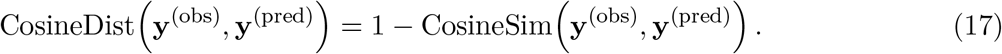

Cosine similarity is undefined if either centroid vector has zero Euclidean norm.

#### D.2.3 Spearman Correlation

Let rank_*S*_(*x*) denote the rank of *x* within a finite set *S*, with ties assigned average ranks. For Spearman correlation, define the rank vectors

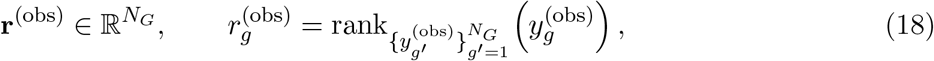

and analogously for **r**^(pred)^. We further define

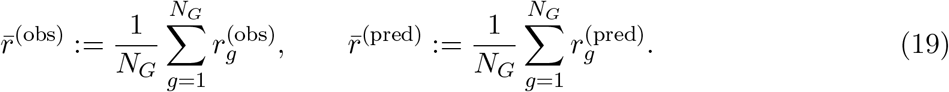

The Spearman correlation is defined as the Pearson correlation between the rank vectors:

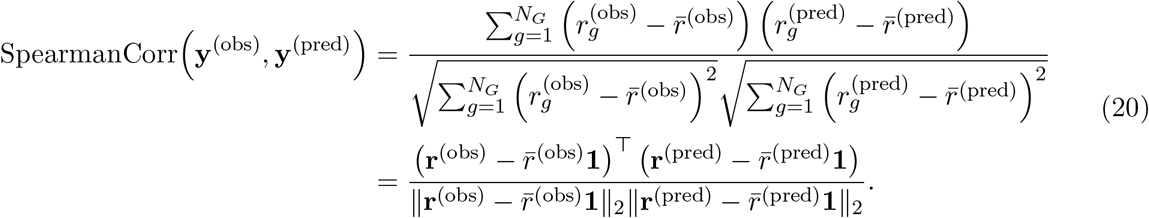

The corresponding Spearman distance is defined as

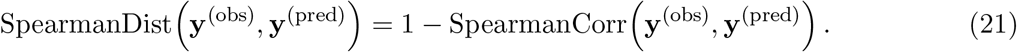

Spearman correlation is undefined if either rank vector is constant.

#### D.2.4 Kendall’s Tau

Kendall’s *τ* is a rank-based measure of association that quantifies agreement in the pairwise ordering of features between two vectors. Unlike Spearman’s correlation, which computes a correlation between rank-transformed vectors, Kendall’s *τ* is defined directly through concordant and discordant feature pairs. It therefore measures a stricter form of ordinal agreement.

∈

Given centroid vectors **y**^(obs)^, 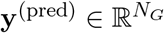, consider all feature pairs (*g, g*^′^) with 1 ≤ *g < g*^′^ ≤ *N*_*G*_. A pair is concordant if

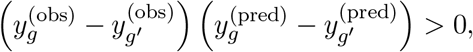

discordant if this product is negative, and tied if either difference is zero. Let *N*_conc_ and *N*_disc_ denote the numbers of concordant and discordant pairs. In the absence of ties, Kendall’s *τ* is

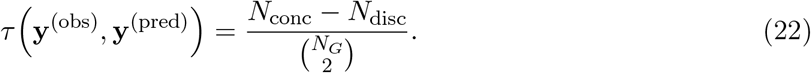

Because ties can occur in sparse or discretized expression representations, a common tie-corrected variant is Kendall’s *τ*_*b*_,

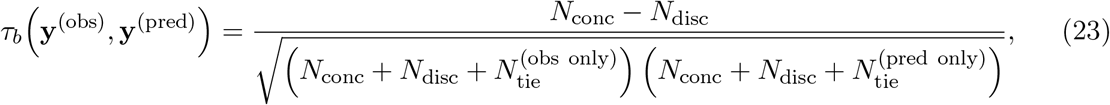

where 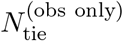 and 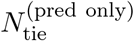 denote the numbers of pairs tied only in the observed or predicted vector, respectively. Pairs tied in both vectors contribute neither to the numerator nor to either one-sided tie count.

Analogously to other correlation-based measures, we define the Kendall distance as

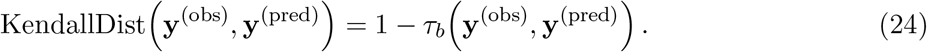

Kendall’s *τ*_*b*_ ranges from −1 to 1, where 1 indicates identical pairwise ordering and −1 indicates complete reversal. Like other rank-based measures, it is invariant under strictly monotonic transformations, but its direct pairwise definition leads to a naive computational cost of 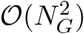.

#### D.2.5 L1 Distance & Mean Absolute Error

The L1 distance is defined as

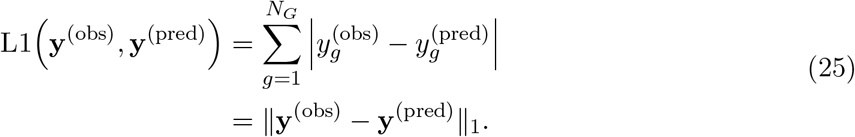

The mean absolute error (MAE) is defined as

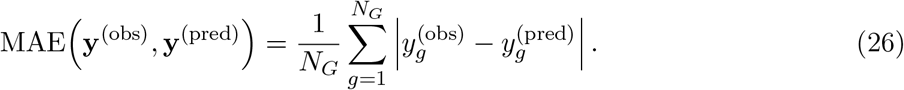

#### D.2.6 L2 Distance & Mean Squared Error

The L2 distance is defined as

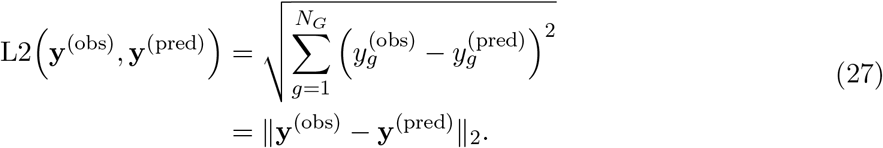

The mean squared error (MSE) is defined as

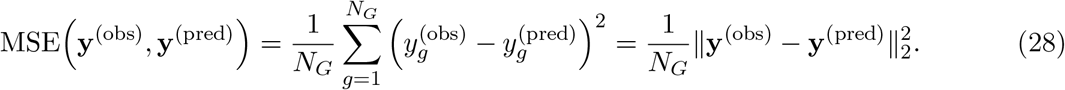

#### D.2.7 Root Mean Squared Error (RMSE)

The root mean squared error (RMSE) is defined as

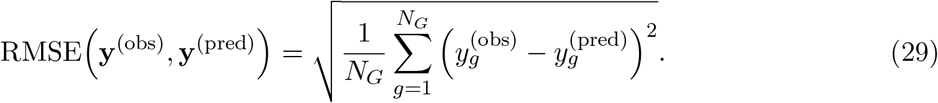

#### D.2.8 Mahalanobis Distance

The Mahalanobis distance is a covariance-weighted centroid distance. 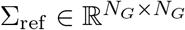 denote a shared positive-definite reference covariance matrix estimated from observed data only, for example from control cells, training cells, or another shared background population. The Mahalanobis distance between the observed and predicted centroids is then

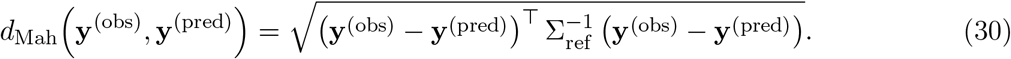

Using a shared covariance estimate keeps the metric compatible with pseudobulk-only predictions, because the covariance does not need to be estimated separately for each target perturbation or from the prediction itself. Intuitively, the metric whitens the mean difference before taking its Euclidean norm, thereby downweighting directions with large background variance and accounting for correlated variation across genes. Because the same reference covariance is used for every comparison, this definition is symmetric in the two centroids.

At the same time, Mahalanobis distance remains a centroid-based metric: it reweights mean differences but does not compare the covariance structures of the observed and predicted populations. It should therefore be distinguished from joint distribution metrics such as the Fréchet distance, which compare both mean and covariance summaries.

In single-cell gene space, classical Mahalanobis distance is often difficult to use directly because the sample covariance is typically rank-deficient or poorly conditioned when the number of cells is smaller than the number of genes. In practice, one usually needs dimensionality reduction, covariance shrinkage or other regularization, a restricted feature space, or a pseudoinverse 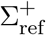 instead of 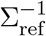. These choices should be reported because they define the effective metric being used.

PertPy implements a related directional variant in which the covariance matrix is estimated from the first cell-level input rather than from a fixed shared reference population [28]. This differs from the symmetric shared-reference definition above and requires cell-level observations in the first argument. For reproducible symmetric comparisons, Σ_ref_ must be fixed independently of the argument order, for example using a pooled or control-reference covariance estimate.

#### D.2.9 Coefficient of Determination (*R*^2^)

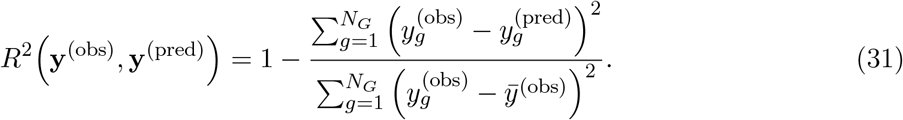

The coefficient of determination *R*^2^ quantifies the extent to which the predictions **y**^(pred)^ explain the variance in the observed centroid **y**^(obs)^. Equivalently, it measures the relative improvement over a trivial baseline that predicts the same global mean 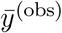 for all features. Values below zero indicate worse squared error than this baseline.

### D.3 Marginal Distribution Metrics

Marginal distribution metrics compare observed and predicted cell populations feature by feature, and then aggregate the resulting feature-wise discrepancies across genes. Because these metrics are constructed from feature-wise discrepancies, feature weighting can usually be implemented by replacing the unweighted average over genes with a weighted average.

#### D.3.1 Poisson Negative Log-Likelihood

Count likelihoods can be used as marginal distribution metrics when they are evaluated on cell-level gene marginals rather than on pseudobulk means. For each feature *g*, one fits a Poisson model to the predicted marginal distribution and evaluates the observed counts under that fitted model. Under a Poisson model, let 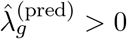 denote the feature-wise rate estimated from the predicted cells. The directional Poisson negative log-likelihood is defined as

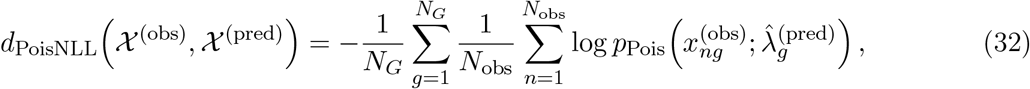

Where

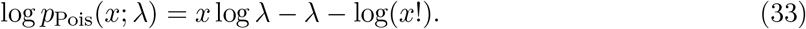

This score compares each gene marginal separately and then averages across genes. It therefore captures mean-dependent count variance under the Poisson assumption, but it ignores cross-gene dependence and is most appropriate for raw counts with appropriate handling of cell-specific library size or exposure.

#### D.3.2 Negative Binomial Negative Log-Likelihood

The negative-binomial analogue replaces the Poisson marginal with an overdispersed count model. For each feature *g*, let 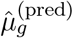 and 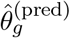 denote the mean and size parameters fitted from the predicted cells, with

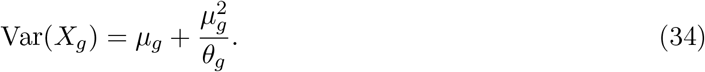

The directional negative-binomial negative log-likelihood is defined as

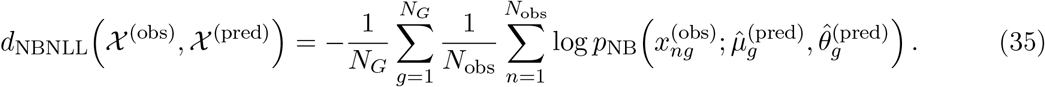

Compared with the Poisson version, this score accounts for feature-wise overdispersion. Like other marginal distribution metrics, however, it treats genes independently. The score is also directional because one population supplies the fitted marginal distributions and the other is evaluated under them. The `nb_ll` metric in [28] follows this gene-wise likelihood logic, fitting a negative-binomial model for each feature in one group and returning the average negative log-likelihood of the other group under those fitted marginals.

#### D.3.3 Welch’s T-Statistic (Two-Sample T-Test)

Given some feature *g*, the unbiased sample variances are defined as

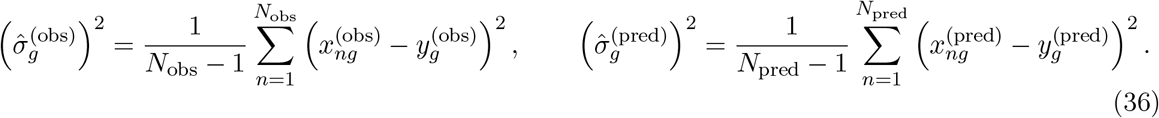

Then Welch’s test statistic is defined as

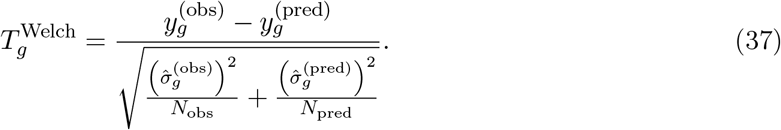

Under the null hypothesis of equal means, the statistic follows a Student-*t* distribution with degrees of freedom approximated by the Welch–Satterthwaite formula.

To obtain a single scalar metric score for the population comparison, one may average the absolute feature-wise statistics across genes,

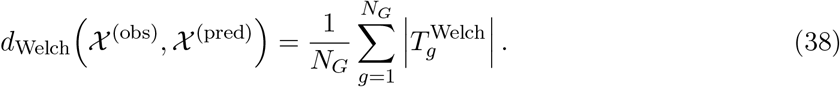

This matches the implementation used in [28, 31].

#### D.3.4 Rank-Sum (Wilcoxon–Mann–Whitney Test)

Given some feature *g*, we form the pooled set

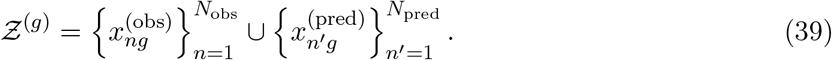

Let rank 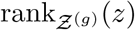 denote the rank of *z* among *Ƶ*^(*g*)^ (with average ranks for ties). The rank-sum statistic for the observed sample is defined as

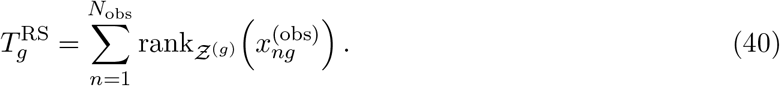

The Mann–Whitney *U* statistic is defined as

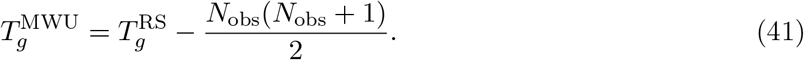

Under the null hypothesis of equal distributions and in the absence of ties, the mean of the null distribution is

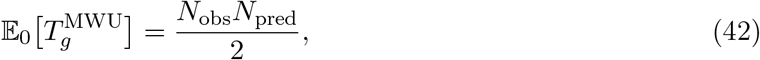

and the variance is

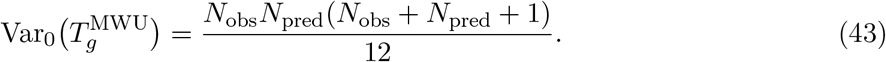

In the presence of ties, the null variance is adjusted by the usual tie-correction factor.

To obtain a single scalar metric score for the population comparison, one may standardize the feature-wise Mann–Whitney statistics under the null and average their absolute values across genes,

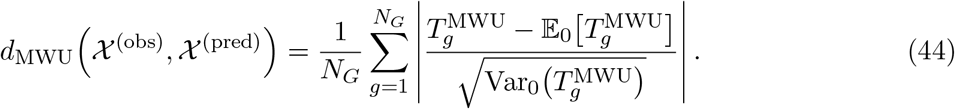

#### D.3.5 Kolmogorov–Smirnov (KS) Statistic

For each feature *g*, we define the empirical cumulative distribution functions (ECDFs)

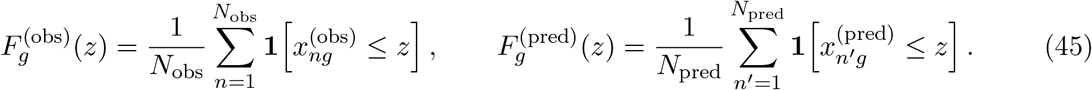

The KS statistic for feature *g* is defined as the maximum absolute deviation between the two ECDFs,

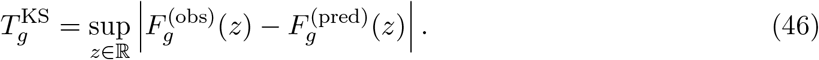

To obtain a single scalar metric score for the population comparison, one may average these feature-wise statistics across genes,

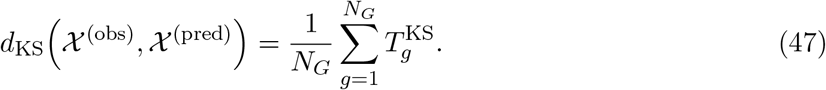

This matches the aggregation used in [28, 31].

#### D.3.6 Symmetric KL Divergence on Gaussian Summaries

Assume that, for each feature *g*, the observed and predicted marginals are approximated by marginal Gaussians with means 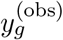 and 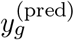 and standard deviations 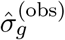 and 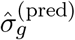, respectively. The directional feature-wise Kullback–Leibler divergence from the observed distribution to the predicted distribution is

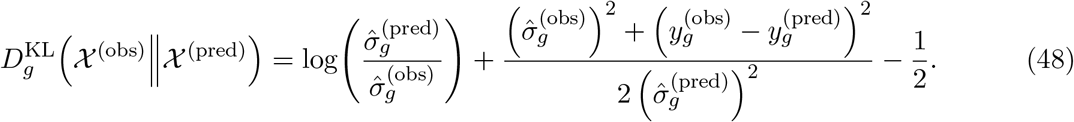

The benchmarked metric used in [28, 31] is the symmetrized variant obtained by averaging both KL directions across genes,

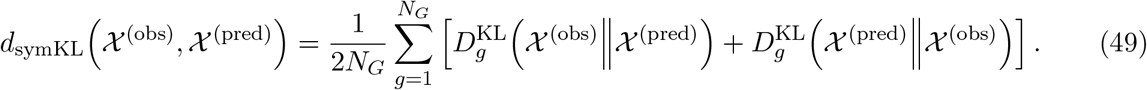

This formulation compares the first two moments of the feature-wise marginals under an independent Gaussian approximation. It therefore ignores cross-feature dependence and any departures from Gaussian shape beyond mean and variance.

Because the directional KL divergence is asymmetric, symmetrization removes the dependence on which population is assigned to the left or right argument while retaining the same Gaussian moment assumptions.

This metric is implemented in [28], benchmarked in [31], and used in [71].

### D.4 Joint Distribution Metrics

Joint distribution metrics compare the full empirical distributions of observed and predicted cell profiles in the chosen feature or embedding space. Unlike centroid metrics, they can capture differences in population geometry beyond mean shifts, including changes in covariance structure, multimodality, or nonlinear dependencies. In this section, all metrics are defined for a fixed perturbation, using the local shorthand *v*^(obs)^ and *v*^(pred)^ introduced in Supplementary Section D.1.

#### D.4.1 Maximum-Mean Discrepancy (MMD)

Maximum-mean discrepancy (MMD) is a kernel-based distance between probability distributions [25]. An important way to understand it is through the kernel feature-map perspective. If *k* is a positive-definite kernel, then there exists a feature map *ψ* into a reproducing kernel Hilbert space (RKHS) ℋ such that

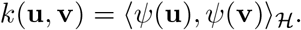

MMD can then be written as the distance between the mean feature embeddings of the two distributions,

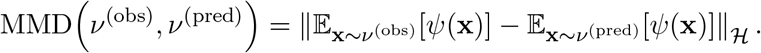

Thus, MMD compares distributions after mapping the data through a potentially very rich collection of nonlinear features. The choice of kernel determines which aspects of the joint distribution are emphasized, allowing MMD to capture differences beyond simple mean shifts.

From the perspective of Section 2, the kernel therefore induces an implicit representation space: MMD compares distributions after mapping cells into the RKHS defined by the kernel, rather than after an explicit embedding such as PCA or a pretrained latent model.

In practice, this population quantity is not available directly and must be estimated from samples through kernel evaluations between observed and predicted cells. For a positive-definite kernel 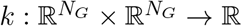, the squared MMD between the empirical distributions *v*^(obs)^ and *v*^(pred)^ can be estimated in different ways [25].

##### Biased estimator

The biased empirical estimator is

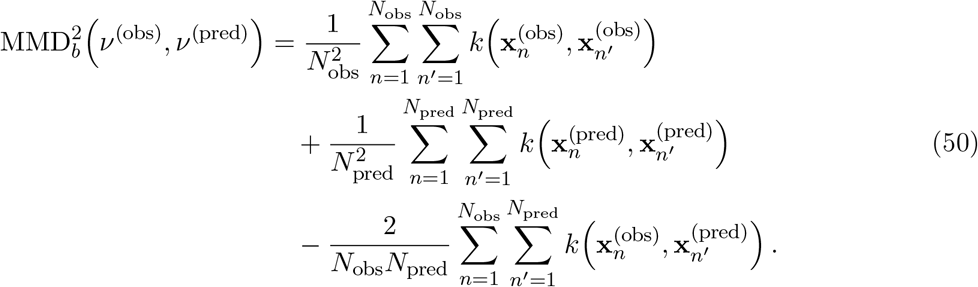

##### Unbiased estimator

The unbiased empirical estimator excludes the diagonal terms:

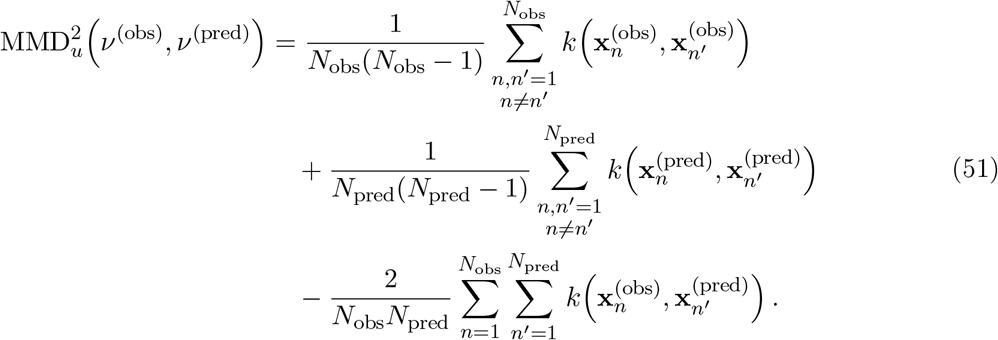

The biased estimator is always nonnegative but has a finite-sample upward bias under the null. The unbiased estimator removes this bias, but can take negative values in finite samples even though the population quantity is nonnegative.

A common choice is the radial basis function (RBF) kernel, also known as the squared exponential kernel,

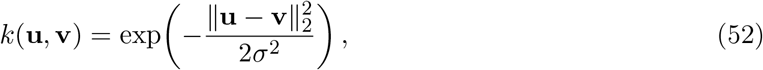

where *σ >* 0 denotes the kernel bandwidth. With this choice, nearby cells contribute more strongly to one another’s similarity, so the resulting discrepancy becomes sensitive to broad differences in population geometry rather than only linear moment differences.

In practice, MMD^2^ is often computed for several bandwidths 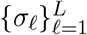, and the resulting values are averaged across bandwidths [8, 18, 35]. Another option is the inverse multiquadratic kernel,

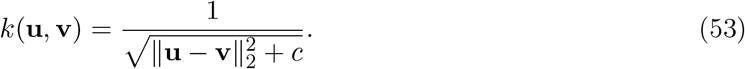

More generally, MMD naturally extends beyond vector-valued data: as long as a suitable kernel can be defined, it can also be applied to more general objects such as strings [7]. Compared with many other joint distribution metrics, MMD is relatively inexpensive to compute and can capture nonlinear discrepancies through the kernel. However, its behavior depends strongly on the chosen kernel and associated hyperparameters, especially the bandwidth, and the resulting score is typically less directly interpretable than a centroid metric. MMD is used, for example, in [18].

#### D.4.2 Energy Distance

The energy distance [57] is a related joint distribution metric. Under the Rizzo convention, the population quantity

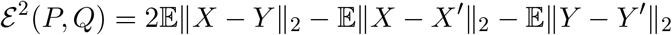

is the squared energy distance, with *X, X*^′^ ∼ *P* and *Y, Y* ^′^ ∼ *Q*. The energy distance itself is 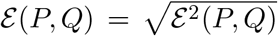. Using all-pairs empirical averages, the corresponding empirical squared energy distance is

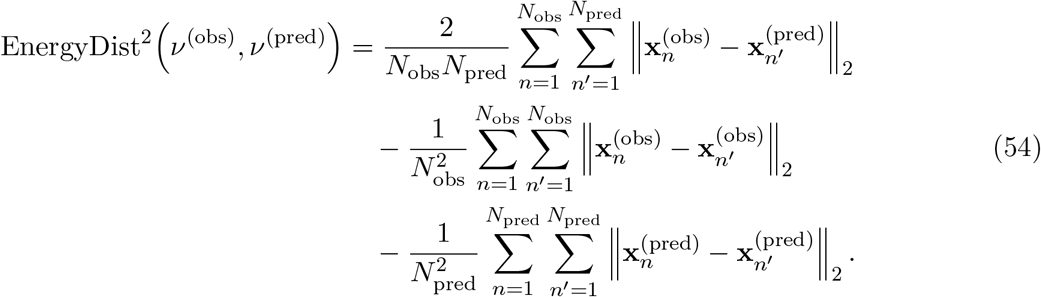

Some single-cell papers refer to this displayed unsquare-rooted statistic as “E-distance”; benchmark reports should therefore state explicitly whether they report EnergyDist^2^ or EnergyDist.

The first term measures average cross-population distances, whereas the second and third terms subtract average within-population distances. The energy distance is therefore small when the two populations are difficult to distinguish by their pairwise distance structure, and it increases when cross-population distances are large relative to within-population distances.

Energy distance is used in Peidli et al. [52], where it is computed in PCA space, with perturbation groups subsampled to equal cell counts within each dataset because empirical energy-distance estimates can vary with sample size.

The signature distance [38] compares the full distribution of distances from each reference point to the within-set and cross-set populations, rather than reducing these distances to a single mean as in the energy distance. For a reference point, these profiles are represented by sorted within-set and cross-set distance-profile vectors:

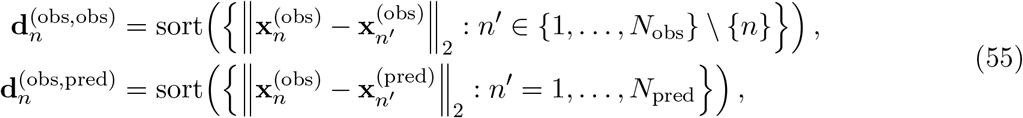

and analogously for 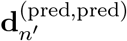 and 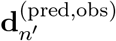. The pointwise discrepancy is the one-dimensional Wasserstein distance between the empirical distributions induced by these distance-profile vectors, which for equal-length sorted arrays reduces to the mean absolute difference of aligned entries. The resulting squared signature distance is

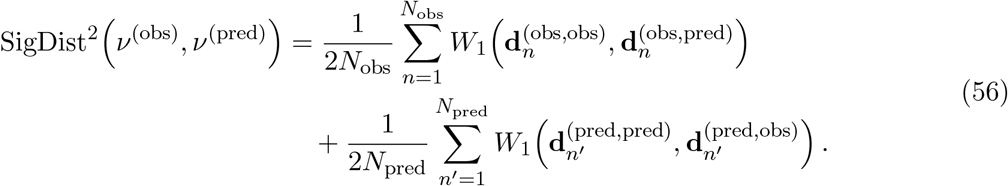

Relative to energy distance, signature distance retains the shape of each point’s radial distance distribution and can therefore detect density and geometric differences that are invisible to a pure mean-distance reduction, while remaining in the same *O* (*N* ^2^) complexity class up to the one-dimensional profile comparisons.

#### D.4.3 Wasserstein Distance and Entropic Transport Cost

Using the Kantorovich formulation, the optimal transport cost between the empirical measures *v*^(obs)^ and *v*^(pred)^ is defined as

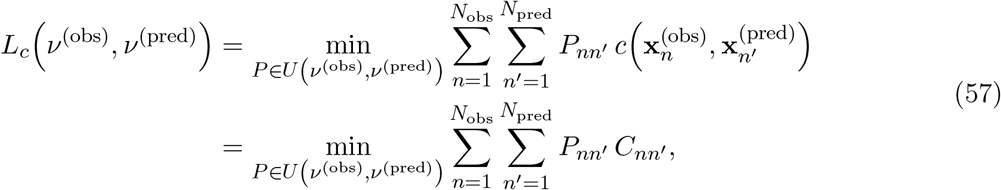

where 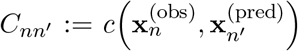 and 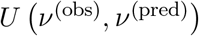 denotes the set of discrete couplings with prescribed marginals:

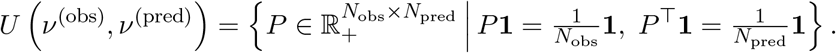

Here, 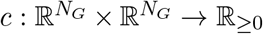 is the cost function determining how expensive it is to move probability mass from any 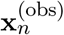 to any 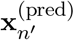.

This optimization is a linear program with *N*_obs_*N*_pred_ variables and *N*_obs_ + *N*_pred_ equality constraints, and therefore does not scale well in the number of cells. If

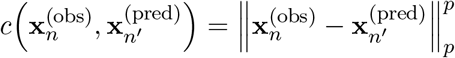

for *p* ≥ 1, then the induced quantity 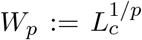 defines the family of Wasserstein *p*-distances, which are symmetric, positive definite, and satisfy the triangle inequality. In particular, choosing 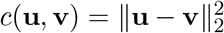 yields the *W*_2_ distance.

To improve computational efficiency, one may introduce entropic regularization [15]:

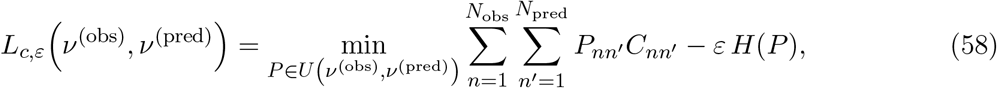

Where

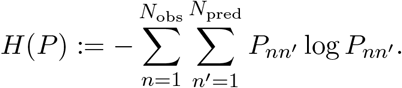

This problem can be solved efficiently using Sinkhorn’s algorithm. For the quadratic cost 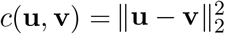,we denote the corresponding regularized transport cost by

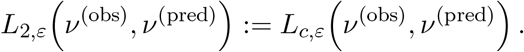

Thus, *L*_2,*ε*_ denotes the entropically regularized quadratic transport cost.

Entropic regularization improves computational tractability and yields smoother optimization problems. However, the regularized transport cost is biased by the entropy term: for *ε >* 0, the identity

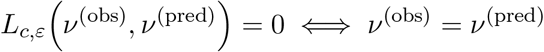

does not generally hold. Entropically regularized transport costs are used in [35, 50].

#### D.4.4 Sinkhorn Divergence

By subtracting self-interactions from the entropically regularized quadratic transport cost, Genevay, Peyre, and Cuturi [23] and Feydy et al. [20] introduced the Sinkhorn divergence:

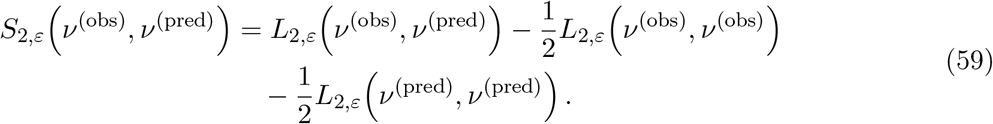

This correction ensures that *S*_2,*ε*_ *v*^(obs)^, *v*^(pred)^ = 0 if and only if *v*^(obs)^ = *v*^(pred)^ (see Theorem 1 in [20]), thereby recovering the separation property that is lost in the entropically regularized transport cost for *ε >* 0. Sinkhorn divergence is used in [35, 50].

#### D.4.5 Fréchet Distance

Another joint distribution comparison strategy is to approximate each population by a Gaussian summary and then compare these summaries. For a fixed perturbation, let

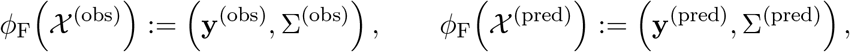

where Σ^(obs)^ and Σ^(pred)^ denote the sample covariance matrices of the two populations, e.g.

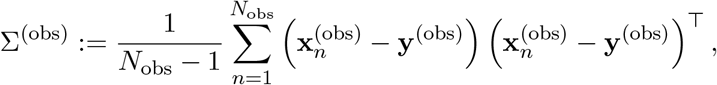

and analogously for Σ^(pred)^.

The squared Fréchet distance between these Gaussian summaries is

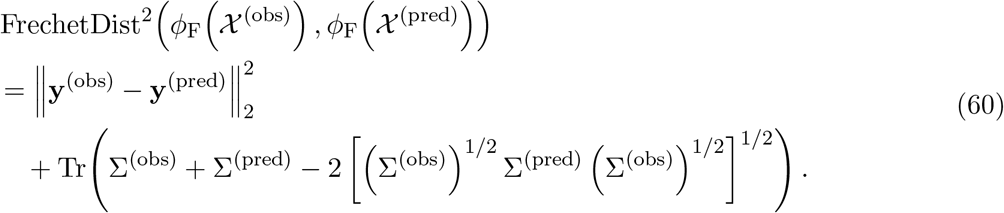

If the final square root of this quantity is not taken, the reported value should be described as the squared Fréchet distance.

This metric compares the first and second moments of Gaussian approximations to the two populations and therefore depends on how well such summaries capture the relevant biological variation. Conceptually, the squared Fréchet distance is the squared 2-Wasserstein distance between Gaussian approximations of the two populations, and may therefore be viewed as a moment-based approximation to full optimal transport.

Unlike centroid-level Mahalanobis distance, which whitens the mean difference using a shared reference covariance, the Fréchet distance compares both mean and covariance summaries. It does not require covariance inversion, and positive-semidefinite singular covariance matrices still possess positive-semidefinite matrix square roots. Rank deficiency is therefore not, by itself, a mathematical reason for the Fréchet distance to be undefined. In high-dimensional finite-sample settings, however, rank deficiency can cause numerical instability in matrix-square-root implementations. Eigenvalue clipping, covariance shrinkage, a small diagonal regularizer, or dimensionality reduction can be used for numerical robustness, but these choices alter the numerical estimator and should be reported.

The most common use of this construction is the Fréchet Inception Distance (FID) from computer vision [29], where real and generated images are passed through a fixed pretrained Inception-v3 network [65], the activations of the final pooling layer are summarized by their mean and covariance, and the Fréchet distance between these Gaussian feature distributions is reported. In [56], this idea was transferred to single-cell transcriptomics as single-cell Fréchet Inception Distance (scFID): cells were first embedded with the pretrained single-cell foundation model scGPT [14], then the Fréchet distance was computed between the Gaussian summaries of the real and generated cell embeddings for each unique test condition, defined by the combination of cell type, perturbation, and exposure duration, and finally averaged across conditions.

### D.5 Composition Metrics

When perturbation responses are summarized by proportions over predefined cell groups, the resulting vectors are compositional. Let

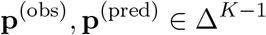

denote the observed and predicted proportions over *K* cell groups, where

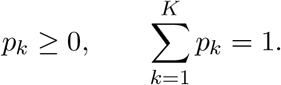

Because the components are constrained to sum to one, they cannot be interpreted as independent features: increasing the proportion of one group necessarily decreases the relative proportions of others. Composition metrics therefore compare vectors on the simplex rather than treating them as unconstrained Euclidean vectors.

A simple metric for composition vectors is total variation distance,

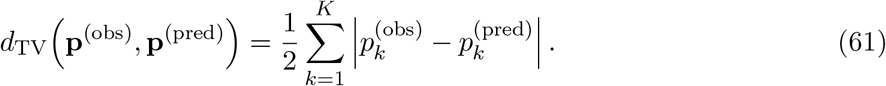

Total variation distance ranges from 0 to 1 and measures the largest discrepancy that the two compositions assign to any subset of cell groups. It is easy to interpret and directly quantifies how much probability mass must be reassigned to transform one composition into the other.

Another common choice is Jensen–Shannon divergence. Let

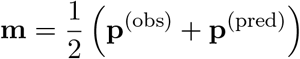

denote the midpoint composition. Then

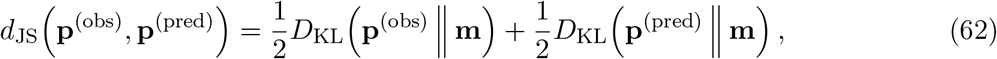

Where

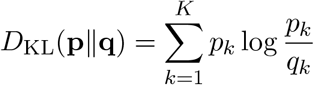

is the Kullback–Leibler divergence. Jensen–Shannon divergence is symmetric and finite whenever the two compositions have zeros in different entries. Its square root is a metric.

A third option is Aitchison distance, which compares compositions in log-ratio geometry. Using the centered log-ratio transformation,

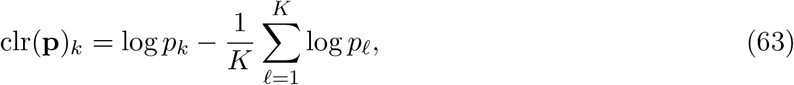

the Aitchison distance is

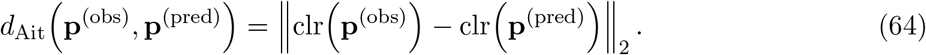

This metric is invariant to the absolute scale of the underlying positive abundances before closure to proportions and is therefore natural when only relative composition matters. However, because the logarithm is undefined for zero entries, practical use requires an explicit zero-handling strategy, such as pseudocount addition or zero replacement before applying the centered log-ratio transformation.

These metrics emphasize different aspects of composition error. Total variation has the most direct probability-mass interpretation, Jensen–Shannon divergence compares the two compositions as discrete probability distributions, and Aitchison distance uses the relative log-ratio geometry of compositional data. In perturbation-response benchmarks, the chosen metric should therefore reflect whether the goal is to measure absolute changes in cell-group proportions, distributional discrepancy over discrete groups, or relative compositional shifts.

### D.6 Differential Expression Metrics

Differential-expression metrics compare observed and predicted perturbation responses after both have been converted into gene-wise DE summaries relative to a shared reference population. For each perturbation *a*, differential expression is computed twice: once by comparing the ground-truth perturbed cells *X*^*(a,obs)*^ *to t*he reference population *X* ^*(ref)*^, *and o*nce by comparing the predicted cells *X*^*(a,pred)*^ *to* the same reference population.

For each gene *g* ∈ *G, let*

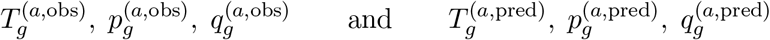

*denote the* resulting DE test statistic, raw *p*-value, and adjusted *p*-value obtained from these two comparisons, respectively. Likewise, let

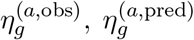

denote effect-size estimates (e.g. log-fold changes) for the ground-truth and predicted data.

A DEG-based metric then compares the DE evidence vectors across genes, either via set agreement of declared DEGs, ranking agreement of DE evidence scores, directionality agreement, or continuous comparison of DE-derived scores. In many benchmarks, direction is ignored and only magnitude is considered, e.g. via 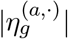 or 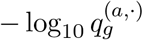.

To define the observed DEG set, we need to define a rule that specifies which genes are considered DE for perturbation *a*, for example

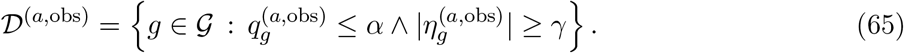

Analogously define *D*^*(a,pred)*^ *fro*m 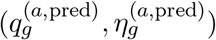.

#### D.6.1 AUPRC from Predicted DE Evidence

Define binary DE labels based on the observed DEG set,

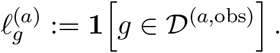

To compute the precision-recall curve, each gene is assigned a prediction score 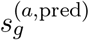 derived from the predicted DE evidence. For example, one may use

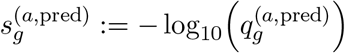

as in [2], where predicted adjusted *p*-values are used as confidence scores. [17] uses the same ranking idea with − log_10_ of predicted *p*-values. Alternatively, one may use

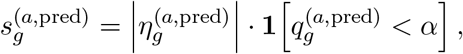

which combines predicted effect size and statistical significance, as proposed in [78]. Genes are ranked by 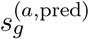 and the area under the precision-recall curve (AUPRC) is computed against the binary labels 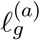.When scores are based on − log_10_-transformed *p*-values or adjusted *p*-values, zero values should be clipped to a positive lower bound before transformation.

#### D.6.2 AUROC from predicted DE evidence

As for AUPRC, define binary DE labels from the observed DEG set,

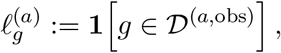

and let 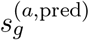 be a prediction score derived from the predicted DE evidence, for example

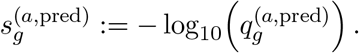

In [2], this score is explicitly the predicted − log_10_ adjusted *p*-value. [70] likewise describes AUROC as a ranking over predicted DE scores and gives log_10_ adjusted *p*-values as the stated score example. The area under the receiver operating characteristic curve (AUROC) is then computed by ranking genes according to 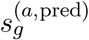 and comparing these rankings against the binary labels 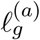 Equivalently, AUROC can be interpreted as the probability that a randomly chosen observed DE gene receives a higher prediction score than a randomly chosen observed non-DE gene.

#### D.6.3 Directionality Agreement

Beyond asking whether the same genes are identified as differentially expressed, one may also ask whether the direction of the predicted perturbation effect agrees with the ground truth. Let

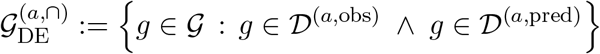

denote the set of genes called significantly DE in both prediction and ground truth for perturbation a.The DE direction match score is then defined as

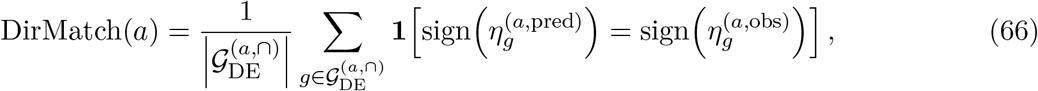

provided that 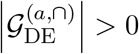.

This score quantifies agreement in the direction of regulation (up- versus down-regulation) among genes that are called DE in both prediction and ground truth. It is used in [2, 70] and called “DE direction match” in [70]. In contrast, in GEARS [58], sign agreement is not evaluated on the overlap of predicted and observed DE calls, but on the top-20 observed DE genes for each perturbation.

#### D.6.4 Macro F1 for DEG classification

An alternative is to treat DEG prediction as a three-class classification problem over down-regulated, unchanged, and up-regulated genes. Using the same thresholds as above, define for each gene *g*∈ *G the class labels*

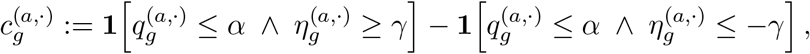

so that 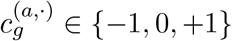 encodes down-regulated, not called DE, and up-regulated genes, respectively. For each class *c* ∈ {−1, 0, +1}, define

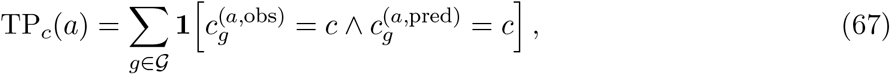

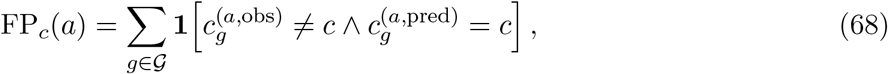

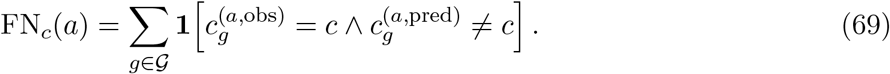

The class-wise precision and recall are then

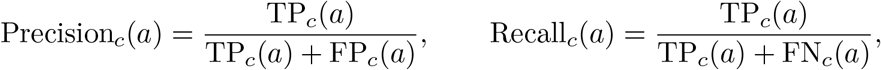

with the usual convention that undefined precision or recall values caused by zero denominators must be handled explicitly, for example by setting the corresponding value to zero or omitting the undefined class according to the benchmark protocol. The corresponding class-wise F1 score is

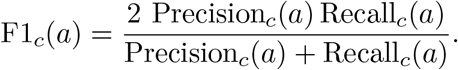

Macro F1 is obtained by averaging these class-wise scores,

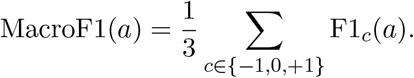

This metric therefore evaluates balanced agreement across down-regulated, unchanged, and upregulated genes, incorporates directionality explicitly, and is less dominated by the majority class than plain accuracy. This three-class DEG formulation is used in [13].

#### D.6.5 Top-*K* Overlap

Let TopK*K* (**s**^(*a*)^) ⊆ *G* denote the set of the *K* genes with largest values of a gene-wise score 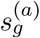 (ties broken deterministically). Common choices are

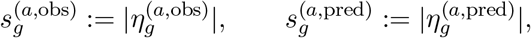

or alternatives such as 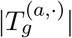 or 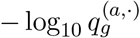. The Top-*K* overlap score is

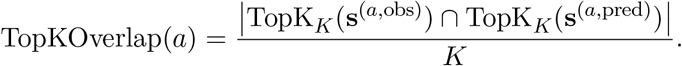

This metric is used in [2, 17], where overlap-based variants are reported as *DE overlap accuracy* when *K* := |D ^(*a*,obs)^| and *DE precision@K* when *K* := |D ^(*a*,pred)^ |. A fixed-*K* variant is also used in [73] with *K* = 20, where it is called *DEG recall*, and in [32, 71] with *K* = 100, where it is called *Common-DEGs*.

≤

When *K* = |D^(*a*,obs)^ |, the metric is recall-like, quantifying the fraction of top ground-truth DE genes recovered among the top predicted genes. When *K* = |D^(*a*,pred)^|, it is precision-like, reflecting the reliability of the predicted DEG list. For [73], the ranking score is based on DEG test statistics from `scanpy.tl.rank_genes_groups`; for [71], the intended ranking is described in the paper as the top 100 genes by absolute effect size. In [59], the differential expression score (DES) is a recall-like asymmetric variant rather than a pure Top-*K* overlap: with *K* := ^(*a*,obs)^, the full predicted DEG set is used when |D^(*a*,pred)^| ≤ *K*, and only when |D^(*a*,pred)^ | *> K* is it truncated to its top-*K* genes ranked by absolute log-fold change.

#### D.6.6 Jaccard Similarity of DEG Sets

For each perturbation *a*, the Jaccard similarity between observed and predicted DEG sets is defined as

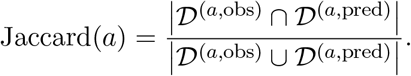

This metric is used in [17]. If both DEG sets are empty, the denominator is zero and the benchmark must define a convention, for example treating the score as perfect agreement or excluding that perturbation from the average.

#### D.6.7 Continuous DE-score comparison

Instead of thresholding differential-expression evidence into DEG sets or class labels, one can compare continuous DE-derived score vectors directly. For example, define a signed significance score

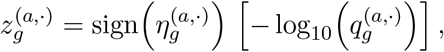

possibly after clipping very small adjusted *p*-values, e.g. 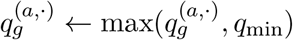, to avoid domination by distinctions among already highly significant genes.

The observed and predicted DE-output vectors can then be compared using a standard continuous metric, such as

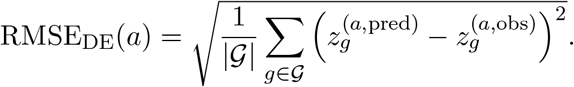

This metric evaluates agreement in both the direction and strength of gene-wise DE evidence without first converting genes into discrete DEG sets. However, it depends on the scale and calibration of the chosen DE score and can be sensitive to extreme significance values unless these are clipped or otherwise transformed.

### D.7 Weighting Schemes from Differential-Expression Evidence

This section describes constructions of feature weights from perturbation-specific differential-expression (DE) evidence. Such weights can be used in feature-wise or centroid-based metrics to give greater influence to genes with stronger evidence of a perturbation response. Most constructions derive the weights from observed data alone, such that the model prediction does not affect the weighting scheme under which it is evaluated. Some benchmark scores instead define weights using both observed and predicted responses, as discussed below.

#### General construction

For perturbation *a*, let 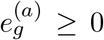 denote the DE evidence for gene *g*. Examples include an absolute effect size 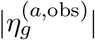, the magnitude of a test statistic 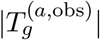, or a significance score 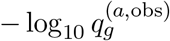.Unnormalized weights can be defined as

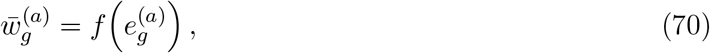

where *f* is a non-negative monotone transformation. To prevent genes with extremely large effect sizes or test statistics from dominating the resulting metric, the transformed weights can optionally be clipped:

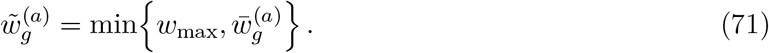

If clipping is not applied, we set 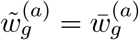. The weights can then be normalized as

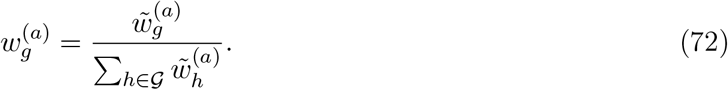

The choice of 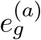 determines which evidence is emphasized, whereas the transformation *f* determines how strongly the weights are concentrated on genes with large values.

#### Example 1: Control-referenced effect-size weights

A particularly simple construction uses the magnitude of the observed perturbation-induced change, 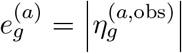, applies the identity transformation *f* (*e*) = *e*, and uses no clipping. The resulting values are then normalized. This assigns greater weight to genes with larger observed perturbation effects, irrespective of whether they are up- or down-regulated. Analogous constructions can use test-statistic magnitudes or significance scores derived from adjusted *p*-values.

#### Example 2: Perturbation-versus-rest test-statistic weights

Mejia et al. [45] derive perturbation-specific gene weights from test statistics comparing perturbation *a* with the remaining perturbations in the training set. Let

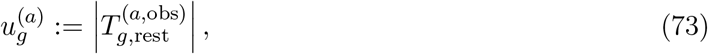

where 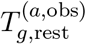 is the DE test statistic obtained by comparing perturbation *a* with the remaining perturbations.

The test-statistic magnitudes are first min–max normalized:

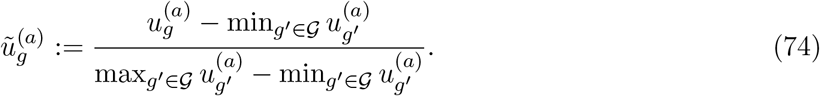

The normalized values are then squared and normalized to sum to one:

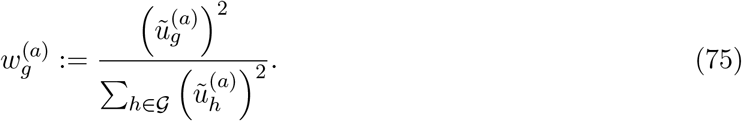

Min–max normalization places the test-statistic magnitudes on a common scale, while squaring increases the concentration of weight on genes with the strongest evidence. Because the underlying contrast compares perturbation *a* with the remaining perturbations, the resulting weights emphasize genes that distinguish that perturbation from other perturbations in the training set.

#### Example 3: Prediction-dependent smooth-gate weights

The weighted cosine similarity used in the Myllia competition is computed jointly across all perturbations and genes [48]. For each perturbation, the evaluation represents the observed response as a pseudobulk log-fold-change vector relative to the corresponding control reference, and participants submit a predicted vector in the same format. The observed perturbation-specific vectors are then concatenated into one long vector **a**, and the corresponding predicted vectors are concatenated in the same order into one long vector Thus, each coordinate *i* corresponds to one perturbation–gene pair, and *a*_*i*_ and *b*_*i*_ denote the observed and predicted log fold changes for that pair. For each coordinate, define

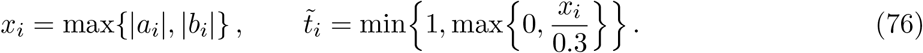

The coordinate-specific weight is then obtained using a smooth gating function,

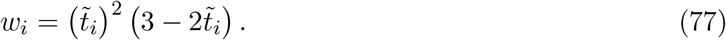

These weights enter the weighted cosine similarity as

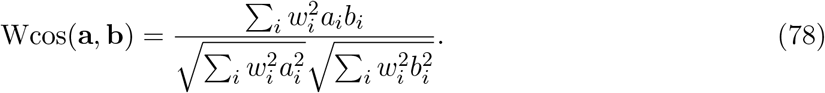

The weighting suppresses coordinates for which both the observed and predicted effects are small. Because the weight depends on the larger of the observed and predicted absolute effects, a large predicted change receives substantial weight even when the corresponding observed change is small. This makes strong false-positive effects consequential for the cosine similarity. However, because the weights depend on the prediction, predictions from different models are evaluated under different weighting schemes.

### D.8 Weighted Centroid-Based Metrics

Centroid metrics are a primary setting in which feature weighting can be introduced directly. This section defines generic weighted variants of several centroid-based metrics. Let

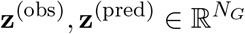

denote a target vector and a predicted vector, and let *w*_1_, …, *w*_*NG*_ ≥ 0 be feature weights with

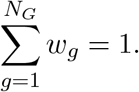

Depending on the application, **z** may represent either an absolute pseudobulk profile **y** or a reference-centered delta vector Δ.

#### Weighted mean absolute error

The weighted mean absolute error (WMAE) is defined as

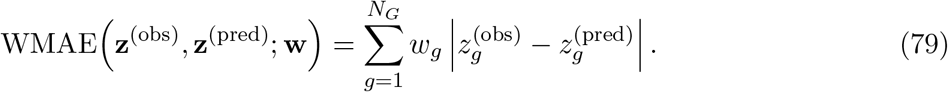

This reduces to the usual MAE when all weights are equal. A weighted MAE of this kind is used in hybrid form in [48].

#### Weighted mean squared error

The weighted mean squared error (WMSE) is defined as

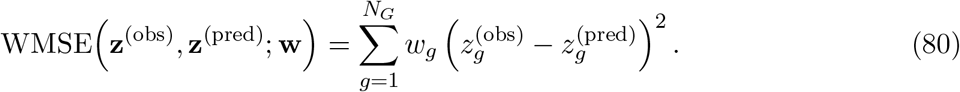

This reduces to the usual MSE when all weights are equal. DEG-based weighted variants of this form are used in [45] and [46].

#### Weighted Pearson correlation

Define the weighted means of the target and predicted vectors by

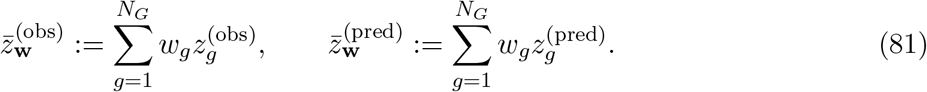

The weighted Pearson correlation is then defined as

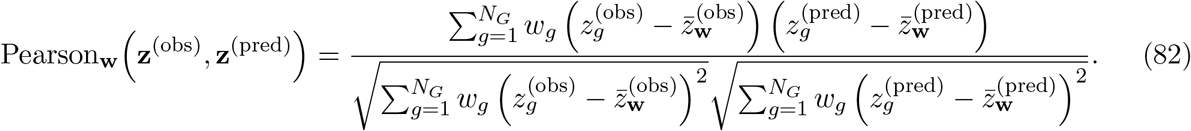

This reduces to the usual Pearson correlation when all weights are equal. The weighted Pearson correlation is undefined if either weighted variance term in the denominator is zero. In the SBB framework, a DEG-weighted Pearson-Δ_Pert_ metric is used by applying weighted Pearson correlation to perturbation-effect vectors, with weights chosen to emphasize genes carrying perturbation-specific signal [69].

#### Weighted coefficient of determination

Define the weighted mean of the target vector by

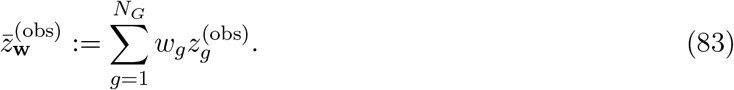

The weighted coefficient of determination is then defined as

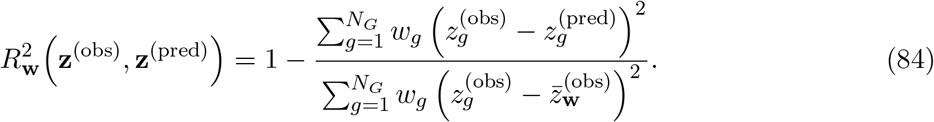

This reduces to the usual *R*^2^ when all weights are equal. The weighted *R*^2^ is undefined if the weighted variance of the target vector is zero. The delta-based weighted *R*^2^ used in [46] is obtained by substituting the perturbation delta vectors for **z**^(obs)^ and **z**^(pred)^ in the same formula.

### D.9 Classifier-Based Distances

Classifier-based distances quantify the discrepancy between two populations by how well a supervised classifier can separate their cells. Given labeled samples from *v*^(obs)^ and *v*^(pred)^, one trains a classifier and converts its held-out performance into a score, for example classification accuracy, AUROC, or cross-entropy. If the two populations are indistinguishable, performance should remain near chance; stronger-than-chance discrimination indicates a discrepancy between the underlying distributions. This makes classifier-based scores closely related to classifier two-sample tests and, in principle, to joint distribution metrics. However, unlike closed-form distances, the reported value depends not only on the distributions themselves but also on the chosen hypothesis class, optimization procedure, regularization, train-test split, and sample size [7, 42].

Classifier-based separability has been used in settings such as PhenoDissim [77]. In perturbation-response model evaluation, however, direct classifier-based distances remain relatively uncommon compared to other metrics, which is why they are not treated as a main category in the body text.

#### D.9.1 Classifier Control Probability

Ji et al. [31] describe a classifier-based score that compares a perturbation population *X*^*(a)*^ *against* a control population *X* ^*(ctrl)*^. *A lo*gistic-regression classifier is trained to distinguish perturbation from control cells, with 20% of the perturbation cells held out for evaluation. Writing 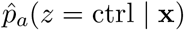 for the fitted posterior probability of the control label, the control-probability score described in the supplement is

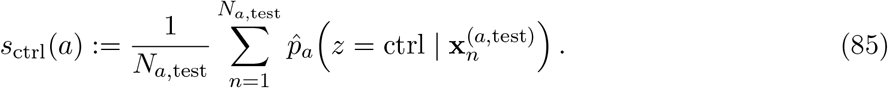

Low values of *s*_ctrl_(*a*) indicate that held-out perturbation cells are confidently distinguished from controls. The corresponding distance-like quantity is

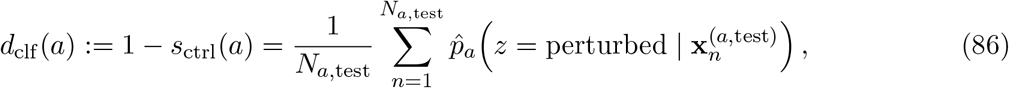

where the equality follows because the classifier is binary. The released PertPy implementation directly returns the perturbation-class probability in the final expression, and therefore implements *d*_clf_ (*a*) rather than *s*_ctrl_(*a*) itself. Accordingly, larger PertPy values indicate greater classifier separability.

#### D.9.2 Classifier class projection

A related variant described in the supplementary material of [31] and in [28] is the classifier class projection score. For perturbation *a*, a logistic-regression classifier is trained on all cells not belonging to perturbation *a* together with the control cells, and the average predicted probability of the matched control label is then evaluated on cells from perturbation *a*. In notation parallel to the above,

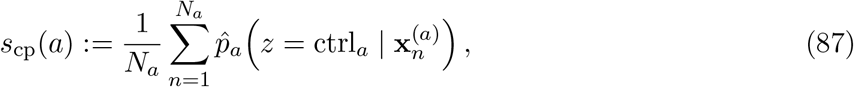

where ctrl_*a*_ denotes the relevant control label for perturbation *a*. Again, Ji et al. [31] summarize the corresponding distance-like version as 1−*s*_cp_(*a*). They also report that this variant performed substantially worse than the other benchmarked metrics and excluded it from their higher-level performance grouping.

### D.10 Retrieval-Based Evaluation Metrics

Retrieval-based metrics evaluate whether predicted perturbation responses preserve the relative ordering of perturbations under a chosen representation and comparison score. Depending on the benchmark, retrieval may be defined at the level of matching observed and predicted response profiles, recovering phenocopy relationships, or retrieving perturbations with downstream phenotype labels. In contrast to one-to-one distance metrics, retrieval-based metrics are inherently relative: a prediction is judged not only by its distance to the matching observed response, but also by its distances or similarities to other candidate perturbations.

#### D.10.1 Retrieval Rank

For perturbation *a*, the retrieval-rank metric of [73] is defined as

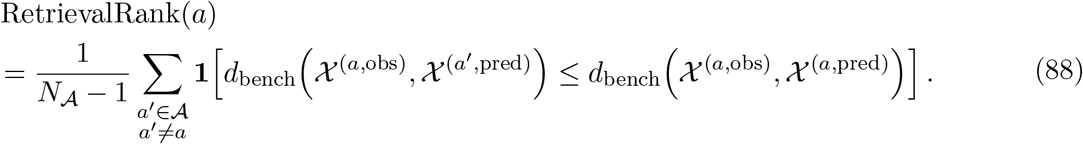

Here, *N*_*A*_ denotes the total number of considered perturbations, and *d*_bench_ is the shorthand for a benchmark-specific metric characterized by the representation map *ϕ* and metric *d*. This score fixes the observed target for perturbation *a* and asks how many non-matching predicted perturbation responses are at least as close to that target as the matching prediction is. Thus, lower values indicate better retrieval, with 0 corresponding to the case where no non-matching prediction is closer than the matching one.

PerturBench randomly permutes candidates before sorting, producing random tie-breaking. The formula above gives the expected normalized rank under uniform random ordering within each tie group. Exact code reproduction instead requires fixing and reporting the random seed.

#### D.10.2 Transposed Retrieval Rank

The transposed retrieval-rank metric is defined as

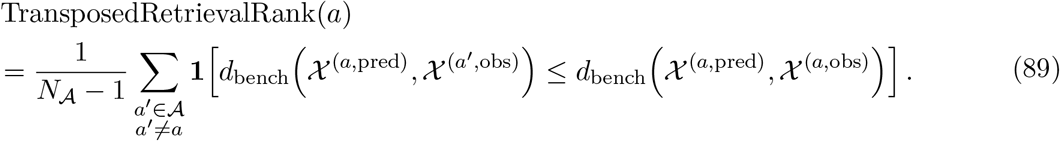

The two metrics differ only in which object is held fixed during retrieval: retrieval rank fixes the observed target and ranks predictions against it, whereas transposed retrieval rank fixes the prediction and ranks observed perturbation states against it. The transposed variant therefore asks whether the prediction for perturbation *a* is closer to its matching observed perturbation state than to observed states from other perturbations.

Several related metrics can be viewed as variants of this same transposed retrieval principle. For example, centroid accuracy evaluates whether the predicted centroid for perturbation *a* is closest to the matching observed centroid among a set of candidate observed perturbation centroids. In this case, *d*_bench_ is instantiated as a centroid-level discrepancy, and the desired event is

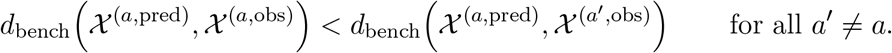

Similarly, PDS-style perturbation discrimination scores use the same ordering logic: a predicted response is considered correctly discriminated if it is closer to the matched observed perturbation response than to mismatched observed perturbation responses. These variants differ mainly in the representation *ϕ*, the base metric *d*, the candidate set of perturbations, and whether the result is reported as a rank, a top-1 accuracy, or a permutation-calibrated discrimination statistic.

#### D.10.3 Phenocopy Retrieval

A related retrieval formulation is used by Littman et al. [40] to evaluate whether predicted perturbation responses recover phenocopy relationships between perturbations. Let **z**^(*a*,obs)^ and **z**^(*a*,pred)^ denote benchmark-specific perturbation representations derived from observed and predicted responses, respectively. In [40], these representations are obtained from pseudobulk log-fold-change profiles after PCA, sphering, and cosine-similarity-based comparison.

For an anchor perturbation *a*, define the observed and predicted similarities to a candidate perturbation *b* as

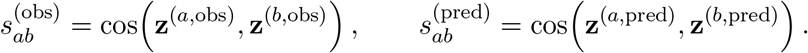

For a fixed retrieval budget *K*, the ground-truth and predicted phenocopy sets are

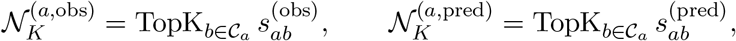

where *C*_*a*_ is the candidate set of perturbations. The fixed-budget phenocopy retrieval score is then

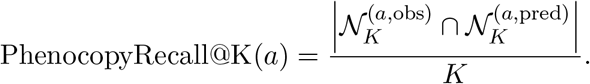

Littman et al. [40] use *K* = 10 and primarily treat significant-effect training perturbations as anchors, with test perturbations as candidates. They also report an AUROC variant in which ground-truth phenocopies are defined by a similarity threshold based on the median plus four median absolute deviations of the anchor’s observed similarity distribution, rather than by a fixed top-*K* cutoff.

#### D.10.4 Hit-Based Retrieval

Retrieval can also be defined with respect to downstream phenotype or hit labels rather than response-profile matching. Let A denote a candidate set of perturbations. For each perturbation *a* ∈ *A, let h*_*a*_ ∈ {0, 1} denote whether *a* is a ground-truth hit for a specified phenotype, and let *s*_*a*_ ∈ ℝ denote a predicted score, with larger values indicating a higher predicted probability of being a hit.

In Littman et al. [40], this construction is applied to gene-set phenotypes. For a gene set *m*, let 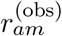 and 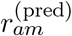 denote the observed and predicted gene-set scores for perturbation *a*, respectively.With median_*m*_ and MAD_*m*_ denoting the median and median absolute deviation of the observed scores 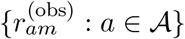, the up-regulation and down-regulation hit labels can be written as

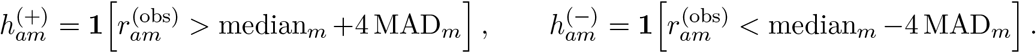

Predicted gene-set scores are then used to rank perturbations. The up-regulation score is 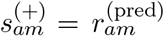, whereas the down-regulation score is sign-flipped, 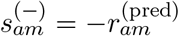. Performance is summarized by AUROC for the binary labels 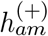 and 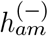 under these corresponding scores.

Bereket and Leskovec [6] use the same broad outcome-retrieval structure for hit discovery. Given binary hit labels *h*_*a*_ and predicted hit scores *s*_*a*_, let

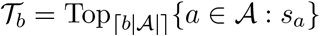

denote the top-ranked perturbations selected under experiment budget *b* ∈ [0, 1]. The prioritization score is recall at that budget,

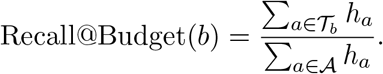

For simulation, let *T*_*k*_ denote the top *k* ranked perturbations and define the empirical false discovery rate at cutoff *k* as

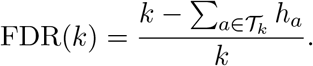

For a target FDR *q*, the reported score is the maximum recall over all cutoffs satisfying the FDR constraint,

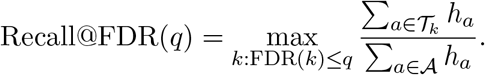

Thus, both formulations evaluate whether predicted rankings retrieve perturbations with a desired downstream phenotype, rather than whether each prediction retrieves its matching observed response profile.

### D.11 Bound Discrimination Score

Introduced in [69], the bound discrimination score (BDS) tests whether an evaluation protocol can distinguish a technical duplicate from an uninformative control reference. Let *P be the set of evaluated* perturbations and let

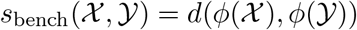

denote the benchmark-specific score induced by the representation map *ϕ* and metric *d*. We assume throughout this subsection that smaller values of *s*_bench_ indicate better agreement; similarities for which larger values are better should first be converted to smaller-is-better scores.

For each perturbation *a* ∈ *P, its observed cells ar*e randomly divided into two disjoint half-bags, 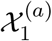 and 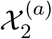. In the formal BDS definition, 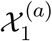 serves as the ground truth and 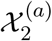 as the positive reference. The negative reference is a bag X^(ctrl)^ of observed control cells. The corresponding positive-reference score is

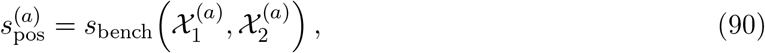

and the negative-reference score is

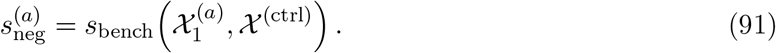

The protocol gives the desired ordering for perturbation *a* when 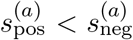. The point-estimate BDS is therefore

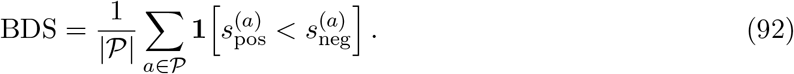

It reports the fraction of perturbations for which the protocol correctly orders the positive and negative references, but it does not by itself quantify how surprising the observed separation is under sampling variation.

#### Permutation test

Vollenweider and Bühlmann [69] supplement this point estimate with a separate permutation test for each perturbation. In the released implementation, the test statistic treats the two perturbation halves symmetrically by averaging their respective scores against the same control bag,

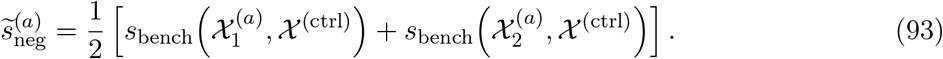

The observed separation is then

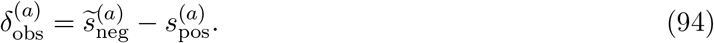

Because smaller scores indicate better agreement, a positive value indicates that the two technical-duplicate halves are closer to one another than either half is, on average, to the controls.

For perturbation *a*, let *n*_*a*_ and *n*_ctrl_ be the numbers of perturbation and control cells used in the comparison. The *n*_*a*_ + *n*_ctrl_ cells are pooled, and their perturbation-versus-control labels are randomly reassigned while preserving these group sizes. The first *n*_*a*_ permuted cells form a pseudo-perturbation group and the remaining *n*_ctrl_ cells form a pseudo-control group. The pseudo-perturbation group is then divided into two half-bags with the same sizes as the observed split. Thus, unlike the PDS permutation test, the BDS test does not keep preformed bags fixed or permute perturbation identities across bags; it permutes individual-cell labels between one perturbation and the controls.

For permutation *m*, the same symmetric calculation gives

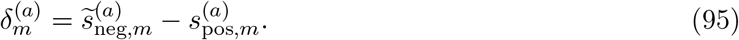

Repeating this procedure *M* times yields a null distribution under the hypothesis that cells from perturbation *a* and the controls are exchangeable. Because larger positive values provide stronger evidence that technical duplicates score better than controls, SBB uses the upper-tail empirical p-value

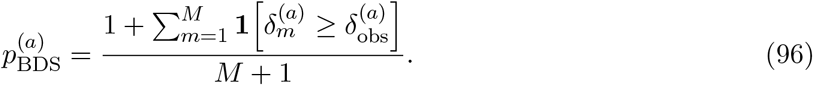

The added one in the numerator and denominator prevents a zero p-value and gives the usual finite-permutation correction. SBB use *M* = 999 permutations in their reported supplementary comparison. A small p-value indicates that the protocol separates the perturbation from the control background more strongly than expected under random reassignment.

For weighted protocols, such as DEG-weighted MSE or Pearson correlation, the perturbation-specific gene weights are held fixed at their observed values throughout the permutations; they are not re-estimated after each label reassignment. Any reference vectors used by delta-based metrics are likewise held fixed. The resulting test therefore asks whether the observed separation exceeds chance separation under the specified, fixed protocol. Because the weights may themselves have been estimated using the evaluated data, this does not remove all data dependence associated with selecting the weighting scheme.

Finally, the perturbation-wise p-values are adjusted across *P using the Benjamini*–Hochberg procedure. SBB summarize the result for a protocol by reporting the fraction of perturbations whose adjusted p-values remain below the chosen significance threshold. This significance fraction complements the BDS point estimate: BDS records whether the ordering is correct, whereas the permutation analysis asks whether the separation is sufficiently large to be distinguished from sampling variation.

### D.12 Dynamic Range Fraction

Introduced in [46], the dynamic range fraction (DRF) is a meta-metric for assessing the signal recovery of a benchmark metric. Let

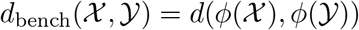

denote the benchmark-specific metric induced by the representation map *ϕ* and metric *d*. For perturbation *a*, let *X*^(*a*,obs)^ denote the ground-truth observed population, let *X*^(*a*,pos)^ denote a positive reference population, such as a technical duplicate, and let *X*^(*a*,neg)^ denote an uninformative negative reference population. The corresponding benchmark scores are

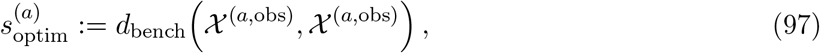

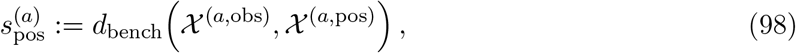

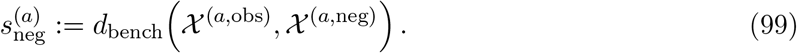

Ideally, these scores satisfy

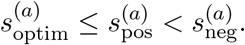

The bound discrimination score only checks the weaker condition 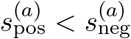.

To quantify not only whether the positive reference outperforms the negative reference, but also how much of the available dynamic range is recovered, define

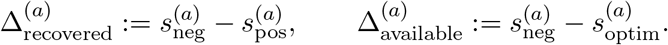

The dynamic-range fraction is then

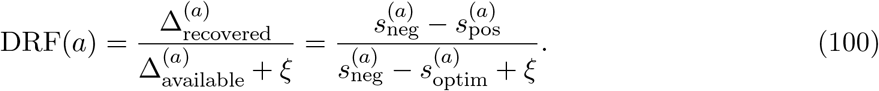

Here, *ξ >* 0 is a small numerical constant that stabilizes cases where little dynamic range is available.

The interpretation is direct. If DRF(*a*) = 1, the positive reference achieves the optimal score. If DRF(*a*) = 0, the positive and negative references perform equally. If DRF(*a*) *<* 0, the positive reference performs worse than the negative reference. Values outside [0, 1] can occur because of sampling variation, for example if the positive reference scores better than the self-comparison or worse than the negative reference.

In the original DRF formulation of [46], both references are defined after restricting to a pseudobulk, or centroid-based, representation. Concretely, each population *X*^*(a)*^ *is mappe*d to its mean expression profile. As a negative reference, the training-perturbation mean is used: a global mean baseline that assigns the same average response to every perturbation *a* and therefore ignores perturbation-specific effects.

For the positive reference, two constructions are considered. The first is the *technical duplicate* (Section 5.1), obtained by randomly splitting the observed cells for perturbation *a* into two disjoint subsets. One split is used as the observed response, while the mean profile of the other, **y**^(*a*,td)^ = *ϕ* (*X* (^(*a*,td)^), serves as a surrogate prediction. This provides an estimate of the performance achievable under an ideal repeat of the same experiment, where discrepancies arise only from intrinsic variability and sampling noise.

Because this estimator can be unstable when the number of cells per perturbation is small, a second construction, the *interpolated technical duplicate*, is introduced. It is defined as a gene-wise interpolation between the technical duplicate and the training-perturbation mean,

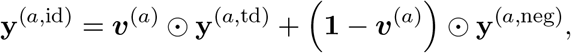

where the weights are defined as 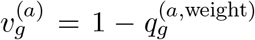, and 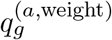 corresponds to adjusted *p*-values derived from differential expression for gene *g* under perturbation *a* as compared to all other perturbed cells. This yields a more stable positive reference that preserves perturbation-specific signal for strongly affected genes while reverting to the training-perturbation mean for unaffected genes.

## Footnotes

1 Throughout this work, we use the term *metric* in a broad sense, without requiring the mathematical properties of a metric.

## References

[1] Britt Adamson et al. “A Multiplexed Single-Cell CRISPR Screening Platform Enables Systematic Dissection of the Unfolded Protein Response”. In: Cell 167.7 (Dec. 2016), 1867–1882.e21. ISSN: 0092-8674. DOI: 10.1016/j.cell.2016.11.048.

[2] Abhinav K. Adduri et al. “Predicting Cellular Responses to Perturbation across Diverse Contexts with State”. In: bioRxiv (July. 2025), p. 2025.06.26.661135. ISSN: 2692-8205. DOI: 10.1101/2025.06.26.661135.

[3] Constantin Ahlmann-Eltze and Wolfgang Huber. “Comparison of Transformations for Single-Cell RNA-Seq Data”. In: Nature Methods 20.5 (May. 2023), pp. 665–672. ISSN: 1548-7105. DOI: 10.1038/s41592-023-01814-1. (Visited on 01/12/2025).

[4] Constantin Ahlmann-Eltze, Wolfgang Huber, and Simon Anders. “Deep-Learning-Based Gene Perturbation Effect Prediction Does Not yet Outperform Simple Linear Baselines”. In: Nature Methods 22.8 (Aug. 2025), pp. 1657–1661. ISSN: 1548-7105. DOI:10.1038/s41592-025-02772-6.

[5] Arc Institute. Virtual Cell Challenge 2025 Wrap-Up: Winners and Reflections. News article. Dec. 2025. URL: https://arcinstitute.org/news/virtual-cell-challenge-2025-wrap-up.

[6] Ding Bai et al. “AttentionPert: Accurately Modeling Multiplexed Genetic Perturbations with Multi-Scale Effects”. In: Bioinformatics 40.Supplement_1 (June. 2024), pp. i453–i461. ISSN: 1367-4803, 1367-4811. DOI: 10.1093/bioinformatics/btae244. (Visited on 12/07/2024).

[7] Ihab Bendidi et al. “Benchmarking Transcriptomics Foundation Models for Perturbation Analysis : One PCA Still Rules Them All”. In: arXiv arXiv: 2410.13956 (Nov. 2024). DOI: 10.48550/arXiv.2410.13956. arXiv: 2410.13956 [cs].

[8] Michael D. Bereket and Jure Leskovec. Are Current AI Virtual Cell Models Useful for Scientific Discovery? Apr. 2026. DOI: 10.64898/2026.04.23.719015. (Visited on 04/27/2026).

[9] Judith Bernett et al. From Hype to Health Check: Critical Evaluation of Drug Response Prediction Models with DrEval. May 2025. DOI: 10.1101/2025.05.26.655288. (Visited on 01/29/2026).

[10] Mikołaj Bińkowski et al. “Demystifying MMD GANs”. In: International Conference on Learning Representations. Feb. 2018. (Visited on 04/13/2026).

[11] Sebastian Bischoff et al. “A Practical Guide to Sample-Based Statistical Distances for Evaluating Generative Models in Science”. In: arXiv arXiv: 2403.12636 (Oct. 2024). DOI: 10.48550/arXiv.2403.12636. arXiv: 2403.12636.

[12] Niek Brouwer et al. Drug Response Prediction Provides a Biologically Relevant Benchmark for Perturbation Response Models. Dec. 2025. DOI: 10.64898/2025.12.09.693213. (Visited on 04/14/2026).

[13] Charlotte Bunne et al. “How to Build the Virtual Cell with Artificial Intelligence: Priorities and Opportunities”. In: Cell 187.25 (Dec. 2024), pp. 7045–7063. ISSN: 0092-8674, 1097-4172. DOI: 10.1016/j.cell.2024.11.015.

[14] Charlotte Bunne et al. “Learning Single-Cell Perturbation Responses Using Neural Optimal Transport”. In: Nature Methods 20.11 (Nov. 2023), pp. 1759–1768. ISSN: 1548-7091, 1548-7105. DOI: 10.1038/s41592-023-01969-x.

[15] Tiffany Callahan et al. “Virtual Cells as Causal World Models: A Perspective on Evaluation”. In: Proceedings of the Workshop on AI Virtual Cells and Instruments: A New Era in Drug Discovery and Development at NeurIPS 2025. San Diego, CA, USA, 2025. URL: https://neurips.cc/virtual/2025/loc/san-diego/129556.

[16] Safiye Celik et al. “Building, Benchmarking, and Exploring Perturbative Maps of Transcriptional and Morphological Data”. In: PLOS Computational Biology 20.10 (Oct. 2024). Ed. by Thouis Ray Jones, e1012463. ISSN: 1553-7358. DOI: 10.1371/journal.pcbi.1012463. (Visited on 12/07/2024).

[17] Srinivas Niranj Chandrasekaran et al. JUMP Cell Painting Dataset: Morphological Impact of 136,000 Chemical and Genetic Perturbations. Mar. 2023. DOI: 10.1101/2023.03.23.534023. (Visited on 12/07/2024).

[18] Yubao Cheng et al. “Sequencing-Free Whole-Genome Spatial Transcriptomics at Single-Molecule Resolution”. In: Cell 188.24 (Nov. 2025), 6953–6970.e12. ISSN: 0092-8674, 1097-4172. DOI: 10.1016/j.cell.2025.09.006. (Visited on 06/21/2026).

[19] Mathieu Chevalley et al. “A Large-Scale Benchmark for Network Inference from Single-Cell Perturbation Data”. In: Communications Biology 8.1 (Mar. 2025), pp. 1–18. ISSN: 2399-3642. DOI: 10.1038/s42003-025-07764-y.

[20] Helena L. Crowell et al. “Muscat Detects Subpopulation-Specific State Transitions from Multi-Sample Multi-Condition Single-Cell Transcriptomics Data”. In: Nature Communications 11.1 (Nov. 2020), p. 6077. ISSN: 2041-1723. DOI: 10.1038/s41467-020-19894-4. (Visited on 12/07/2024).

[21] Gerold Csendes. Assessing a Virtual Cell’s Utility. https://blog.turbine.ai/p/assessing-a-virtual-cells-utility. mJune 2025. (Visited on 05/29/2026).

[22] Gerold Csendes et al. “Benchmarking Foundation Cell Models for Post-Perturbation RNA-Seq Prediction”. In: BMC Genomics 26.1 (Apr. 2025), p. 393. ISSN: 1471-2164. DOI: 10.1186/s12864-025-11600-2.

[23] Haotian Cui et al. “scGPT: Toward Building a Foundation Model for Single-Cell Multi-Omics Using Generative AI”. In: Nature Methods 21.8 (Aug. 2024), pp. 1470–1480. ISSN: 1548-7105. DOI: 10.1038/s41592-024-02201-0.

[24] Payam Dibaeinia et al. Virtual Cells Need Context, Not Just Scale. Feb. 2026. DOI: 10.64898/2026.02.04.703804. (Visited on 05/22/2026).

[25] Daniel Dimitrov et al. “Interpretation, Extrapolation, and Perturbation of Single Cells”. In: Nature Reviews Genetics (Jan. 2026), pp. 1–22. ISSN: 1471-0064. DOI: 10.1038/s41576-025-00920-4. (Visited on 01/07/2026).

[26] Atray Dixit et al. “Perturb-Seq: Dissecting Molecular Circuits with Scalable Single-Cell RNA Profiling of Pooled Genetic Screens”. In: Cell 167.7 (Dec. 2016), 1853–1866.e17. ISSN: 00928674. DOI: 10.1016/j.cell.2016.11.038.

[27] Mingze Dong et al. “Stack: In-Context Learning of Single-Cell Biology”. In: bioRxiv (2026). DOI: 10.64898/2026.01.09.698608. URL: https://connect.biorxiv.org/relate/content/224/channel.

[28] Laura M. Drepanos et al. Direct Comparison of CRISPR Knockout and Interference with Perturb-Seq. July 2026. DOI: 10.64898/2026.07.04.736492. (Visited on 07/07/2026).

[29] Caleb N. Ellington et al. Cell-Level Virtual Screening. May 2026. DOI: 10.64898/2026.05.11.724149. (Visited on 05/14/2026).

[30] Eric and Wendy Schmidt Center at the Broad Institute and Diabetes Initiative at the Broad Institute of MIT and Harvard. Obesity Machine Learning Competition: Tackling Metabolic Diseases. Cell Perturbation Prediction Challenge. 2025. URL: https://www.broadinstitute.org/videos/obesity-machine-learning-competition-tackling-metabolic-diseases-overview.

[31] Federico Borra. “In Silico Perturbation of Single Cells-A Spiking Protocol for Metric Evaluation”. MA thesis. Politecnico di Torino, Oct. 2024. URL: https://webthesis.biblio.polito.it/view/year/2024/corso %3D2Bdi% 3D2Blaurea%3D2Bmagistrale%3D2Bin%3D2Bdata%3D2Bscience%3D2Band%3D2Bengineering.title.html.

[32] Nicolas Fournier and Arnaud Guillin. “On the Rate of Convergence in Wasserstein Distance of the Empirical Measure”. In: Probability Theory and Related Fields 162.3 (Aug. 2015), pp. 707–738. ISSN: 1432-2064. DOI: 10.1007/s00440-014-0583-7. (Visited on 04/18/2026).

[33] Chris J. Frangieh et al. “Multimodal Pooled Perturb-CITE-Seq Screens in Patient Models Define Mechanisms of Cancer Immune Evasion”. In: Nature Genetics 53.3 (Mar. 2021), pp. 332–341. ISSN: 1546-1718. DOI: 10.1038/s41588-021-00779-1. (Visited on 02/17/2025).

[34] Chris J. Frangieh et al. “Multimodal Pooled Perturb-CITE-Seq Screens in Patient Models Define Mechanisms of Cancer Immune Evasion”. In: Nature Genetics 53.3 (Mar. 2021), pp. 332–341. ISSN: 1546-1718. DOI: 10.1038/s41588-021-00779-1.

[35] Siyu He et al. “Squidiff: Predicting Cellular Development and Responses to Perturbations Using a Diffusion Model”. In: Nature Methods (Nov. 2025), pp. 1–13. ISSN: 1548-7105. DOI: 10.1038/s41592-025-02877-y.

[36] Mahshid Heidari et al. “Evaluating Single-Cell Perturbation Response Models Is Far from Straightforward”. In: bioRxiv (Feb. 2026), p. 2026.02.14.705879. ISSN: 2692-8205. DOI: 10.64898/2026.02.14.705879.

[37] Lukas Heumos et al. “Pertpy: An End-to-End Framework for Perturbation Analysis”. In: Nature Methods 23.2 (Feb. 2026), pp. 350–359. ISSN: 1548-7105. DOI: 10.1038/s41592-025-02909-7.

[38] Martin Heusel et al. “GANs Trained by a Two Time-Scale Update Rule Converge to a Local Nash Equilibrium”. In: Advances in Neural Information Processing Systems (NeurIPS). arXiv:1706.08500. Jan. 2018. DOI: 10.48550/arXiv.1706.08500. eprint: 1706.08500.

[39] Kexin Huang et al. Sequential Optimal Experimental Design of Perturbation Screens Guided by Multi-Modal Priors. Dec. 2023. DOI: 10.1101/2023.12.12.571389. (Visited on 02/17/2025).

[40] Xingfan Huang et al. “Single-Cell, Whole-Embryo Phenotyping of Mammalian Developmental Disorders”. In: Nature 623.7988 (Nov. 2023), pp. 772–781. ISSN: 1476-4687. DOI: 10.1038/s41586-023-06548-w. (Visited on 05/22/2026).

[41] Trey Ideker, Janusz Dutkowski, and Leroy Hood. “Boosting Signal-to-Noise in Complex Biology: Prior Knowledge Is Power”. In: Cell 144.6 (Mar. 2011), pp. 860–863. ISSN: 0092-8674. DOI: 10.1016/j.cell.2011.03.007. (Visited on 05/04/2026).

[42] Diego Adhemar Jaitin et al. “Dissecting Immune Circuits by Linking CRISPR-Pooled Screens with Single-Cell RNA-Seq”. In: Cell 167.7 (Dec. 2016), 1883–1896.e15. ISSN: 0092-8674, 1097-4172. DOI: 10.1016/j.cell.2016.11.039. (Visited on 04/19/2026).

[43] Yunhui Jang et al. Towards Autonomous Mechanistic Reasoning in Virtual Cells. May 2026. DOI: 10.48550/arXiv.2604.11661. arXiv: 2604.11661 [cs.LG]. (Visited on 05/29/2026).

[44] Sadeep Jayasumana et al. “Rethinking FID: Towards a Better Evaluation Metric for Image Generation”. In: 2024 IEEE/CVF Conference on Computer Vision and Pattern Recognition (CVPR). Seattle, WA, USA: IEEE, June 2024, pp. 9307–9315. ISBN: 979-8-3503-5300-6. DOI: 10.1109/CVPR52733.2024.00889. (Visited on 03/31/2026).

[45] Yuge Ji et al. “Optimal Distance Metrics for Single-Cell RNA-Seq Populations”. In: bioRxiv (Dec. 2023), p. 2023.12.26.572833. DOI: 10.1101/2023.12.26.572833.

[46] Qun Jiang et al. “scPRAM Accurately Predicts Single-Cell Gene Expression Perturbation Response Based on Attention Mechanism”. In: Bioinformatics 40.5 (May. 2024), btae265. ISSN: 1367-4811. DOI: 10.1093/bioinformatics/btae265.

[47] Alexandr A. Kalinin et al. “A Versatile Information Retrieval Framework for Evaluating Profile Strength and Similarity”. In: Nature Communications 16.1 (June. 2025), p. 5181. ISSN: 2041-1723. DOI: 10.1038/s41467-025-60306-2.

[48] Sayash Kapoor and Arvind Narayanan. “Leakage and the Reproducibility Crisis in Machine-Learning-Based Science”. In: Patterns 4.9 (Sept. 2023), p. 100804. ISSN: 2666-3899. DOI: 10.1016/j.patter.2023.100804. (Visited on 11/05/2025).

[49] Eric Kernfeld. We predict that the transcript abundance from the gene we are knocking down will (checks notes) decrease. Blog post. Dec. 2025. URL: https://ekernf01.github.io/target_gene_shenanigans (visited on 07/11/2026).

[50] Eric Kernfeld et al. “A Comparison of Computational Methods for Expression Forecasting”. In: Genome Biology 26.1 (Nov. 2025), p. 388. ISSN: 1474-760X. DOI: 10.1186/s13059-025-03840-y.

[51] Dominik Klein et al. “CellFlow Enables Generative Single-Cell Phenotype Modeling with Flow Matching”. In: bioRxiv (Apr. 2025), p. 2025.04.11.648220. DOI: 10.1101/2025.04.11.648220.

[52] Soheil Kolouri et al. “Generalized Sliced Wasserstein Distances”. In: Advances in Neural Information Processing Systems. Vol. 32. Curran Associates, Inc., 2019. URL:https://proceedings.neurips.cc/paper/2019/hash/f0935e4cd5920aa6c7c996a5ee53a70f-Abstract.html.

[53] Takamasa Kudo et al. “Multiplexed, Image-Based Pooled Screens in Primary Cells and Tissues with PerturbView”. In: Nature Biotechnology 43.7 (July. 2025), pp. 1091–1100. ISSN: 1546-1696. DOI: 10.1038/s41587-024-02391-0. (Visited on 02/20/2026).

[54] Chen Li, Lei Wei, and Xuegong Zhang. Unified Multimodal Learning Enables Generalized Cellular Response Prediction to Diverse Perturbations. Jan. 2026. DOI: 10.1101/2025.11.13.688367. (Visited on 04/28/2026).

[55] Chen Li et al. “Benchmarking AI Models for In Silico Gene Perturbation of Cells”. In: bioRxiv (Jan. 2025), p. 2024.12.20.629581. DOI: 10.1101/2024.12.20.629581.

[56] Lanxiang Li et al. “A Systematic Comparison of Single-Cell Perturbation Response Prediction Models”. In: bioRxiv (Sept. 2025), p. 2024.12.23.630036. ISSN: 2692-8205. DOI: 10.1101/2024.12.23.630036.

[57] Arthur Liberzon et al. “The Molecular Signatures Database Hallmark Gene Set Collection”. In: Cell Systems 1.6 (Dec. 2015), pp. 417–425. ISSN: 24054712. DOI: 10.1016/j.cels.2015.12.004. (Visited on 12/07/2024).

[58] Zaikang Lin et al. Interpretable Neural ODEs for Gene Regulatory Network Discovery under Perturbations. Jan. 2025. DOI: 10.48550/arXiv.2501.02409. arXiv: 2501.02409 [cs]. (Visited on 01/11/2025).

[59] Russell Littman et al. “Gene-Embedding-Based Prediction and Functional Evaluation of Perturbation Expression Responses with PRESAGE”. In: bioRxiv (June. 2025), p. 2025.06.03.657653. DOI: 10.1101/2025.06.03.657653.

[60] Qiyuan Liu et al. Effects of Distance Metrics and Scaling on the Perturbation Discrimination Score. Nov. 2025. DOI: 10.48550/arXiv.2511.16954. arXiv: 2511.16954 [stat].

[61] Romain Lopez et al. “Deep Generative Modeling for Single-Cell Transcriptomics”. In: Nature Methods 15.12 (Dec. 2018), pp. 1053–1058. ISSN: 1548-7091, 1548-7105. DOI: 10.1038/s41592-018-0229-2.

[62] Mohammad Lotfollahi et al. “Predicting Cellular Responses to Complex Perturbations in High-throughput Screens”. In: Molecular Systems Biology 19.6 (June. 2023), e11517. ISSN: 1744-4292, 1744-4292. DOI: 10.15252/msb.202211517.

[63] Malte D. Luecken et al. “Benchmarking Atlas-Level Data Integration in Single-Cell Genomics”. In: Nature Methods 19.1 (Jan. 2022), pp. 41–50. ISSN: 1548-7091, 1548-7105. DOI: 10.1038/s41592-021-01336-8. (Visited on 12/07/2024).

[64] Lena Maier-Hein et al. “Metrics Reloaded: Recommendations for Image Analysis Validation”. In: Nature Methods 21.2 (Feb. 2024), pp. 195–212. ISSN: 1548-7105. DOI: 10.1038/s41592-023-02151-z. (Visited on 11/27/2025).

[65] Lena Maier-Hein et al. “Why Rankings of Biomedical Image Analysis Competitions Should Be Interpreted with Care”. In: Nature Communications 9.1 (Dec. 2018), p. 5217. ISSN: 2041-1723. DOI: 10.1038/s41467-018-07619-7. (Visited on 04/14/2026).

[66] Xinjie Mao et al. Benchmarking Virtual Cell Models for In-the-Wild Perturbation Response. Apr. 2026. DOI: 10.48550/arXiv.2604.27646. arXiv: 2604.27646 [q-bio]. (Visited on 05/01/2026).

[67] Matthew B. A. McDermott et al. “A Closer Look at AUROC and AUPRC under Class Imbalance”. In: arXiv arXiv: 2401.06091 (Jan. 2025). DOI: 10.48550/arXiv.2401.06091. arXiv: 2401.06091 [cs].

[68] Gabriel M. Mejia et al. “Diversity by Design: Addressing Mode Collapse Improves scRNA-seq Perturbation Modeling on Well-Calibrated Metrics”. In: arXiv (2025). DOI: 10.48550/arXiv.2506.22641. arXiv: 2506.22641. URL: https://arxiv.org/abs/2506.22641.

[69] Henry E. Miller et al. “Deep Learning-Based Genetic Perturbation Models Do Outperform Uninformative Baselines on Well-Calibrated Metrics”. In: bioRxiv (Oct. 2025), p. 2025.10.20.683304. ISSN: 2692-8205. DOI: 10.1101/2025.10.20.683304.

[70] Jean-Baptiste Morlot et al. TwinCell: Large Causal Cell Model for Reliable and Interpretable Therapeutic Target Prioritisation. Mar. 2026. DOI: 10.64898/2026.01.29.702072. (Visited on 03/11/2026).

[71] Myllia. Echoes of Silenced Genes: A Cell Challenge. https://www.kaggle.com/competitions/echoes-of-silenced-genes. Kaggle competition page. Accessed: 2026-04-19. 2026.

[72] Ajay Nadig et al. “Transcriptome-Wide Analysis of Differential Expression in Perturbation Atlases”. In: Nature Genetics 57.5 (May. 2025), pp. 1228–1237. ISSN: 1546-1718. DOI: 10.1038/s41588-025-02169-3.

[73] Phillip B. Nicol, Shriya Shivakumar, and Rafael A. Irizarry. Spurious Correlation Inflates Performance in Single-Cell Perturbation Prediction. May 2026. DOI: 10.64898/2026.05.07.723486. (Visited on 05/18/2026).

[74] Emmanuel Noutahi et al. Virtual Cells: Predict, Explain, Discover. June 2025. DOI: 10.48550/arXiv.2505.14613. arXiv: 2505.14613 [cs]. (Visited on 09/07/2025).

[75] Giovanni Palla. Virtual Cell Perturbation Metrics Reloaded. Substack Newsletter. Nov. 2025. (Visited on 11/27/2025).

[76] Giovanni Palla et al. “Scalable Single-Cell Gene Expression Generation with Latent Diffusion Models”. In: arXiv arXiv: 2511.02986 (Nov. 2025). DOI: 10.48550/arXiv.2511.02986. arXiv: 2511.02986 [stat].

[77] Alexander Partin et al. Benchmarking Community Drug Response Prediction Models: Datasets, Models, Tools, and Metrics for Cross-Dataset Generalization Analysis. Mar. 2025. DOI: 10.48550/arXiv.2503.14356. arXiv: 2503.14356 [cs]. (Visited on 04/14/2026).

[78] Stefan Peidli et al. “scPerturb: Harmonized Single-Cell Perturbation Data”. In: Nature Methods 21.3 (Mar. 2024), pp. 531–540. ISSN: 1548-7091, 1548-7105. DOI: 10.1038/s41592-023-02144-y.

[79] Lawrence Phillips et al. SynthPert: Enhancing LLM Biological Reasoning via Synthetic Reasoning Traces for Cellular Perturbation Prediction. Sept. 2025. DOI: 10.48550/arXiv.2509.25346. arXiv: 2509.25346 [cs]. (Visited on 10/28/2025).

[80] Yinhua Piao et al. “Learning Adaptive Perturbation-Conditioned Contexts for Robust Transcriptional Response Prediction”. In: arXiv arXiv:2602.18885 (Feb. 2026). DOI: 10.48550/arXiv.2602.18885. arXiv: 2602.18885 [cs].

[81] Zoe Piran et al. “Disentanglement of Single-Cell Data with Biolord”. In: Nature Biotechnology 42.11 (Nov. 2024), pp. 1678–1683. ISSN: 1087-0156, 1546-1696. DOI: 10.1038/s41587-023-02079-x. (Visited on 12/07/2024).

[82] Jean Radig et al. “scArchon: A Scalable Benchmarking Framework for Assessing Single-Cell Perturbation Models”. In: Genome Biology 27.1 (May. 2026), p. 162. ISSN: 1474-760X. DOI: 10.1186/s13059-026-04104-z. (Visited on 05/18/2026).

[83] Eliot Ragueneau et al. “The Reactome Knowledgebase 2026”. In: Nucleic Acids Research 54.D1 (Jan. 2026), pp. D673–D681. ISSN: 1362-4962. DOI: 10.1093/nar/gkaf1223. (Visited on 06/11/2026).

[84] Felix Reisen et al. “Benchmarking of Multivariate Similarity Measures for High-Content Screening Fingerprints in Phenotypic Drug Discovery”. In: SLAS Discovery. Special Issue: Phenotypic Drug Discovery (Part 1 of 2) 18.10 (Dec. 2013), pp. 1284–1297. ISSN: 2472-5552. DOI: 10.1177/1087057113501390.

[85] Joseph M. Replogle et al. “Mapping Information-Rich Genotype-Phenotype Landscapes with Genome-Scale Perturb-Seq”. In: Cell 185.14 (July. 2022), 2559–2575.e28. ISSN: 00928674. DOI: 10.1016/j.cell.2022.05.013.

[86] Eve Richardson et al. “The Receiver Operating Characteristic Curve Accurately Assesses Imbalanced Datasets”. In: Patterns 5.6 (June. 2024), p. 100994. ISSN: 2666-3899. DOI: 10.1016/j.patter.2024.100994.

[87] Syed Asad Rizvi et al. “Scaling Large Language Models for Next-Generation Single-Cell Analysis”. In: bioRxiv (Apr. 2025), p. 2025.04.14.648850. DOI: 10.1101/2025.04.14.648850.

[88] Martin Rohbeck et al. “Bicycle: Intervention-Based Causal Discovery with Cycles”. In: Proceedings of Machine Learning Research 236 (2024), pp. 209–242.

[89] Jennifer E. Rood, Anna Hupalowska, and Aviv Regev. “Toward a Foundation Model of Causal Cell and Tissue Biology with a Perturbation Cell and Tissue Atlas”. In: Cell 187.17 (Aug. 2024), pp. 4520–4545. ISSN: 00928674. DOI: 10.1016/j.cell.2024.07.035. (Visited on 12/07/2024).

[90] Yusuf Roohani, Kexin Huang, and Jure Leskovec. “Predicting Transcriptional Outcomes of Novel Multigene Perturbations with GEARS”. In: Nature Biotechnology 42.6 (June. 2024), pp. 927–935. ISSN: 1087-0156, 1546-1696. DOI: 10.1038/s41587-023-01905-6.

[91] Yusuf H. Roohani et al. “Virtual Cell Challenge: Toward a Turing Test for the Virtual Cell”. In: Cell 188.13 (June. 2025), pp. 3370–3374. ISSN: 0092-8674, 1097-4172. DOI: 10.1016/j.cell.2025.06.008.

[92] Jiafa Ruan et al. Beyond Independent Genes: Learning Module-Inductive Representations for Gene Perturbation Prediction. Feb. 2026. DOI: 10.48550/arXiv.2602.04901. arXiv: 2602.04901 [q-bio]. (Visited on 04/28/2026).

[93] Andrea Rubbi et al. “A Standardized Framework For Evaluating Gene Expression Generative Models”. In: arXiv (2026). DOI: 10.48550/arXiv.2603.11244. URL: https://arxiv.org/abs/2603.11244.

[94] Reuben A. Saunders et al. “Perturb-Multimodal: A Platform for Pooled Genetic Screens with Imaging and Sequencing in Intact Mammalian Tissue”. In: Cell 188.17 (Aug. 2025), 4790–4809.e22. ISSN: 0092-8674. DOI: 10.1016/j.cell.2025.05.022. (Visited on 04/29/2026).

[95] Philipp Sven Lars Schäfer et al. “Integrating Single-Cell Multi-Omics and Prior Biological Knowledge for a Functional Characterization of the Immune System”. In: Nature Immunology 25.3 (Mar. 2024), pp. 405–417. ISSN: 1529-2908, 1529-2916. DOI:10.1038/s41590-024-01768-2.

[96] Steven Shave et al. Leak Proof CMap; a Framework for Training and Evaluation of Cell Line Agnostic L1000 Similarity Methods. Apr. 2024. DOI: 10.48550/arXiv.2404.18960. arXiv: 2404.18960 [q-bio]. (Visited on 04/14/2026).

[97] Kaden M. Southard et al. “Comprehensive Transcription Factor Perturbations Recapitulate Fibroblast Transcriptional States”. In: Nature Genetics 57.9 (Sept. 2025), pp. 2323–2334. ISSN: 1546-1718. DOI: 10.1038/s41588-025-02284-1. (Visited on 12/05/2025).

[98] Jan L. Sprengel and Britta Velten. Comparison of Interventional Causal Structure Learning Algorithms for Gene Regulatory Network Inference. Dec. 2025. DOI: 10.64898/2025.12.05.692565.

[99] George Stein et al. Exposing Flaws of Generative Model Evaluation Metrics and Their Unfair Treatment of Diffusion Models. Oct. 2023. DOI: 10.48550/arXiv.2306.04675. arXiv: 2306.04675 [cs]. (Visited on 03/31/2026).

[100] Yangqi Su et al. “pertTF: Context-Aware AI Modeling for Genome-Scale and Cross-System Perturbation Prediction”. In: bioRxiv (Mar. 2026), p. 2026.03.12.711379. ISSN: 2692-8205. DOI: 10.64898/2026.03.12.711379.

[101] Junwei Sun, Ouyang Zhu, and Yiqun T. Chen. “Rethinking Perturbation Prediction Baselines”. In: ICLR 2026 Workshop on Machine Learning for Genomics Explorations. Mar. 2026. (Visited on 03/05/2026).

[102] Artur Szałata et al. “A Benchmark for Prediction of Transcriptomic Responses to Chemical Perturbations across Cell Types”. In: Advances in Neural Information Processing Systems 37 (Dec. 2024), pp. 20566–20616. DOI: 10.52202/079017-0650. URL: https://proceedings.neurips.cc/paper_files/paper/2024/hash/24c4d51f3ef48dd2dbab78243ecb26a1-Abstract-Datasets_and_Benchmarks_Track.html.

[103] Juan M. Vaquerizas et al. “A Census of Human Transcription Factors: Function, Expression and Evolution”. In: Nature Reviews Genetics 10.4 (Apr. 2009), pp. 252–263. ISSN: 1471-0056, 1471-0064. DOI: 10.1038/nrg2538. (Visited on 12/07/2024).

[104] Ramon Viñas Torné et al. “Systema: A Framework for Evaluating Genetic Perturbation Response Prediction beyond Systematic Variation”. In: Nature Biotechnology (Aug. 2025), pp. 1–10. ISSN: 1546-1696. DOI: 10.1038/s41587-025-02777-8.

[105] Michael Vollenweider and Peter Bühlmann. Signal, Bounds, and Baselines: Principles for Rigorous Evaluation of High-Dimensional Biological Perturbation Prediction. Apr. 2026. DOI: 10.64898/2026.04.20.719650. (Visited on 04/27/2026).

[106] Chloe Wang et al. “X-Cell: Scaling Causal Perturbation Prediction Across Diverse Cellular Contexts via Diffusion Language Models”. In: bioRxiv (2026). DOI: 10.64898/2026.03.18.712807. URL: https://www.biorxiv.org/content/10.64898/2026.03.18.712807v1.

[107] Jonathan Weed and Francis Bach. “Sharp Asymptotic and Finite-Sample Rates of Convergence of Empirical Measures in Wasserstein Distance”. In: Bernoulli 25.4A (Nov. 2019), pp. 2620–2648. ISSN: 1350-7265. DOI: 10.3150/18-BEJ1065. (Visited on 04/16/2026).

[108] Zhiting Wei et al. “Benchmarking Algorithms for Generalizable Single-Cell Perturbation Response Prediction”. In: Nature Methods (Dec. 2025). scPerturBench, https://github.com/bm2-lab/scPerturBench, pp. 1–14. ISSN: 1548-7105. DOI: 10.1038/s41592-025-02980-0.

[109] Frederik Wenkel et al. “TxPert: Leveraging Biochemical Relationships for Out-of-Distribution Transcriptomic Perturbation Prediction”. In: arXiv arXiv: 2505.14919 (May. 2025). DOI: 10.48550/arXiv.2505.14919. arXiv: 2505.14919.

[110] A. Wenteler et al. “PertEval-scFM: Benchmarking Single-Cell Foundation Models for Perturbation Effect Prediction”. In: bioRxiv (Oct. 2024). DOI: 10.1101/2024.10.02.616248.

[111] Hans-Hermann Wessels et al. “Efficient Combinatorial Targeting of RNA Transcripts in Single Cells with Cas13 RNA Perturb-Seq”. In: Nature Methods 20.1 (Jan. 2023), pp. 86–94. ISSN: 1548-7091, 1548-7105. DOI: 10.1038/s41592-022-01705-x.

[112] Daniel R Wong, Abby S Hill, and Rob Moccia. “Simple Controls Exceed Best Deep Learning Al-gorithms and Reveal Foundation Model Effectiveness for Predicting Genetic Perturbations”. In: Bioinformatics 41.6 (June. 2025), btaf317. ISSN: 1367-4811. DOI: 10.1093/bioinformatics/btaf317.

[113] Menghua Wu et al. Contextualizing Biological Perturbation Experiments through Language. Feb. 2025. DOI: 10.48550/arXiv.2502.21290. arXiv: 2502.21290 [cs]. (Visited on 09/25/2025).

[114] Yan Wu et al. “PerturBench: Benchmarking Machine Learning Models for Cellular Perturbation Analysis”. In: NeurIPS 2024 Workshop on AI for New Drug Modalities. NeurIPS 2024 Workshop on AI for New Drug Modalities. Dec. 2024. URL: https://arxiv.org/abs/2408.10609.

[115] Yuanfang Xiang and Lun Ai. “Adaptive Data-Knowledge Alignment in Genetic Perturbation Prediction”. In: The Fourteenth International Conference on Learning Representations. Oct. 2025. (Visited on 04/22/2026).

[116] Jake Yeung et al. Joint Analysis of Multiply Perturbed Cells Improves Statistical Power and Cost Efficiency in Perturb-Seq. July 2026. DOI: 10.64898/2026.07.10.737863. (Visited on 07/13/2026).

[117] Hengshi Yu et al. “PerturbNet Predicts Single-Cell Responses to Unseen Chemical and Genetic Perturbations”. In: Molecular Systems Biology 21.8 (Aug. 2025), pp. 960–982. ISSN: 1744-4292. DOI: 10.1038/s44320-025-00131-3.

[118] Luke Zappia et al. “Feature Selection Methods Affect the Performance of scRNA-Seq Data Integration and Querying”. In: Nature Methods (Mar. 2025), pp. 1–11. ISSN: 1548-7105. DOI: 10.1038/s41592-025-02624-3.

[119] Jesse Zhang et al. “Tahoe-100M: A Giga-Scale Single-Cell Perturbation Atlas for Context-Dependent Gene Function and Cellular Modeling”. In: bioRxiv (Feb. 2025), p. 2025.02.20.639398. DOI: 10.1101/2025.02.20.639398.

[120] Jiaqi Zhang et al. “Active Learning for Optimal Intervention Design in Causal Models”. In: Nature Machine Intelligence 5.10 (Oct. 2023), pp. 1066–1075. ISSN: 2522-5839. DOI: 10.1038/s42256-023-00719-0. (Visited on 02/13/2025).

[121] Hongxu Zhu et al. “AUPRC: A Metric for Evaluating the Performance of in-Silico Perturbation Methods in Identifying Differentially Expressed Genes”. In: Briefings in Bioinformatics 26.5 (Sept. 2025), bbaf426. ISSN: 1477-4054. DOI: 10.1093/bib/bbaf426.

[122] Ronghui Zhu et al. “Genome-Scale Perturb-Seq in Primary Human CD4+ T Cells Maps Context-Specific Regulators of T Cell Programs and Human Immune Traits”. In: bioRxiv (Dec. 2025), p. 2025.12.23.696273. ISSN: 2692-8205. DOI: 10.64898/2025.12.23.696273.

## Supplementary References

[2] Abhinav K. Adduri et al. “Predicting Cellular Responses to Perturbation across Diverse Contexts with State”. In: bioRxiv (July 2025), p. 2025.06.26.661135. ISSN: 2692-8205. DOI: 10.1101/2025.06.26.661135.

[3] Constantin Ahlmann-Eltze and Wolfgang Huber. “Comparison of Transformations for Single-Cell RNA-Seq Data”. In: Nature Methods 20.5 (May 2023), pp. 665–672. ISSN: 1548-7105. DOI: 10.1038/s41592-023-01814-1. (Visited on 01/12/2025).

[4] Timothy Barry et al. “Robust Differential Expression Testing for Single-Cell CRISPR Screens at Low Multiplicity of Infection”. In: Genome Biology 25.1 (May 2024), p. 124. ISSN: 1474-760X. DOI: 10.1186/s13059-024-03254-2.

[5] Timothy Barry et al. “SCEPTRE Improves Calibration and Sensitivity in Single-Cell CRISPR Screen Analysis”. In: Genome Biology 22.1 (Dec. 2021), p. 344. ISSN: 1474-760X. DOI: 10.1186/s13059-021-02545-2.

[6] Michael D. Bereket and Jure Leskovec. Are Current AI Virtual Cell Models Useful for Scientific Discovery? Apr. 2026. DOI: 10.64898/2026.04.23.719015. (Visited on 04/27/2026).

[7] Sebastian Bischoff et al. “A Practical Guide to Sample-Based Statistical Distances for Evaluating Generative Models in Science”. In: arXiv arXiv: 2403.12636 (Oct. 2024). DOI: 10.48550/arXiv.2403.12636. arXiv: 2403.12636.

[8] Charlotte Bunne, Andreas Krause, and Marco Cuturi. “Supervised Training of Conditional Monge Maps”. In: arXiv arXiv:2206.14262 (Mar. 2023). DOI: 10.48550/arXiv.2206.14262. arXiv: 2206.14262 [cs].

[9] Yingying Cao et al. “UMI or Not UMI, That Is the Question for scRNA-Seq Zero-Inflation”. In: Nature Biotechnology 39.2 (Feb. 2021), pp. 158–159. ISSN: 1087-0156, 1546-1696. DOI: 10.1038/s41587-020-00810-6. (Visited on 12/07/2024).

[10] Yunshun Chen et al. “edgeR v4: Powerful Differential Analysis of Sequencing Data with Expanded Functionality and Improved Support for Small Counts and Larger Datasets”. In: Nucleic Acids Research 53.2 (Jan. 2025), gkaf018. ISSN: 1362-4962. DOI: 10.1093/nar/gkaf018.

[11] Saket Choudhary and Rahul Satija. “Comparison and Evaluation of Statistical Error Models for scRNA-Seq”. In: Genome Biology 23.1 (Jan. 2022), p. 27. ISSN: 1474-760X. DOI: 10.1186/s13059-021-02584-9. (Visited on 12/07/2024).

[12] Oliver P. Christensen et al. “Causal Effect Estimation from Trans-Regulatory Single-Cell CRISPR Screens”. In: Cell Genomics 6.6 (June 2026). ISSN: 2666-979X. DOI: 10.1016/j.xgen.2026.101251. (Visited on 06/11/2026).

[13] Elijah Cole et al. “Foundation Models Improve Perturbation Response Prediction”. In: bioRxiv (Feb. 2026), p. 2026.02.18.706454. ISSN: 2692-8205. DOI: 10.64898/2026.02.18.706454.

[14] Haotian Cui et al. “scGPT: Toward Building a Foundation Model for Single-Cell Multi-Omics Using Generative AI”. In: Nature Methods 21.8 (Aug. 2024), pp. 1470–1480. ISSN: 1548-7105. DOI: 10.1038/s41592-024-02201-0.

[15] Marco Cuturi. “Sinkhorn Distances: Lightspeed Computation of Optimal Transportation Distances”. In: arXiv arXiv: 1306.0895 (June 2013). DOI: 10.48550/arXiv.1306.0895. arXiv: 1306.0895.

[16] Atray Dixit et al. “Perturb-Seq: Dissecting Molecular Circuits with Scalable Single-Cell RNA Profiling of Pooled Genetic Screens”. In: Cell 167.7 (Dec. 2016), 1853–1866.e17. ISSN: 00928674. DOI: 10.1016/j.cell.2016.11.038.

[17] Mingze Dong et al. “Stack: In-Context Learning of Single-Cell Biology”. In: bioRxiv (2026). DOI: 10.64898/2026.01.09.698608. URL: https://connect.biorxiv.org/relate/content/224/channel.

[18] Eric and Wendy Schmidt Center at the Broad Institute and Diabetes Initiative at the Broad Institute of MIT and Harvard. Obesity Machine Learning Competition: Tackling Metabolic Diseases. Cell Perturbation Prediction Challenge. 2025. URL: https://www.broadinstitute.org/videos/obesity-machine-learning-competition-tackling-metabolic-diseases-overview.

[19] Claudia Feng et al. “A Genome-Scale Single-Cell CRISPRi Map of Trans Gene Regulation across Human Pluripotent Stem Cell Lines”. In: Cell Genomics 6.2 (Feb. 2026). ISSN: 2666-979X. DOI: 10.1016/j.xgen.2025.101076. (Visited on 04/13/2026).

[20] Jean Feydy et al. “Interpolating between Optimal Transport and MMD Using Sinkhorn Divergences”. In: Proceedings of the Twenty-Second International Conference on Artificial Intelligence and Statistics. PMLR, Apr. 2019, pp. 2681–2690. URL: https://proceedings.mlr.press/v89/feydy19a.html.

[21] Greg Finak et al. “MAST: A Flexible Statistical Framework for Assessing Transcriptional Changes and Characterizing Heterogeneity in Single-Cell RNA Sequencing Data”. In: Genome Biology 16.1 (2015), p. 278. ISSN: 1474-7596. DOI: 10.1186/s13059-015-0844-5.

[22] Chris J. Frangieh et al. “Multimodal Pooled Perturb-CITE-Seq Screens in Patient Models Define Mechanisms of Cancer Immune Evasion”. In: Nature Genetics 53.3 (Mar. 2021), pp. 332–341. ISSN: 1546-1718. DOI: 10.1038/s41588-021-00779-1.

[23] Aude Genevay, Gabriel Peyre, and Marco Cuturi. “Learning Generative Models with Sinkhorn Divergences”. In: Proceedings of the Twenty-First International Conference on Artificial Intelligence and Statistics. PMLR, Mar. 2018, pp. 1608–1617. URL: https://proceedings.mlr.press/v84/genevay18a.html.

[24] Ryan M. J. Genga et al. “Single-Cell RNA-Sequencing-Based CRISPRi Screening Resolves Molecular Drivers of Early Human Endoderm Development”. In: Cell Reports 27.3 (Apr. 2019), 708–718.e10. ISSN: 2211-1247. DOI: 10.1016/j.celrep.2019.03.076.

[25] Arthur Gretton et al. “A Kernel Two-Sample Test”. In: Journal of Machine Learning Research 13.25 (2012), pp. 723–773. ISSN: 1533-7928. (Visited on 07/27/2025).

[26] Christoph Hafemeister and Rahul Satija. “Normalization and Variance Stabilization of Single-Cell RNA-Seq Data Using Regularized Negative Binomial Regression”. In: Genome Biology 20.1 (Dec. 2019), p. 296. ISSN: 1474-760X. DOI: 10.1186/s13059-019-1874-1. (Visited on 12/07/2024).

[27] Lukas Heumos et al. “Best Practices for Single-Cell Analysis across Modalities”. In: Nature Reviews Genetics 24.8 (Aug. 2023), pp. 550–572. ISSN: 1471-0056, 1471-0064. DOI: 10.1038/s41576-023-00586-w.

[28] Lukas Heumos et al. “Pertpy: An End-to-End Framework for Perturbation Analysis”. In: Nature Methods 23.2 (Feb. 2026), pp. 350–359. ISSN: 1548-7105. DOI: 10.1038/s41592-025-02909-7.

[29] Martin Heusel et al. “GANs Trained by a Two Time-Scale Update Rule Converge to a Local Nash Equilibrium”. In: Advances in Neural Information Processing Systems (NeurIPS). arXiv:1706.08500. Jan. 2018. DOI: 10.48550/arXiv.1706.08500. eprint: 1706.08500.

[30] Trey Ideker, Janusz Dutkowski, and Leroy Hood. “Boosting Signal-to-Noise in Complex Biology: Prior Knowledge Is Power”. In: Cell 144.6 (Mar. 2011), pp. 860–863. ISSN: 0092-8674. DOI: 10.1016/j.cell.2011.03.007. (Visited on 05/04/2026).

[31] Yuge Ji et al. “Optimal Distance Metrics for Single-Cell RNA-Seq Populations”. In: bioRxiv (Dec. 2023), p. 2023.12.26.572833. DOI: 10.1101/2023.12.26.572833.

[32] Qun Jiang et al. “scPRAM Accurately Predicts Single-Cell Gene Expression Perturbation Response Based on Attention Mechanism”. In: Bioinformatics 40.5 (May 2024), btae265. ISSN: 1367-4811. DOI: 10.1093/bioinformatics/btae265.

[33] Ruochen Jiang et al. “Statistics or Biology: The Zero-Inflation Controversy about scRNA-Seq Data”. In: Genome Biology 23.1 (Jan. 2022), p. 31. ISSN: 1474-760X. DOI: 10.1186/s13059-022-02601-5. (Visited on 12/07/2024).

[34] Min Cheol Kim et al. “Method of Moments Framework for Differential Expression Analysis of Single-Cell RNA Sequencing Data”. In: Cell 187.22 (Oct. 2024), 6393–6410.e16. ISSN: 0092-8674, 1097-4172. DOI: 10.1016/j.cell.2024.09.044.

[35] Dominik Klein et al. “CellFlow Enables Generative Single-Cell Phenotype Modeling with Flow Matching”. In: bioRxiv (Apr. 2025), p. 2025.04.11.648220. DOI: 10.1101/2025.04.11.648220.

[36] Matthew A. Lalli et al. “High-Throughput Single-Cell Functional Elucidation of Neurodevelopmental Disease–Associated Genes Reveals Convergent Mechanisms Altering Neuronal Differentiation”. In: Genome Research 30.9 (Jan. 2020), pp. 1317–1331. ISSN: 1088-9051, 1549-5469. DOI: 10.1101/gr.262295.120.

[37] Jan Lause, Philipp Berens, and Dmitry Kobak. “Analytic Pearson Residuals for Normalization of Single-Cell RNA-Seq UMI Data”. In: Genome Biology 22.1 (2021), p. 258. ISSN: 1474-7596. DOI: 10.1186/s13059-021-02451-7.

[38] Nicolo’ Lazzaro et al. “Signature Distance: Generalizing Energy Statistics”. In: bioRxiv (Mar. 2026), p. 2024.10.23.619602. ISSN: 2692-8205. DOI: 10.1101/2024.10.23.619602.

[39] Hechen Li et al. “scMultiSim: Simulation of Single-Cell Multi-Omics and Spatial Data Guided by Gene Regulatory Networks and Cell–Cell Interactions”. In: Nature Methods 22.5 (May 2025), pp. 982–993. ISSN: 1548-7105. DOI: 10.1038/s41592-025-02651-0. (Visited on 06/05/2026).

[40] Russell Littman et al. “Gene-Embedding-Based Prediction and Functional Evaluation of Perturbation Expression Responses with PRESAGE”. In: bioRxiv (June 2025), p. 2025.06.03.657653. DOI: 10.1101/2025.06.03.657653.

[41] Romain Lopez et al. “Deep Generative Modeling for Single-Cell Transcriptomics”. In: Nature Methods 15.12 (Dec. 2018), pp. 1053–1058. ISSN: 1548-7091, 1548-7105. DOI: 10.1038/s41592-018-0229-2.

[42] David Lopez-Paz and Maxime Oquab. “Revisiting Classifier Two-Sample Tests”. In: International Conference on Learning Representations. Feb. 2017. URL: https://openreview.net/forum?id=SJkXfE5xx.

[43] Michael I Love, Wolfgang Huber, and Simon Anders. “Moderated Estimation of Fold Change and Dispersion for RNA-Seq Data with DESeq2”. In: Genome Biology 15.12 (Dec. 2014), p. 550. ISSN: 1474-760X. DOI: 10.1186/s13059-014-0550-8.

[44] Aaron T. L. Lun, Karsten Bach, and John C. Marioni. “Pooling across Cells to Normalize Single-Cell RNA Sequencing Data with Many Zero Counts”. In: Genome Biology 17.1 (2016), p. 75. ISSN: 1474-7596. DOI: 10.1186/s13059-016-0947-7.

[45] Gabriel M. Mejia et al. “Diversity by Design: Addressing Mode Collapse Improves scRNA-seq Perturbation Modeling on Well-Calibrated Metrics”. In: arXiv (2025). DOI: 10.48550/arXiv.2506.22641. arXiv: 2506.22641. URL: https://arxiv.org/abs/2506.22641.

[46] Henry E. Miller et al. “Deep Learning-Based Genetic Perturbation Models Do Outperform Uninformative Baselines on Well-Calibrated Metrics”. In: bioRxiv (Oct. 2025), p. 2025.10.20.683304. ISSN: 2692-8205. DOI: 10.1101/2025.10.20.683304.

[47] Eleni P. Mimitou et al. “Multiplexed Detection of Proteins, Transcriptomes, Clonotypes and CRISPR Perturbations in Single Cells”. In: Nature Methods 16.5 (May 2019), pp. 409–412. ISSN: 1548-7105. DOI: 10.1038/s41592-019-0392-0.

[48] Myllia. Echoes of Silenced Genes: A Cell Challenge. https://www.kaggle.com/competitions/echoes-of-silenced-genes. Kaggle competition page. Accessed: 2026-04-19. 2026.

[49] Ajay Nadig et al. “Transcriptome-Wide Analysis of Differential Expression in Perturbation Atlases”. In: Nature Genetics 57.5 (May 2025), pp. 1228–1237. ISSN: 1546-1718. DOI: 10.1038/s41588-025-02169-3.

[50] Alessandro Palma et al. “Multi-Modal and Multi-Attribute Generation of Single Cells with CFGen”. In: The Thirteenth International Conference on Learning Representations. Oct. 2025. URL: https://openreview.net/forum?id=3MnMGLctKb.

[51] Efthymia Papalexi et al. “Characterizing the Molecular Regulation of Inhibitory Immune Checkpoints with Multimodal Single-Cell Screens”. In: Nature Genetics 53.3 (Mar. 2021), pp. 322–331. ISSN: 1546-1718. DOI: 10.1038/s41588-021-00778-2.

[52] Stefan Peidli et al. “scPerturb: Harmonized Single-Cell Perturbation Data”. In: Nature Methods 21.3 (Mar. 2024), pp. 531–540. ISSN: 1548-7091, 1548-7105. DOI: 10.1038/s41592-023-02144-y.

[53] Jean Radig et al. “scArchon: A Scalable Benchmarking Framework for Assessing Single-Cell Perturbation Models”. In: Genome Biology 27.1 (May 2026), p. 162. ISSN: 1474-760X. DOI: 10.1186/s13059-026-04104-z. (Visited on 05/18/2026).

[54] Joseph M. Replogle et al. “Combinatorial Single-Cell CRISPR Screens by Direct Guide RNA Capture and Targeted Sequencing”. In: Nature Biotechnology 38.8 (Aug. 2020), pp. 954–961. ISSN: 1546-1696. DOI: 10.1038/s41587-020-0470-y.

[55] Matthew E. Ritchie et al. “Limma Powers Differential Expression Analyses for RNA-Sequencing and Microarray Studies”. In: Nucleic Acids Research 43.7 (2015), e47–e47. ISSN: 0305-1048. DOI: 10.1093/nar/gkv007. (Visited on 04/19/2026).

[56] Syed Asad Rizvi et al. “Scaling Large Language Models for Next-Generation Single-Cell Analysis”. In: bioRxiv (Apr. 2025), p. 2025.04.14.648850. DOI: 10.1101/2025.04.14.648850.

[57] Maria L. Rizzo and Gábor J. Székely. “Energy Distance”. In: WIREs Computational Statistics 8.1 (2016), pp. 27–38. ISSN: 1939-0068. DOI: 10.1002/wics.1375.

[58] Yusuf Roohani, Kexin Huang, and Jure Leskovec. “Predicting Transcriptional Outcomes of Novel Multigene Perturbations with GEARS”. In: Nature Biotechnology 42.6 (June 2024), pp. 927–935. ISSN: 1087-0156, 1546-1696. DOI: 10.1038/s41587-023-01905-6.

[59] Yusuf H. Roohani et al. “Virtual Cell Challenge: Toward a Turing Test for the Virtual Cell”. In: Cell 188.13 (June 2025), pp. 3370–3374. ISSN: 0092-8674, 1097-4172. DOI: 10.1016/j.cell.2025.06.008.

[60] Philipp Sven Lars Schäfer et al. “Integrating Single-Cell Multi-Omics and Prior Biological Knowledge for a Functional Characterization of the Immune System”. In: Nature Immunology 25.3 (Mar. 2024), pp. 405–417. ISSN: 1529-2908, 1529-2916. DOI: 10.1038/s41590-024-01768-2.

[61] Daniel Schraivogel et al. “Targeted Perturb-Seq Enables Genome-Scale Genetic Screens in Single Cells”. In: Nature Methods 17.6 (June 2020), pp. 629–635. ISSN: 1548-7091, 1548-7105. DOI: 10.1038/s41592-020-0837-5.

[62] Dongyuan Song et al. “scDesign3 Generates Realistic in Silico Data for Multimodal Single-Cell and Spatial Omics”. In: Nature Biotechnology 42.2 (Feb. 2024), pp. 247–252. ISSN: 1087-0156, 1546-1696. DOI: 10.1038/s41587-023-01772-1. (Visited on 12/07/2024).

[63] Jordan W. Squair et al. “Confronting False Discoveries in Single-Cell Differential Expression”. In: Nature Communications 12.1 (Sept. 2021), p. 5692. ISSN: 2041-1723. DOI: 10.1038/s41467-021-25960-2.

[64] Valentine Svensson. “Droplet scRNA-Seq Is Not Zero-Inflated”. In: Nature Biotechnology 38.2 (2020), pp. 147–150. ISSN: 1087-0156. DOI: 10.1038/s41587-019-0379-5.

[65] Christian Szegedy et al. “Rethinking the Inception Architecture for Computer Vision”. In: 2016 IEEE Conference on Computer Vision and Pattern Recognition (CVPR). Las Vegas, NV, USA: IEEE, June 2016, pp. 2818–2826. ISBN: 978-1-4673-8851-1. DOI: 10.1109/CVPR.2016.308.

[66] F. William Townes et al. “Feature Selection and Dimension Reduction for Single-Cell RNA-Seq Based on a Multinomial Model”. In: Genome Biology 20.1 (Dec. 2019), p. 295. ISSN: 1474-760X. DOI: 10.1186/s13059-019-1861-6. (Visited on 12/07/2024).

[67] Cole Trapnell. “Revealing Gene Function with Statistical Inference at Single-Cell Resolution”. In: Nature Reviews Genetics 25.9 (Sept. 2024), pp. 623–638. ISSN: 1471-0064. DOI: 10.1038/s41576-024-00750-w.

[68] Oana Ursu et al. “Massively Parallel Phenotyping of Coding Variants in Cancer with PerturbSeq”. In: Nature Biotechnology 40.6 (June 2022), pp. 896–905. ISSN: 1546-1696. DOI: 10.1038/s41587-021-01160-7.

[69] Michael Vollenweider and Peter Bühlmann. Signal, Bounds, and Baselines: Principles for Rigorous Evaluation of High-Dimensional Biological Perturbation Prediction. Apr. 2026. DOI: 10.64898/2026.04.20.719650. (Visited on 04/27/2026).

[70] Chloe Wang et al. “X-Cell: Scaling Causal Perturbation Prediction Across Diverse Cellular Contexts via Diffusion Language Models”. In: bioRxiv (2026). DOI: 10.64898/2026.03.18.712807. URL: https://www.biorxiv.org/content/10.64898/2026.03.18.712807v1.

[71] Zhiting Wei et al. “Benchmarking Algorithms for Generalizable Single-Cell Perturbation Response Prediction”. In: Nature Methods (Dec. 2025). scPerturBench, https://github.com/bm2-lab/scPerturBench, pp. 1–14. ISSN: 1548-7105. DOI: 10.1038/s41592-025-02980-0.

[72] Hans-Hermann Wessels et al. “Efficient Combinatorial Targeting of RNA Transcripts in Single Cells with Cas13 RNA Perturb-Seq”. In: Nature Methods 20.1 (Jan. 2023), pp. 86–94. ISSN: 1548-7091, 1548-7105. DOI: 10.1038/s41592-022-01705-x.

[73] Yan Wu et al. “PerturBench: Benchmarking Machine Learning Models for Cellular Perturbation Analysis”. In: NeurIPS 2024 Workshop on AI for New Drug Modalities. NeurIPS 2024 Workshop on AI for New Drug Modalities. Dec. 2024. URL: https://arxiv.org/abs/2408.10609.

[74] Lin Yang et al. “scMAGeCK Links Genotypes with Multiple Phenotypes in Single-Cell CRISPR Screens”. In: Genome Biology 21.1 (Dec. 2020), p. 19. ISSN: 1474-760X. DOI: 10.1186/s13059-020-1928-4.

[75] Luke Zappia, Belinda Phipson, and Alicia Oshlack. “Splatter: Simulation of Single-Cell RNA Sequencing Data”. In: Genome Biology 18.1 (Sept. 2017), p. 174. ISSN: 1474-760X. DOI: 10.1186/s13059-017-1305-0. (Visited on 06/05/2026).

[76] Luke Zappia et al. “Feature Selection Methods Affect the Performance of scRNA-Seq Data Integration and Querying”. In: Nature Methods (Mar. 2025), pp. 1–11. ISSN: 1548-7105. DOI: 10.1038/s41592-025-02624-3.

[77] Xian Zhang and Michael Boutros. “A Novel Phenotypic Dissimilarity Method for Image-Based High-Throughput Screens”. In: BMC bioinformatics 14.1 (Nov. 2013), p. 336. ISSN: 1471-2105. DOI: 10.1186/1471-2105-14-336.

[78] Hongxu Zhu et al. “AUPRC: A Metric for Evaluating the Performance of in-Silico Perturbation Methods in Identifying Differentially Expressed Genes”. In: Briefings in Bioinformatics 26.5 (Sept. 2025), bbaf426. ISSN: 1477-4054. DOI: 10.1093/bib/bbaf426.

